# Evolutionary history and recurrent host adaptation in ancient *Salmonella enterica*

**DOI:** 10.1101/2025.09.18.676881

**Authors:** Iseult Jackson, Gunnar U. Neumann, Anna-Theresa Mayr, Aida Andrades Valtueña, Alina N. Hiss, Lyazzat Musralina, Kurt W. Alt, Arman Beisenov, Lorenc Bejko, Natalia Berezina, Margit Berner, Aparajita Bhattacharya, Michaela Binder, Lia Bitadze, Mária Bondár, Marina Bretos, Igor Bruyako, Ivan Bugarski, Alexandra Buzhilova, Alexandra Charami, Kevin Daly, Jocelyne Desideri, Marta Díaz-Zorita Bonilla, Leyla Djansugurova, Déborah Rosselet-Christ, Paula Doumani Dupuy, Sabine Eggers, Sturla Ellingvåg, Michal Ernée, Mirosław Furmanek, Anja Furtwängler, Dániel Gerber, Belen Gimeno Martínez, Brigitte Haas-Gebhard, Agata Hałuszko, Michaela Harbeck, Daniela Heilmann, Vujadin Ivanišević, Xiaowen Jia, Marina Karapetian, Marcel Keller, Elmira Khussainova, Yuri F. Kiryushin, Rüdiger Krause, Mária Krošláková, Thomas Kubenz, Stephanie Larson, Maria A. Liston, Hellen Mager, Igor Manzura, Michael McCormick, Balázs G. Mende, Ilya V. Merts, Victor K. Merts, Nataša Miladinović-Radmilović, Arnold Muhl, Erdene Myagmar, Nicole Nicklisch, Jörg Orschiedt, Mathieu Ott, Doris Pany-Kucera, Luka Papac, Aleksandra Papazovska, María Victoria Pastor, Jaroslav Peška, Piroska Rácz, Claude Raynaud, Roberto Risch, Matej Ruttkay, Zainolla Samashev, David R. Scahill, Svetlana Sharapova, Elena Sintes Olives, Eirini Skourtanioti, Adéla Sobotkova, Astrid Stobbe, Tatjana Stojoska Vidovska, Eliza Stolarczyk, Anna Szécsényi-Nagy, Nino Tavartkiladze, Alexey A. Tishkin, Solenn Troadec, Emma Usmanova, Diana Vicente, Dragana Vulović, Aidyn Zhuniskhanov, Maria A. Spyrou, Harald Ringbauer, Taylor Hermes, Vanessa Villalba-Mouco, Philipp W. Stockhammer, Thilo Fuchs, Ainash Childebayeva, Christina Warinner, Wolfgang Haak, Johannes Krause, Alexander Herbig

**Affiliations:** Department of Archaeogenetics, Max Planck Institute for Evolutionary Anthropology, Deutscher Pl. 6, Leipzig 04103, Germany; Institute for Pre- and Protohistoric Archaeology and Archaeology of the Roman Provinces, Ludwig Maximilian University Munich, Geschwister-Scholl-Platz 1, Munich 80539, Germany; Center of Natural and Cultural Human History, Danube Private University, Steiner Landstraße 124, Krems, Lower-Austria, 3500, Austria; Begazy-Tasmola Research Center of History and Archeology, Almaty, 50008, Kazakhstan; Institute of Archaeology by A.Kh. Margulan, Dostyk Ave 44, Almaty, 050010, Kazakhstan; Department of Archaeology and Heritage Studies, University of Tirana, Elbasanit 69, Tirana, Tirana, 1010, Albania; Research Institute and Museum of Anthropology, Lomonosov Moscow State University, Mokhovaya 11, Moscow, 125009, Russia; Department of Anthropology, Natural History Museum Vienna, Burging 7, Vienna, 1010, Austria; Novetus GmbH, Belvederegasse 41, Vienna, 1040, Austria; Department of Anthropology, Institute of History and Ethnology, Tbilisi State University, University St. 2, Tbilisi, Georgia; Institute of Archaeology, ELTE Research Centre for the Humanities, Tóth Kálmán u. 4, Budapest, 1097, Hungary; Institute for Research in Environmental Sciences of Aragon (IUCA), University of Zaragoza, Calle de Pedro Cerbuna 12, Zaragoza, 50009, Spain; Odesa Archaeological Museum, Odesa, Ukraine; Institute of Archaeology, Kneza Mihaila 35/IV, Belgrade, 11000, Serbia; Hellenic Ministry of Culture and Sports, Ephorate of Antiquities of Boeotia, Greece; Department of Classics & Ancient Mediterranean Studies, Bucknell University, Dent Drive 1, Lewisburg, Pennsylvania, 17837, USA; Archaeology of Africa & Anthropology Laboratory (ARCAN), University of Geneva, Quai Ernest Ansermet 30, Geneva, Geneva, 1205, Switzerland; Institute for Pre- and Protohistory and Medieval Archaeology, Eberhard Karls Universität Tübingen, Burgsteige 11, Tübingen 72070, Germany; Center of Paleogenetics and Ethnogenomics, Institute of Genetics and Physiology, al-Farabi ave. 93, Almaty, 50060, Kazakhstan; Department of Sociology and Anthropology, Nazarbayev University, Kabanbay Batyr Prospect 53, Almaty, 10000, Kazakhstan; Explico - Historical Research Foundation, Florø, Norway; Department of Prehistory, Institute of Archaeology of the Czech Academy of Sciences, Prague, Letenská 4, Prague, 118 00, Czech Republic; Institute of Archaeology, University of Wrocław, Szewska 48, Wrocław, Silesia, 50-139, Poland; Institute of Archaeogenomics, ELTE Research Centre for the Humanities, Tóth Kálmán u. 4, Budapest, 1097, Hungary; Dirección General de Patrimonio Cultural del Gobierno de Aragón, Zaragoza, 50071, Spain; Department of Early Medieval and Later Archaeology, Bavarian State Archaeological Collection, Lerchenfeldstr. 2, Munich 80538, Germany; Institute of Archeology, Maria Curie-Skłodowska University, M. Curie-Skłodowska Square 4A, Lublin, 20-031, Poland; Archeolodzy.org Foundation, Rynek 21/6, Świdnica, 58-100, Poland; State Collection for Anthropology Munich, SNSB, Karolinenplatz 2a, Munich 8033, Germany; Roman-Germanic Commission, German Archaeological Institute; Department of Historical Sciences, Serbian Academy of Sciences and Arts, Kneza Mihaila 35, Belgrade, 11000, Serbia; Department of Environmental Sciences, University of Basel, Spalenring 145, Basel, 4055, Switzerland; Institute of Genomics, University of Tartu, Riia 23b, Tartu, 51010, Estonia; Department of Archaeology, Ethnography and Museology, Altai State University, Lenin Avenue 61, Barnaul, Altai Krai, 656049, Russia; Institute of Archaeological Sciences, Goethe University Frankfurt am Main, Norbert-Wollheim-Platz 1, Frankfurt 60629, Germany; Institute of Archaeology, Slovak Academy of Sciences, Akademická 2, Nitra, Nitra, 94921, Slovakia; Landesamt für Denkmalpflege und Archäologie Sachsen-Anhalt, Richard-Wagner-Straße 9, Halle/Saale 06114, Germany; Department of Anthropology, University of Waterloo, University Avenue West 200, Waterloo, Ontario, N2L3G1, Canada; Department of History, Archaeology and Museology, National Museum of History of Moldova, 31 August 1989 St 121A, Chişinău, 2012, Moldova; Initiative for the Science of the Human Past at Harvard, Department of History, Harvard University, Quincy St 35, Cambridge, Massachusetts, 02138, USA; Max Planck-Harvard Research Center for the Archaeoscience of the Ancient Mediterranean, Harvard University, Cambridge, Massachusetts, USA; Joint Research Center for Archeological Studies named after A.Kh. Margulan, Toraigyrov University, Lomova 64, Pavlodar, 140008, Kazakhstan; Department of Biochemical Engineering, International Engineering Technological University, al-Farabi ave. 89/21, Almaty, 050060, Kazakhstan; Department of Anthropology and Archaeology, National University of Mongolia, Ikh surguuliin gudamj 1, Ulaanbaatar, 14192, Mongolia; Institut National de Recherches Archéologiques Préventives, Rue Etienne Lenoir 561, Nîmes, 30900, France; UMR5140 - Archéologie des Sociétés Méditerranéennes, Université Paul Valery, Rue du Professeur Henri Serre 2, Montpellier, 34090, France; Archaeological Museum of the Republic of North Macedonia, Kej Dimitar Vlahov, Skopje, 1000, North Macedonia; Independent researcher; Archaeological Centre Olomouc, U Hradiska 42/6, Olomouc, 779 00, Czechia; Unité Mixte de Recherche, UMR 5140, Centre National de la Recherche Scientifique, Montpellier, 34199, France; Department of Prehistory, Universitat Autònoma de Barcelona, Bellaterra, Barcelona, 8193, Spain; State Historical and Cultural Museum-Reserve “Berel”, village Zhambyl, Katon-Karagay, 070906, Kazakhstan; Branch of Institute of Archaeology by A.Kh. Margulan, Republic Ave 24, Astana, 010011, Kazakhstan; Austrian Archaeological Institute, Leoforos Alexandras 26, Athens, 106 83, Greece; Institute of History and Archaeology, Urals Branch of Russian Academy of Sciences, Ekaterinburg, 620108, Russia; Section of History and Archaeology, Institut Menorquí d’Estudis, s’Androna 2, Alaior, Balearic Islands, 7730, Spain; Department of History and Classical Studies, Aarhus University, Jens Christian Skous Vej 5, Aarhus, Midtjylland, 8000, Denmark; Institute for protection of monuments of culture and museum, Boro Shain Street 10, Ohrid, 6000, North Macedonia; Saryarka Archaeological Institute of Karaganda Buketov University, Karaganda, Kazakhstan; Archaeological Institute named after A. Kh. Khalikov, Russia; Kazakh Language and Turkic Studies, Nazarbayev University, Kabanbay Batyr Prospect 53, Astana, Akmola, 10000, Kazakhstan; Department of Anthropology, University of Arkansas, Fayetteville, Arkansas, 72703, USA; Institute of Evolutionary Biology, CSIC-Universitat Pompeu Fabra, Carrer Dr. Aiguader 88, Barcelona, Barcelona, 08003, Spain; Max Planck Harvard Research Center for the Archaeoscience of the Ancient Mediterranean, Max Planck Institute for Evolutionary Anthropology, Deutscher Pl. 6, Leipzig 04103, Germany; Institute of Molecular Pathogenesis, Friedrich-Loeffler-Institut, Naumburger Str. 96a, Jena 07743, Germany; Department of Anthropology, University of Texas at Austin, Speedway 2201, Austin, Texas, 78712, USA; Department of Anthropology, Harvard University, Divinity Ave 11, Cambridge, Massachusetts, 02138, USA; Department of Human Evolutionary Biology, Harvard University, Divinity Ave 11, Cambridge, Massachusetts, 02138, USA; Archaeogenetics Unit, Leibniz Institute for Natural Products Research and Infection Biology, Beutenbergstraße 11, Jena 07745, Germany; Archaeo- and Palaeogenetics, Institute for Archaeological Sciences, Department of Geosciences, Eberhard Karls University of Tübingen, Tübingen 72074, Germany

## Abstract

*Salmonella enterica* subsp. *enterica* is an extremely diverse bacterial pathogen causing frequent infections and foodborne disease among human populations. More than 1500 different bacterial strains (serovars) have been described, many with a wide host range. A small number of serovars are adapted to infect specific hosts: of these, serovars Typhi and Paratyphi A, B, and C cause primate-specific systemic infections (typhoid and paratyphoid fever). Although Paratyphi C is one of the rarest human-specific serovars today, it was once widespread, and all ancient *Salmonella* genomes published to date belong to or are ancestral to this lineage. Here, we present 53 new ancient *Salmonella* genomes spanning Eurasia and dating between 3500 BCE and 1300 CE. This rich genomic dataset allows us to reconstruct the evolutionary history of this pathogen in unprecedented detail. We identify multiple extinct prehistoric lineages that caused infections throughout Eurasia. Multiple lineage replacement events are observed throughout prehistoric and historic times, and Bayesian phylogenetic analysis is used to date and identify host adaptation events within this lineage. We find that host-adapted sublineages Paratyphi C, Choleraesuis, and Typhisuis continued to evolve host specificity independently from each other. We reconstruct signals of convergent host adaptation in the studied lineages and other host-adapted strains by analysing shared pseudogenes and recurrent gene gain and loss events. This analysis demonstrates a role for host interactions as a particular target of selection, highlighting the gradual adaptation of this *S. enterica* lineage to humans that coincides with the intensification of animal husbandry in pastoralist and sedentary farming societies.

## Introduction

*Salmonella enterica* subsp. *enterica* is an extremely diverse pathogen, with over 1500 known serovars capable of infecting the gastrointestinal tracts of various host species, including humans, other mammals, reptiles, fish, and birds^1,2^. Although the majority of these serovars originate from the environment and exhibit a broad host range, some are host-adapted and infect only specific taxa. Infections caused by these host-adapted lineages differ from typical salmonellosis (e.g. foodborne disease), as they lead to extra-intestinal, invasive infections (often referred to as typhoidal infections) with significantly higher mortality rates than non-invasive gastroenteritis. Today, *S. enterica* serovar Typhi is responsible for the majority of invasive infections in humans. However, Paratyphi A and sub-lineages of commonly non-invasive serovars such as Typhimurium and Enteritidis are increasingly associated with invasive salmonellosis cases and deaths in some regions of the world^3,4^.

Evidence from ancient genomes has painted a different picture of invasive salmonellosis in the past. The vast majority of ancient *S. enterica* genomes recovered so far fall into the Para C lineage including modern-day serovars Paratyphi C, Choleraesuis, and Typhisuis^5–11^. Despite this, Paratyphi C infections are rare today^5,12^.

All other ancient *S. enterica* genomes reported to date have either been extremely early host-generalist serovars, or else form a sister lineage to modern-day serovar Birkenhead ^7^, which is rare and host-generalist. This serovar was first described in 1948 in patients from the United Kingdom ^13^. Infections with the same serovar in New South Wales in Australia have been associated with the consumption of unwashed or unpeeled fruit and vegetables, implying an environmental source ^14^.

The Para C lineage seems to have been widespread for over two thousand years, with previously identified prehistoric cases ranging from Switzerland to Western China ^7,8^. Evidence from ancient genomes is particularly important for both reconstructing the evolutionary history of *S. enterica* and in uncovering its impact on human health and disease. Unlike other important human pathogens such as *Mycobacterium tuberculosis* or *Mycobacterium leprae*, it causes acute illness and does not lead to characteristic, recognisable skeletal lesions that allow the identification of past *Salmonella* epidemics^15–17^. Previous ancient DNA studies have identified outbreaks caused by *S. enterica* Paratyphi C in sixteenth century Mexico and fourteenth century Germany ^6,9^.

In addition to identifying past outbreaks, denser sampling can facilitate the reconstruction of the evolutionary history of this lineage. Previous work, particularly by Key et al. (2020), linked the rise of this serovar to the emergence of farming, and attributed candidate convergent pseudogenisation (gene loss) events to host adaptation ^7^. Pseudogenisation is a common mechanism in the evolution of host specificity in *S. enterica* serovars ^18^, and has also been demonstrated in other bacterial pathogens, for example, *Mycobacterium leprae* ^19^. However, the number of ancient genomes used to make these inferences was relatively low, meaning that the past genetic diversity and lineage structure of *S. enterica* remains poorly understood. This restricted the power of any analyses of convergent adaptation; furthermore, it was not possible to date any of these potentially adaptive pseudogenisation events, which is a unique feature of ancient DNA data.

Another open question raised by previous work was the ability of prehistoric, putatively host-generalist strains to cause systemic infection. Despite the huge genetic diversity observed in *S. enterica* today, nearly all ancient genomes reported thus far fall into the Para C lineage, with lower rates of pseudogenisation than in modern host-adapted lineages. However, the presence of these prehistoric infections in the dental pulp chamber implies an invasive infection ^15^. In addition, the identification of intermediate levels of pseudogenisation in more recent genomes implies a more intermediately host-adapted phenotype, which has yet to be explored in the context of ancient *S. enterica*.

In this study, we report 53 new ancient *S. enterica* genomes, detected from a screening dataset of 8271 individuals, with mean sequencing coverages ranging from 0.16X-116.5X (median: 8.7X) (see **Table S1** for alignment statistics and Figures **S1-S7** for data authentication). Of these, 52 genomes fall into either the Birkenhead sister lineage or the Para C lineage. This includes 14 new genomes falling into a now-extinct basal Para C clade, predominantly from the Bronze Age, first described by Neumann et al. (2022) ^11^, as well as six new genomes in the sister lineage to the modern-day Birkenhead serovar, which we associate with Copper Age and Late Neolithic agropastoralist societies. This increased sampling density allows us to recover a diverse range of ancient genomes, which has also allowed us to uncover ancient sub-lineage diversity and reveal signals of convergent adaptation to mammalian hosts, both within the Para C lineage and across different host-adapted serovars.

### Prehistoric Lineages: Repeated Replacements

We find that *S. enterica* infections were more widespread, both geographically and temporally, than previously reported. We identify *S. enterica* in archaeological tooth samples spanning from the Iberian Peninsula in the west to present-day Mongolia in the east, and from as early as 3634-3383 cal BCE in Spain (SJL020; 2*σ*) to as late as the medieval period in Austria and Mongolia (FSM027 (12th-17th century) and TAV007 (11th-14th century); both contextually dated).

The earliest circulating *Salmonella* lineage identified in this study is a sister lineage to the modern-day Birkenhead lineage, the first genomes of which are detected between 3600-3300 BCE (**Figure 1B, Table S2**). The majority of these cases are associated with European Late Neolithic or Chalcolithic archaeological contexts, with the exception of BOY008, an Early Bronze Age individual from Bulgaria, which forms a clade with the previously published samples IKI003 and IV3002, from Late Chalcolithic and Early Bronze Age contexts in Turkey and southern Russia, respectively (**Figure S8**) ^7^. Another lineage began circulating in Europe around 3300-3100 BCE (**Figure 1B, Table S2**). We term this the “Neolithic/Eneolithic/Bronze Age” (“NEBA”) lineage, as the vast majority of its members are dated to the Bronze Age, with the exception of MAJ022 from an Eneolithic/Copper Age context in Ukraine and OBP001, from a Late Neolithic Swiss context ^7^. Due to insufficient quality for confident phylogenetic placement, we tentatively placed OBP001 basal to the NEBA lineage, which is consistent with Key et al. (2020) ^7^ (**Figure S8; Methods 2.3**). The NEBA lineage spread throughout Europe and Asia and lasted for several thousand years (ca. 3400 BCE-1200 BCE). In contrast, the Birkenhead sister lineage was dated to a much narrower time window (ca. 3650-2500 BCE), and is restricted to south-western Eurasia. However, we note that although the geographic distribution of screened samples does not differ dramatically between the Birkenhead time window and the NEBA time window (**Figure S9**), the total number of individuals screened for this project increases from 770 in the Birkenhead window to 2363 in the NEBA window, so the absence of the Birkenhead sister lineage outside of this range could be due to lower sample numbers. Notably, the two lineages do not appear to be separated by the ancestry of those infected: the Birkenhead group includes two individuals with steppe-related ancestry (BOY008 and IV3002) ^20,21^, while the rest carry Late Chalcolithic farmer-related ancestry. The NEBA lineage largely post-dates the spread of steppe-related ancestry (including the earliest-dated MAJ022), but also includes individuals who do not carry steppe-related ancestry, such as SUA004 from Sardinia, and HGC040 and HGC004 from Crete ^11,22^.

**Figure 1.**
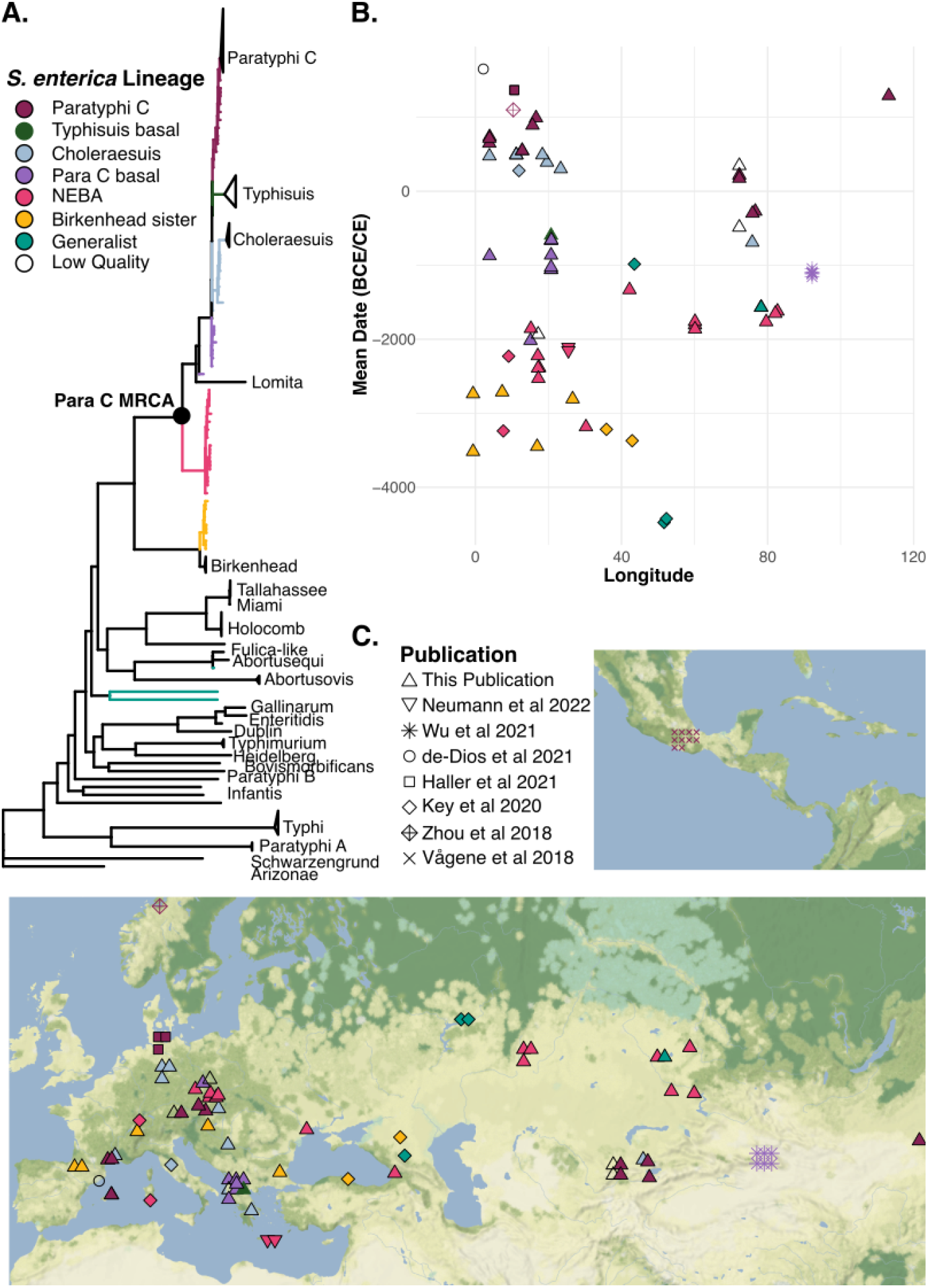
Salmonella-positive samples: geographical location; dates and genetic relationships. **A**. Positions of ancient lineages in the *S. enterica* phylogenetic tree are marked by coloured branches. These lineages are: Birkenhead sister lineage (yellow); NEBA (Neolithic/Eneolithic/Bronze Age) (pink); basal to the rest of the Para C lineage (purple); Choleraesuis (blue) Typhisuis (dark green); Paratyphi C (maroon); non-specific host generalists (turquoise). Modern lineages are denoted by black branches and are labelled. The most recent common ancestor (MRCA) of the Para C lineage is also labelled. **B**. Relationship between sampling date (in years CE/BCE), *S. enterica* lineage assignment (as in **A**), and longitude of each genome (see Table S2 for details). Publication in which each genome was first reported maps to point shapes (as in legend in **C**). **C**. Geographic location of each genome sampled. Paratyphi C genomes from the cocoliztli epidemic in Teposcolula-Yucundaa (reported in Vågene et al 2018) are plotted beside the legend. Map data was obtained from [Stadia Maps](http://stadiamaps.com/) (stadiamaps.com) and [Stamen Design](http://stamen.com/) (stamen.com), and plotted using ggmap.

As well as a lineage turnover associated with this NEBA lineage, we also see the first evidence for a prehistoric *S. enterica* outbreak, which occurred at the site of Nepluyevsky (NEP), where three almost identical *S. enterica* genomes were identified in contemporaneous burials within the same kurgan (**Figure S8, SI Note 1.24**, and Blöcher et al. (2023)^23^). Two *S. enterica* genomes from the basal Para C lineage (KNC041; KNC095) from the site of Kamenice, in modern-day Albania, also likely represent a single outbreak, as they are essentially identical (**Figure S8**). They also have overlapping radiocarbon dates, and the individuals were buried beside each other (**SI Note 1.17**). These observations demonstrate that prehistoric strains of *S. enterica* were already capable of causing invasive, fatal infections, and potentially even epidemic events.

There is a strong temporal and geographic signal in the more derived parts of the tree, although these host-adapted lineages – Paratyphi C (human), Choleraesuis, and Typhisuis (both pig-adapted) – were evolving independently. The Choleraesuis lineage comprises samples primarily from Western Eurasia, although the earliest-diverging sample in this lineage is from an individual from present-day Kazakhstan (CPA002). This sample (796-567 cal BCE (2*σ*)) predates the European samples, which first appear 225-333 cal CE (ETR001; 2*σ*), and it is also relatively divergent from the rest of this lineage. At the same time, the Paratyphi C lineage was also diversifying. The earliest Paratyphi C cases were also found in Central Asia, and they predate the later, primarily European Paratyphi C cases by several hundred years (**Table S2**). By the early medieval period (ca. 700 CE), the Paratyphi C serovar was present throughout Eurasia, and has been detected from as far east as Mongolia (TAV007) to as far west as Spain (ABF001); previous work has shown that it later caused at least one epidemic in Mexico (labelled “Tepos” in Figures 2 and 3) ^6^.

**Figure 2.**
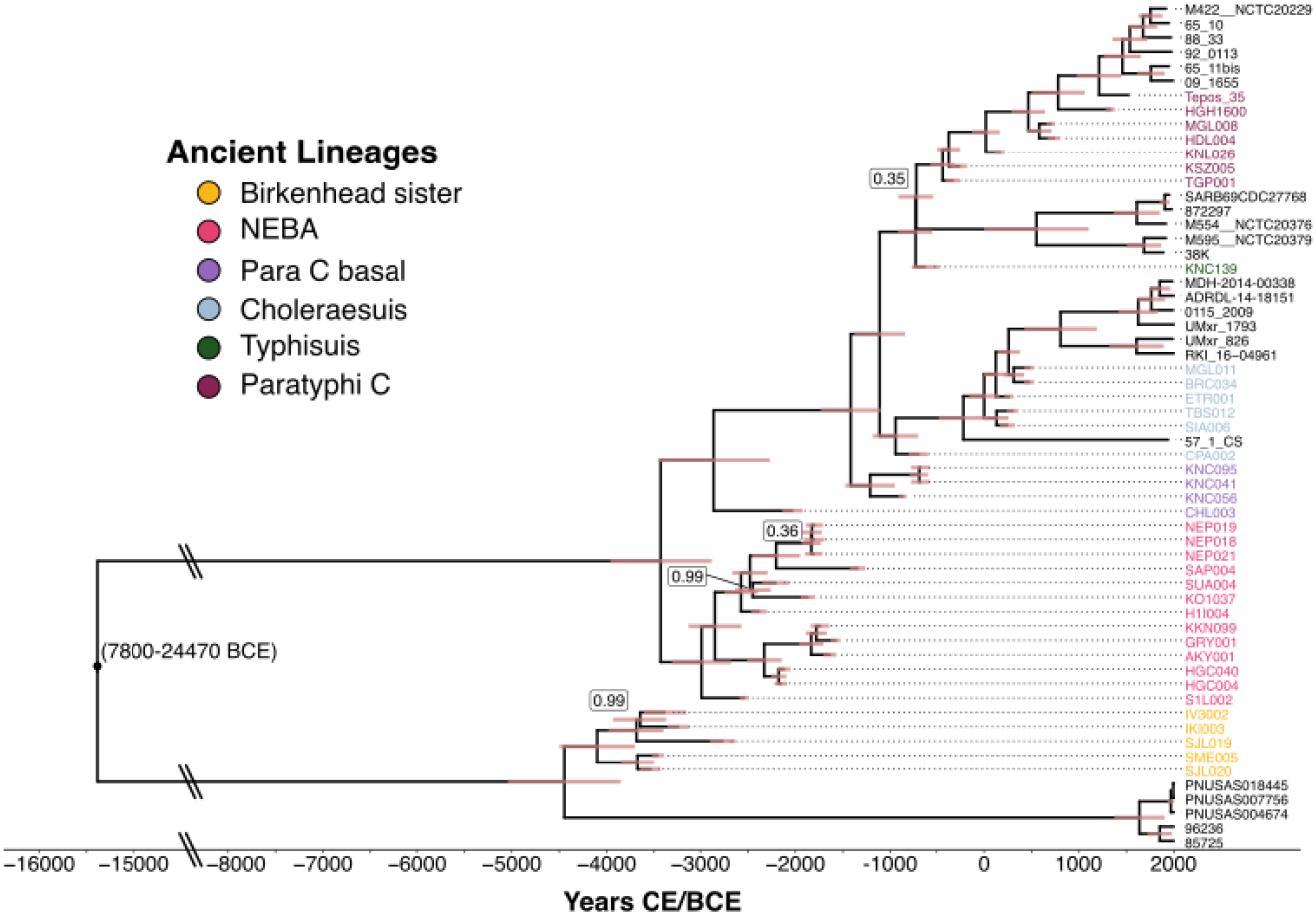
Time-calibrated *S. enterica* Para C lineage phylogeny, constructed using BEAST2. 95% highest posterior density (HPD) heights for each tip and node are represented by error bars, except for the root node, which is labelled with its 95% HPD for ease of visualisation. Similarly, the x-axis (time) is artificially shortened between 8000 BCE and 15000 BCE for ease of visualisation. Ancient genomes are coloured by their lineage label. Any node with posterior support of less than 1 is labelled.

**Figure 3.**
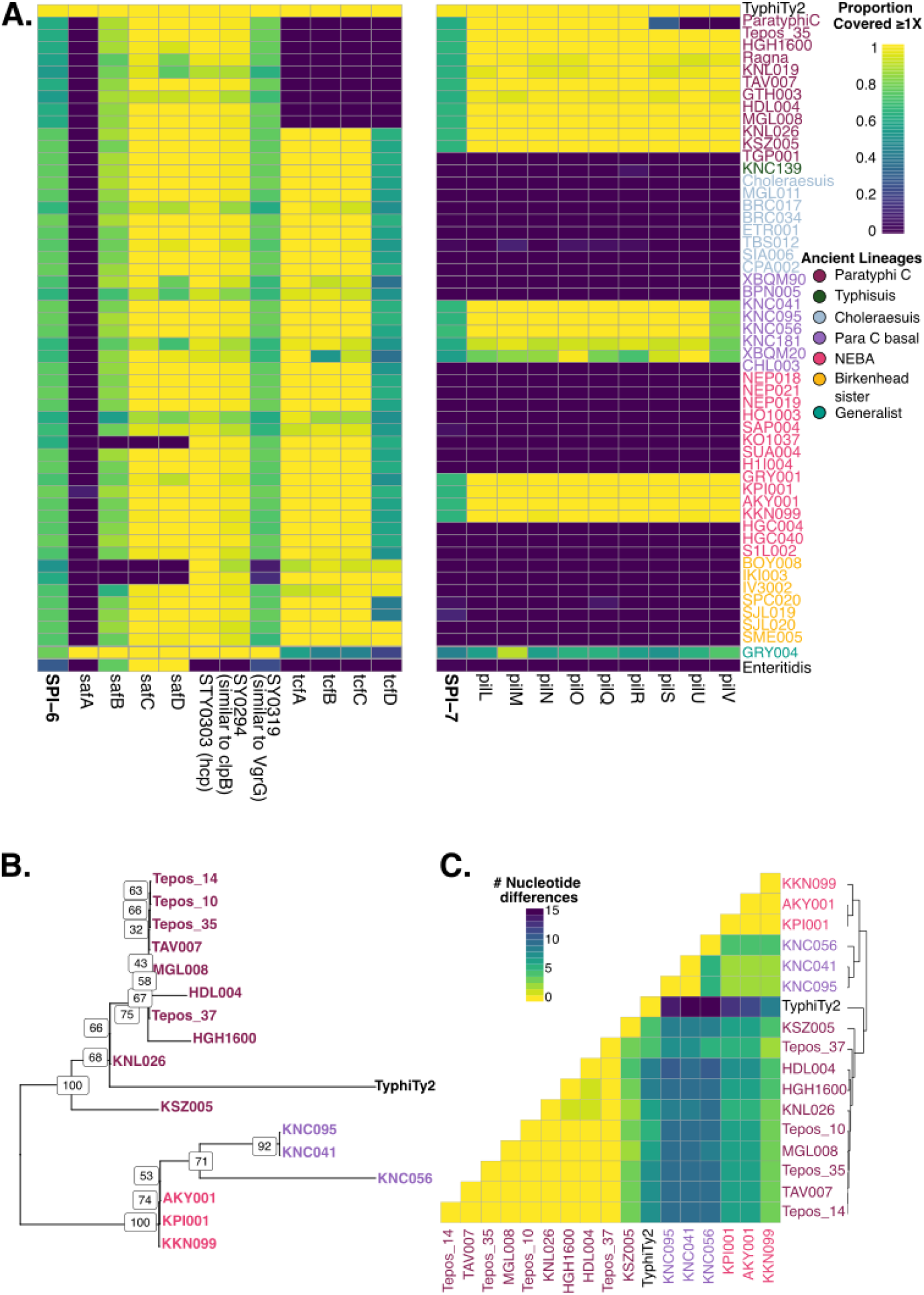
Salmonella Pathogenicity Island (SPI) Acquisition. A. SPI-6 and 7 variability in coverage and presence/absence of a subset of virulence genes. All ancient Para C genomes passing quality filters and with mean coverage ≥3X are included in this plot. Ancient and modern Para C genomes are coloured by their lineage label. Paratyphi C representative: RKS4594; Choleraesuis representative: SC−B67; Enteritidis representative: P125109. Colour within the heatmap represents the proportion of the gene or SPI covered at 1X, where yellow represents a high proportion covered, and dark blue represents 0% covered at 1X. **B. Maximum likelihood tree constructed from samples with maximum per-individual missingness of 25% at pilin locus on SPI-7**. Individual IDs are coloured by lineage, as in parts A and C. **C. Heatmap of nucleotide differences within pilin locus of carriers with maximum per-individual missingness of 25% at pilin locus on SPI-7**. Individual IDs are coloured by lineage, as in parts A and B. The yellow in the heatmap represents fewer differences, and darker colours represent more nucleotide differences. Cluster relationships are shown using the dendrogram.

Our dataset suggests that the most likely location for the diversification of host-adapted Para C lineages is Central or Western Asia. Despite a higher sampling density in Europe during this time period (i.e. 1500-500 BCE) (**Figure S9**), all but one of the individuals identified thus far with evidence of infection by a host-adapted *Salmonella* lineage come from sites in Central Asia. Furthermore, previously published samples from the region of the Tian Shan mountains in Central Asia fall extremely close to this diversification (XBQM samples, **Figure S8**) ^8^. However, we caution that the highest sampling density for this time period is in southeast Europe, which is likely to have much poorer DNA preservation and is therefore more likely to be a source of false negatives in our pathogen screening pipeline. The few European *S. enterica* genomes we have from this time period mainly come from a site in Albania (Kamenice, KNC); these genomes primarily fall into a clade basal to the host-adapted Para C lineages, although they also include a single genome placed at the diversification of the pig-adapted Typhisuis and human-adapted Paratyphi C (KNC139). This ancestral bacterial population may have been circulating among populations across Southeastern Europe and Central Asia.

### Time-Calibrated Phylogenetic Analysis

There is a strong temporal signal within the Para C lineage (R^2^ from root to tip regression = 0.61). Time-calibrated trees were used to infer split times of relevant lineages. The TMRCA (time to the most recent common ancestor) of the first detected circulating lineage of *S. enterica* in human populations in Eurasia (i.e. the Birkenhead sister lineage) was dated to between 5070-3880 BCE (during the European Late Neolithic period); this overlaps with the TMRCA estimated by Key et al. (2020) ^7^. The TMRCA of the Para C lineage is marginally later (3980-2900 BCE), and substantially more recent than previous estimates (14653-4523 BCE). This overlaps with the TMRCA of the NEBA clade (3300-2680 BCE) and the rest of the Para C lineage (3430-2260 BCE). The three host-adapted serovars within the Para C lineage (Choleraesuis, Typhisuis, Paratyphi C) split from each other between 1370-830 BCE, and the two host-adapted strains for which we have ancient genomes started to diversify at around the same time, with Paratyphi C diverging ca. 550-290 BCE and Choleraesuis diverging ca. 1170-700 BCE (**Figure 2**).

### Repeated Pathogenicity Island Acquisition

We next assessed gene gain in the form of *Salmonella* Pathogenicity Islands (SPIs) and dated these acquisitions using our BEAST time tree. SPIs are regions of the *S. enterica* genome acquired through horizontal gene transfer which encode genes involved in virulence, such as pathogenicity effectors ^24^. Here, we followed the approach of Vågene et al. (2018)^6^, by assessing the proportion of annotated pathogenicity islands from the Typhi CT18 reference genome that are covered in our ancient genomes (**Figure S10**). For all ancient samples, SPI-1-5 and SPI-9 were classified as present, while SPI-10 was absent (**Figure S10**). Both SPI-6 and SPI-7 (**Figure 3**) were partially present in both ancient and modern genomes. SPI-6 was partially present in all genomes considered, whereas it appears that SPI-7 has been acquired independently a number of times within the Para C lineage: once at the MRCA of KPI001, KKN099, AKY001 and GRY001 (1950-1690 BCE) (all from Central Asia, in the NEBA lineage); once at the MRCA of the samples from Kamenice (excluding KNC139) and the published sample XBQM20 (1470-950 BCE) (although this SPI appears to be absent in the genome of BPN005, which is placed within this clade), and once at the MRCA of the Paratyphi C lineage, excluding TGP001 (480-230 BCE).

The gene content of the acquired SPI-7 is close to identical to that of the modern Paratyphi C SPI-7, but also includes some pilin-related genes absent in the Paratyphi C reference genome, previously noted in Vågene et al. (2018)^6^. Interestingly, SPI-7 appears to be present at a low coverage in GRY004 (**Figure 3, Figure S10**), which falls with host generalist strains in the phylogeny (**Figure 1, Figure S8**). However, unlike other SPI-7 positive genomes, there is a substantial drop in mean coverage across this region (**Figure S11**). This could point to a co-infection; however, heterozygosity levels for GRY004 are not substantially different to the heterozygosity levels observed for other genomes (**Table S8, Figures S4-5**). An alternative hypothesis is that SPI-7 was integrated into a sub-population of bacteria infecting this individual, which is why we see a reduction in coverage at this locus without any excess of heterozygosity genome-wide. This points to an environmental origin for SPI-7, as suggested by other authors ^25^, with a geographical origin in Central Asia, supported by our observation of the first case of each independent acquisition in genomes from Central Asia.

In terms of SPI-6 gene content, it appears that the gene *tcfD* (encoding the N-terminal domain of a fimbrial adhesin, which interacts with host cell receptors ^26^) has been mostly lost in the Para C lineage (**Figure 3A**). The proportion of this gene covered is well below 90% in samples where it is covered at all, and seems to have been completely lost in the MRCA of modern-day Paratyphi C and HDL004/MGL011 (290-650 CE). However, this low coverage is also seen in the Choleraesuis genome used for comparison, which has a version of *tcfD* annotated in its genome ^27^. This suggests that *tcfD* is in fact present in these earlier Para C genomes, but is sufficiently diverged from the Typhi version of the gene that alignment is substantially worse than for the other components of this fimbrial operon (*tcfABC*).

The pilin gene *safA* also appears to be absent from all Para C genomes (as well as from the more basal NEBA and Birkenhead clades), but *safB* (which performs a chaperone function for the pilin gene *safA*) is present throughout the Para C lineage (**Figure 3A**). Similarly to *tcfD*, there is an annotated version of *safA* in Choleraesuis genomes ^27^, as well as in the Paratyphi C reference used ^28^. Further assessment of alignment to the Paratyphi C reference shows that the Paratyphi C *safA* gene is covered in all ancient genomes, with the exception of the non-Para C GRY004, the two Birkenhead sister lineage genomes BOY008 and IKI003, and the NEBA lineage genome KO1037 (**Figure S12**). This suggests that even in cases where highly variable genes appear to be partially present, they may in fact be fully present in these genomes. This is likely due to high levels of sequence variability, impacting mapping. These results highlight the relevance of surface proteins in the evolution of pathogenic *S. enterica*, as well as the challenges of assessing presence or absence of highly variable genes.

The most variable pathogenicity island genes in our dataset are host-interacting proteins or domains of proteins, such as pilin and fimbrial proteins. The importance of secreted and surface proteins has been highlighted in previous studies of host-specific *S. enterica* strains ^29^. While fimbriae are important in gut colonisation of enteric pathogens ^30^, the loss of fimbriae, whether by phase variation or pseudogenisation is advantageous in evading host immune responses. Mice infected with *S*. Typhimurium knockouts for fimbrial genes showed increased bacterial dissemination to the bloodstream, compared to strains expressing fimbrial genes^31^, and Gewirtz et al. (2001) show that fimbriae induce IL-8 secretion in epithelial cells^32^. It is notable that other related proteins, such as other genes in the tcf operon and *safB*, do not exhibit as much variability. This suggests that they are under much lower host-induced selection pressures.

Sequence variation at the pilin locus, which has been lost in some modern Paratyphi C genomes, was assessed in SPI-7 carriers, and compared to the Typhi Ty2 pilin sequence. We restricted the analysis to individuals with a maximum of 25% missingness at this locus and performed a multiple sequence alignment using ClustalW^33^. There were zero nucleotide differences between all Paratyphi C lineage carriers, except for the earlier-diverging KSZ005 genome, which has approximately 5 nucleotide differences between it and all more derived ancient Paratyphi C genomes. Interestingly, this sequence is also more closely related to the pilin locus sequence found in serovar Typhi strain Ty2 than any of the other ancient genomes (5 nucleotide differences) (**Figure 3C**), and seems to branch basally to it in a midpoint-rooted maximum likelihood tree constructed from this sequence alignment (**Figure 3B**). The NEBA and Kamenice (KNC) pilin sequences also cluster separately, consistent with our hypothesis of multiple independent SPI-7 acquisitions. The products of the pilin locus in SPI-7 have been shown to mediate inflammatory signal transduction in immune cells infected with serovar Typhi^34^.

One pathway for *S*. Typhi adhesion and invasion of gut epithelial cells has been shown to occur via the Cystic Fibrosis Transmembrane Receptor (*CFTR*) using the PilS gene product, which is encoded by this pilin locus on SPI-7 ^35,36^. There is no nonsynonymous sequence difference between the Paratyphi C carriers for the *pilS* locus and the serovar Typhi *pilS* gene, and one predicted amino acid difference between the earlier NEBA and Kamenice (KNC) carriers and the Typhi sequence (**Table S3**), suggesting that this could be a mechanism contributing to invasive disease in Paratyphi C strains. It has also been suggested that the most common cystic fibrosis-causing mutation (CFTR ΔF508) has been under balancing selection due to past exposure to typhoid fever ^35^. However, *S. enterica* serovar Typhi has yet to be reported in the ancient DNA record. The abundance of ancient, invasive Para C lineage infections carrying SPI-7 and this pilin locus offers an alternative hypothesis for a selective pressure. The earliest direct detection of this variant was reported to be approximately 2200 years ago in Britain, with no evidence for positive selection ^37^; however, given the severe fitness cost to carrying two copies of this allele (as well as some reports of increased morbidity and therefore reduced fitness in heterozygous carriers ^38^), more careful modelling of balancing selection at this locus could help to resolve this question.

Another locus encoded by SPI-7 is the Vi antigen locus, which has also been shown to be important in host-pathogen interactions as it allows bacteria to evade host immune response (via reduced complement activation) ^39^. Its regulator, TviA, has also been shown to prevent NF-κB activation by regulating flagellar genes and the type 3 secretion system T3SS-1 ^40^.

SPI-7 covers a large part of the genome (134 kB in S. Typhi), and may impact chromosomal stability ^41^. It is spontaneously lost when passaging S. Typhi in culture ^42,43^, implying that its maintenance in *S. enterica* populations depends on host-driven selection pressure ^41^. The maintenance of SPI-7 in sub-populations of ancient Para C lineage genomes implies that, even in the NEBA lineage, this lineage was already infectious enough to benefit from the immune evasion and virulence phenotype provided by this genomic island. However, it appears to have been lost within the Kamenice/XBQM20 clade in the BPN005 genome, suggesting that this hypothesised selection pressure was not strong enough to prevent its loss in that genome.

The origin of SPI-7 in the Para C lineage remains an open question. Notably, the earliest genomes in our dataset containing this island all originate from a similar geographic location (Central Asia); perhaps this pathogenicity island was maintained in an as yet unsampled reservoir of diversity there. This hypothesis is supported by the detection of low levels of SPI-7 in the host-generalist GRY004 genome from Kazakhstan. The subsequent spread of SPI-7 containing genomes to Southeast Europe (Kamenice) and later globally (in the Paratyphi C lineage) may reflect synergistic interactions between the acquisition of this pathogenicity island and additional host adaptations.

### Convergent Pseudogenisation

We assessed functional sequence variation within our modern and ancient dataset in order to investigate signals of convergent host adaptation. This consisted of a balanced subset of genomes with known serovar designations and host specificities, as well as high-quality ancient genomes (see **Table S4 and Methods 2.5**). As *S. enterica* has an open pangenome, we realigned all sequencing data and modern pseudo-reads to the portion of the *Salmonella* pangenome which are fully covered by the probes used to capture the ancient *S. enterica* genomes, as in Key et al. (2020) ^7^ (see **Methods 2.5** for full details of realignment, filtering and variant calling).

First, the relationship between pseudogenisation rates, host specificity and time was assessed. There was a clear association between host-specific serovars and an increased pseudogenisation rate, as previously reported (**Figure S13**) ^7,44,45^. The rates of pseudogenisation vary widely between ancient samples; this seems to have a reasonably linear relationship with estimated sample date (Pearson correlation coefficient: 0.7951) (**Figure 4A**), suggesting that the accumulation of pseudogenes, and adaptation to specific hosts, was a gradual process. Only the relatively recent TAV007 (Mongol-era Mongolia) and Tepos (colonial Mexico) genomes reach the rate of pseudogenisation seen in modern isolates of human-adapted Paratyphi C. However, older, more ancestral Paratyphi C genomes do cluster with modern isolates based on shared pseudogenes (**Figure S14, S15**), suggesting that these ancient genomes are likely semi-host-adapted, i.e. they have some, but not all, of the adaptations required for host specificity. For example, the pseudogene *slrP* was identified in the Para C ancient genomes, and has also been associated with host specificity ^46^.

**Figure 4.**
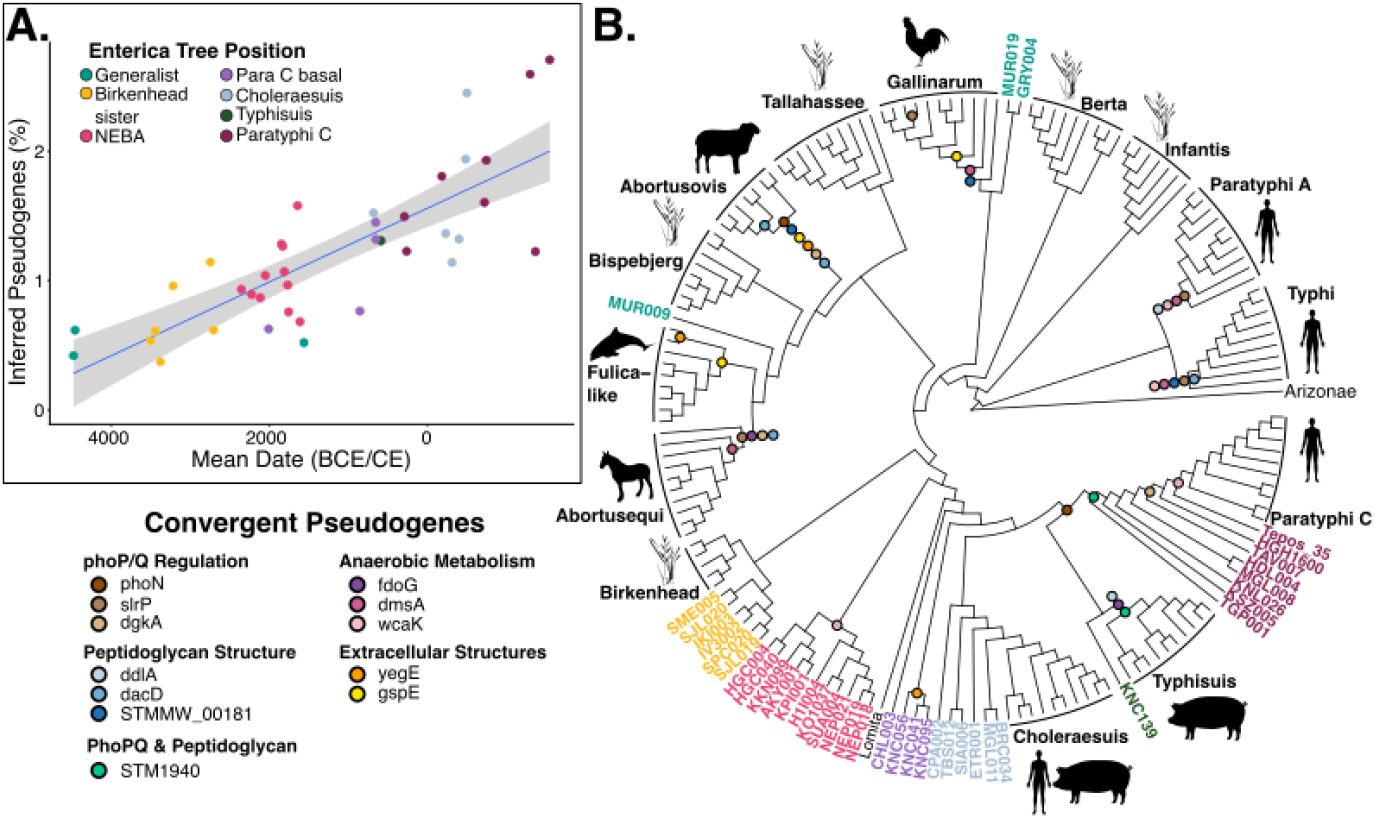
A.Relationship between pseudogenisation rate and time in ancient *Salmonella enterica*. Genomes are restricted to those with a minimum mean coverage of 5X. The midpoint of either the 2-sigma radiocarbon date or the contextual date is used as the mean date for plotting. Ancient genomes are coloured by phylogenetic lineage as in Figure 1. **B. Cladogram displaying convergent pseudogenisation in host-adapted lineages**. Inferred pseudogenisation events discussed in the main text are marked on this figure. Host-adapted lineages (Typhi, Paratyphi A, Gallinarum, Abortusovis, Fulica-like, Abortusequi, Choleraesuis, Typhisuis, and Paratyphi C) are denoted by the outline of the host. Host-generalist lineages (Infantis, Berta, Tallahassee, Bispebjerg, and Birkenhead) are denoted by a plant icon. Ancient sample IDs are coloured by phylogenetic lineage as in **Figure 3A**. Species icons obtained from phylopic.org.

We then sought to assess whether particular pseudogenisation events were associated with host specificity. This was first analysed by associating the presence of a pseudogene with inferred host specificity based on serovar using a chi-square test (see **Methods** for full details) (**Table S5**). Here, we see some enrichment for GO terms related to transmembrane transport, localisation, pilus assembly, carbohydrate catabolism, as well as terms relating to cell membranes and extracellular space. This seems to suggest that key functions or pathways that are modified when adapting to a specific host niche tend to relate to the microbe’s interaction with its environment, a biologically plausible pattern also reported in analyses of convergent adaptation in serovars Paratyphi A and Typhi ^29^.

However, this approach has a number of limitations. It is naïve to both phylogenetic information and whether different pseudogenising mutations are observed in the same or different serovars. Therefore, it is not informative about whether a pseudogene is observed in multiple host-adapted serovars due to convergent evolution, a shared common ancestor, or, potentially, due to recombination between different serovars. In addition, this initial analysis did not include any ancient genomes, as these cannot reliably be labelled as host-specific or host-generalist. Furthermore, this approach does not account for differences in substitution rates between different genes, for example due to differences in gene length or GC content.

As an alternative, we analysed convergent pseudogenisation in modern and ancient genomes using the tool SNPPAR to place each pseudogenising, missense, and synonymous mutation at different nodes in the tree ^47^. In order to mitigate the impact of missingness on this analysis, we restricted the analysis to the core genome (i.e. genes covered ≥90% at 1X in 95% of the samples), leaving a total of 3158 genes for analysis. We assessed genes with at least two independent (i.e. not due to recombination or shared ancestry), fixed pseudogenisation events, and defined excessive pseudogenisation as those with the highest (top 2.5%) difference between the observed and expected number of pseudogenes, termed delta pseudo, based on observed synonymous mutations, which we assume to be neutral. We then restricted this candidate list to those where the inferred pseudogenisation event happens at an ancestral node, and where there is no pseudogenisation event in host-generalist lineages. This left 26 genes with an excess of pseudogenisation in host-specific strains relative to host-generalists (**Table S6, SI Note 4**).

These candidate genes tend to show significant associations with host-specificity in the more naïve chi-square test (**Table S6**), suggesting that, despite its limitations, this test is also recovering biologically meaningful associations between pseudogenes and host specificity. It also allows us to interrogate the role of non-core gene pseudogenisation in host adaptation. This is particularly of interest, as the absence of an accessory gene is functionally equivalent to a pseudogenising mutation, so by restricting our analysis to the core genome, we are losing some meaningful biological signal.

A number of common themes emerge when investigating convergent pseudogenes. Of the 26 candidates, at least four (*phoN, slrP, dgkA, STM1940*) are regulated by the PhoP/Q two component system (**Figure 4B**). The PhoP/Q system is a regulatory system in a number of gram-negative species, and is central in regulating virulence traits in *S. enterica* ^*48*^. It primarily responds to Mg^2+^ levels, and is induced under magnesium starvation conditions, although a subset of phoP/Q regulated genes are also activated in response to acidic pH and cationic antimicrobial peptides ^48^. About 3% of the *S*. Typhimurium genome is regulated in some way by the PhoP/Q system, either directly or indirectly ^49^. The functions it regulates are central to host-pathogen interactions, including inverse regulation of genes encoded by SPIs 1 and 2, which inhibit mammalian epithelial cell invasion and promote intracellular survival once engulfed by macrophages ^50,51^. One of the ways in which pseudogenisation is thought to contribute to the evolution of pathogenicity in bacteria is by “re-wiring” regulatory networks ^52^. The importance of the PhoP/Q two component system in virulence ^48^, and our observations of multiple host-specificity-associated pseudogenes in its regulatory network suggest that changes in this network were important in the evolution of host specificity in these *S. enterica* lineages.

Another subset of these candidate pseudogenes (*fdoG* and *dmsA*) is involved in different aspects of anaerobic metabolism (**Figure 4B**). Anaerobic metabolism is particularly important in gut colonisation, but less relevant for systemic infections ^53^. Another gene disrupted in Typhi, Paratyphi A, Paratyphi C, and the NEBA lineage was *wcaK*, although this did not pass our threshold for excessive pseudogenisation (delta pseudo: 1.6412; threshold: 1.642). The colanic acid biosynthesis pathway (which includes the gene product of *wcaK*) has previously been shown to be disrupted specifically in human typhoid and paratyphoid fever-causing strains ^53^; here, we also observe its disruption in the NEBA *S. enterica* lineage (**Figure 4B**), demonstrating that some of these previously-reported human adaptations may have been acquired independently in this lineage.

We also observe a number of pseudogenes involved in peptidoglycan structure (*ddlA, dacD, STM1940, STMMW_00181*), as well as in functions related to the regulation and assembly of extracellular proteins such as pilins (*yegE, gspE)* (**Figure 4B**), highlighting that there may be some differences in cell envelope composition between some host-specific and host-generalist serovars. Several sub-lineages of non-typhoidal *S. enterica* serovars have been shown to be much more likely to cause invasive, systemic infections (for example, *S*. Typhimurium ST313 and *S*. Enteritidis ST11) ^54^. As with typhoidal serovars, these invasive sublineages show elevated rates of pseudogenisation, although to a lesser extent than typhoidal serovars ^55–57^. The gene *STM1940* (a cell-wall associated hydrolase) has also been demonstrated to be pseudogenised in *S*. Typhimurium ST313 ^58^. In addition, regulatory changes in *csgD* (which regulates the synthesis of curli fimbriae) has been shown to impact biofilm formation in this iNTS lineage, suggesting that human-to-human transmission is more relevant in ST313 ^56^. The gene *yegE* is required for curli expression in *Escherichia coli* and has also been linked to curli expression in *S*. Typhimurium ^*59,60*^. The pseudogenisation of this gene in the basal Kamenice (KNC) clade, along with the presence of SPI-7 in these genomes, could also reflect a greater importance for human-to-human transmission even at this early stage.

## Conclusions

The quantity of ancient genomic data, particularly from humans, has increased rapidly in the past decade ^61^. With this, the ability to mine the non-host data for evidence of past infections has improved drastically, allowing for the analysis of the evolutionary history of important human pathogens ^15^. Here, we increase the representation of ancient *S. enterica* by reporting 53 new genomes, and co-analyse them with 21 previously reported ancient *S. enterica* genomes ^5–11^. We identify two cases of likely prehistoric invasive *S. enterica* outbreaks at the sites of Nepluyevsky (Russia) (ca. 2010-1630 BCE) and Kamenice (Albania) (ca. 770-540 BCE), and show that these genomes either carry pathogenicity islands associated with systemic infection in modern genomes (Kamenice), or seem to carry at least one pseudogene related to host specificity (*wcaK*; Nepluyevsky). These observations demonstrate that although prehistoric *S. enterica* lineages did not display the characteristic high pseudogenisation rate seen in host-adapted lineages today ^7^ (**Figure S13**), they were still capable of causing invasive, fatal infections and did have some genetic characteristics associated with invasive infections today. This suggests that host adaptation was likely a gradual process over time, with prehistoric, intermediately-adapted strains able to cause systemic infections and outbreaks, despite lacking the full repertoire of pseudogenes identified in more recent isolates.

The increased sample density in this study also enabled more precise molecular dating of lineage emergence than was previously possible. The inferred TMRCA of both early circulating lineages (NEBA and Birkenhead sister lineages) corresponds to the European Late Neolithic period, highlighting the role of Neolithisation in the emergence of early circulating *S. enterica* infections ^7^. However, the widespread dispersal and adaptations observed in the prehistoric NEBA lineage date to the Bronze Age, emphasising the role of later innovations, particularly in livestock management, in the evolution of invasive infectious disease.

We also identify several signals of convergent adaptation and invasiveness throughout the dataset, showing repeated acquisitions of the pathogenicity island SPI-7, as well as convergent pseudogenisation in host-adapted lineages. Additionally, we identify a conserved host-interacting pilin gene on SPI-7, which has been implicated in the invasion of gut epithelial cells by *S*. Typhi, and has been hypothesised to have driven balancing selection on the most common Cystic Fibrosis-causing mutation (ΔF508) in Europeans today ^35,62^. Although *S*. Typhi has not yet been identified in the ancient DNA record, we demonstrate that SPI-7 carrying strains of Paratyphi C (and earlier Para C lineages) were widespread throughout Eurasia, offering an alternative mechanism by which selection could have acted.

We demonstrate convergence between pseudogene functions in our ancient *S. enterica* genomes and in emerging iNTS strains today. Invasive non-typhoidal *Salmonella* had previously been linked to infections primarily in the very young (particularly in those also infected by malaria or suffering from nutritional deficiencies), but the HIV pandemic has led to a large population of immunocompromised adults who were susceptible to iNTS infections, uncovering a permissive niche for the evolution of host specificity in these strains ^3,63^.

This raises the question: under what conditions was the Para C lineage evolving that allowed for the evolution of host specificity? Previous work has linked the emergence of this lineage with Neolithisation, i.e. the transition to a sedentary lifestyle based on agriculture and animal husbandry^7^. Increased contact with domestic animals creates a more permissive environment for the exchange of pathogens, as demonstrated by several studies ^64–66^. During the Bronze Age, more widespread intensification of livestock use for secondary products of milk and wool, in addition to the incorporation of horse riding into animal management strategies allowing for larger herds and greater mobility, likely enhanced the exchange of zoonotic pathogens ^67–69^. In tandem with this closer and more sustained relationship with livestock, growing human populations and social networks during this period inevitably facilitated the spread of infectious diseases^70–73^, such as the NEBA lineage.

Another possible factor in the evolution of host specificity relates to the immune status of people infected by these prehistoric lineages. Malnutrition is an important contributor to immunocompetence^74^. For example, periods of drought and famine could contribute to malnutrition-driven immunodeficiency, creating a more conducive environment for invasive infectious diseases. Marciniak et al. (2022) demonstrate reduced stature relative to predicted height in Neolithic European populations, which the authors suggest is due to increased physiological stress and nutritional deficiencies, although this “dip” in height appears to recover in post-Neolithic populations ^75^. In more recent periods, the 6th century CE “Late Antique Little Ice Age” has been suggested to contribute to a high burden of infectious disease, including the first plague pandemic^76^.

For the Bronze Age Trans-Ural region, at least, palynological research indicates that climatic changes would not have been a trigger for this increased infectious disease risk, suggesting newly intensified animal husbandry and settlement dynamics played a more important role here ^77^.

In addition to nutrition-related stresses, other invasive infectious diseases such as plague had been infecting Eurasian humans for as long as *S. enterica*, and the most recent estimate of the *Yersinia pestis* TMRCA is 7951-5844 years BP (i.e. 6000-3894 BCE) ^78^, which overlaps with our inferred TMRCAs for both the Birkenhead (5070-3880 BCE) and Para C lineages (3980-2900 BCE). Furthermore, both *S. enterica* Para C and *Y. pestis* phylogenies have extinct prehistoric lineages – LNBA^−^ for *Y. pestis* ^79,80^ and NEBA for *S. enterica* – which were both widespread throughout Eurasia during the same time period. Moreover, both *S. enterica* and *Y. pestis* infections have been identified in the same site in Bronze Age Crete ^11^. Although no co-infections of *Y. pestis* and *S. enterica* have been identified to date, repeated exposure to infections in susceptible populations likely allowed for a more permissive host niche for the evolution of host specificity, similar to what has been observed for African iNTS lineages.

The emergence of *S. enterica* as a major human pathogen may have been influenced by other invasive diseases, nutritional stress, and the emergence and development of livestock management in the Neolithic and Bronze Age. Further work analysing *S. enterica* from ancient domesticated animals could be particularly informative about the impact of human-animal interactions on *S. enterica* Para C transmission. Additionally, more explicit comparisons of prehistoric pathogen load and markers of malnutrition and other forms of physiological stress could be extremely informative about the kinds of infections humans were exposed to and how this impacted both host and pathogen evolution.

## Data and Code Availability

Sequencing data generated for this study has been deposited in the European Nucleotide Archive, under accession PRJEB97698.

Code needed to reproduce analysis is provided in the github repository https://github.com/iseultj/ParaC_salmonella.

## Acknowledgments

Data was produced by the Ancient DNA Core Unit of the Max Planck Institute for Evolutionary Anthropology which is funded by the Max Planck Society. The *Homo sapiens* outline in Figure 4 was created by Katy Lawler and is licensed under [CC BY 4.0](https://creativecommons.org/licenses/by/4.0/) and obtained from phylopic.org. All other species outlines are in the public domain, and were also obtained from phylopic.org.

## Author Contributions

A. Herbig and J.K initiated and supervised the study. M.A.S supervised the initial phase of the study. M.McC, A.S-N, H.R, T.H, V.V-M, P.W.S, A. Childebayeva, C.W, W.H, J.K, and A. Herbig coordinated the collection of samples for genetic analysis. K.W.A, A. Beisenov, L. Bejko, N.B, M. Berner, M. Binder, L. Bitadze, M. Bondár, I. Bruyako, I. Bugarski, A. Buzhilova, A. Charami, K.D, J.D, M.D.Z.B, L.D, D.R-C, P.D.D, S. Eggers, S. Ellingvåg, M.E, M.F, B. Gimeno Martínez, B.H-G, A. Hałuszko, M.H, D.H, V.I, M. Karapetian, E.K, Y.F.K, R.K, M. Krošláková, T.K, S.L, M.A.L, H.M, I.M, B.G. Mende, I.V.M, V.K.M, N.M-R, A.M, L.M, E.M, N.N, J.O, M.O, D.P-K, A.P, M.V.P, J.P, P.R, C.R, R.R, M.R, Z.S, D.R.S, S.S, E.S.O, A. Sobotkova, A. Stobbe, T.S.V, E. Stolarczyk, N.T, A.A.T, S.T, E.U, S.V, D.B, and A.Z participated in archaeological excavation, provided anthropological assessment and/or contributed to the curation of skeletal materials. G.U.N, A.N.H, M. Bretos, A.F, D.G, X.J, M. Keller, L.M, L.P, and E. Skourtanioti managed samples and metadata and/or conducted laboratory work. G.U.N, A.A.V, and A.N.H conducted metagenomic screening of ancient sequencing datasets. I.J, G.U.N and A. Bhattacharya performed bioinformatic analysis. I.J curated the ancient salmonella datasets and performed computational analyses. A-T.M compiled archaeological metadata and edited site descriptions. All authors contributed to the interpretation of the data. I.J and A. Herbig drafted the manuscript with contributions from all co-authors.

