## Supplementary Information for "Evolutionary history and recurrent host adaptation in ancient *Salmonella enterica*"

Iseult Jackson[1,*,✉], Gunnar U. Neumann[1,*], Anna-Theresa Mayr[2], Aida Andrades Valtueña[1], Alina N. Hiss[1], Lyazzat Musralina[19,43], Kurt W. Alt[3], Arman Beisenov[4,5], Lorenc Bejko[6], Natalia Berezina[7], Margit Berner[8], Aparajita Bhattacharya[1], Michaela Binder[9], Lia Bitadze[10], Mária Bondár[11], Marina Bretos[12], Igor Bruyako[13], Ivan Bugarski[14], Alexandra Buzhilova[7], Alexandra Charami[15], Kevin Daly[16], Jocelyne Desideri[17], Marta Díaz-Zorita Bonilla[18], Leyla Djansugurova[19], Déborah Rosselet-Christ[17], Paula Doumani Dupuy[20], Sabine Eggers[8], Sturla Ellingvåg[21], Michal Ernée[22], Mirosław Furmanek[23], Anja Furtwängler[1], Dániel Gerber[24], Belen Gimeno Martínez[25], Brigitte Haas-Gebhard[26], Agata Hałuszko[27,28], Michaela Harbeck[29], Daniela Heilmann[2,30], Vujadin Ivanišević[31], Xiaowen Jia[1], Marina Karapetian[7], Marcel Keller[32,33], Elmira Khussainova[19], Yuri F. Kiryushin[34], Rüdiger Krause[35], Mária Krošláková[36], Thomas Kubenz[37], Stephanie Larson[16], Maria A. Liston[38], Hellen Mager[1], Igor Manzura[39], Michael McCormick[40,41], Balázs G. Mende[24], Ilya V. Merts[42], Victor K. Merts[42], Nataša Miladinović-Radmilović[14], Arnold Muhl[37], Erdene Myagmar[44], Nicole Nicklisch[3], Jörg Orschiedt[37], Mathieu Ott [45,46], Doris Pany-Kucera[8], Luka Papac[1], Aleksandra Papazovska[47], María Victoria Pastor[48], Jaroslav Peška[49], Piroska Rácz[11], Claude Raynaud[50], Roberto Risch[51], Matej Ruttkay[36], Zainolla Samashev[52,53], David R. Scahill[54], Svetlana Sharapova[55], Elena Sintes Olives[56], Eirini Skourtanioti[1], Adéla Sobotkova[57], Astrid Stobbe[35], Tatjana Stojoska Vidovska[58], Eliza Stolarczyk[35], Anna Szécsényi-Nagy[24], Nino Tavartkiladze[10], Alexey A. Tishkin[34], Solenn Troadec[40], Emma Usmanova[5,59,60], Diana Vicente[48], Dragana Vulović[14], Aidyn Zhuniskhanov[61], Maria A. Spyrou[1,70],Harald Ringbauer[1], Taylor Hermes[1,62], Vanessa Villalba-Mouco[1,63], Philipp W. Stockhammer[1,2,64], Thilo Fuchs[65], Ainash Childebayeva[1,66], Christina Warinner[1,67,68,69], Wolfgang Haak[1], Johannes Krause[1,✉], Alexander Herbig[1,✉]

### 1. Archaeological Information

#### 1.1. Ciutadella de Menorca, s'Albufera: ABF

*Country:* Spain
*Region:* Menorca, Baleares
*Coordinates:* 40.0025°, 3.8353°
*Sample Date (ABF001):* Iron Age – Byzantine, 603–773 cal CE (2σ)
*Radiocarbon Date:* 1370 ± 34 BP; 603–773 cal CE (2σ)
*Excavation/Sample Provenance:* Rescue excavations by Elena Sintes Olives in 2013.
*Contact People:* Elena Sintes Olives; Roberto Risch

*Site Description:*
Two interments were discovered within an Iron Age hypogeum, which came to light during the clearing of an abandoned field by workers, who intended to transform the area into a garden. The stratigraphy and chronology of the site of the Ciutadella are complex. The initial function of the site appears to have been as a collective burial; some of the Iron Age remains are cremated, while others are not, with the deepest strata exhibiting a grey colour. The hypogeum consists of a broad hall and two lobes, separated by a pilaster at the base. At some point, the roof collapsed, and the pilaster was truncated. The resulting void was subsequently filled with soil, presumably sourced from an external location, in preparation for converting the site into an orchard, which also involved the drilling of a well in one corner. Numerous Iron Age remains were recovered from the hall, while few artefacts were found in the lobes where the ground elevation was higher. However, two long brown patches were observed at the base of the damaged pilaster amidst the grey sediment, indicative of two interments.

- **ABF001 (CIU-03-AR-1 samples A *and* B)** – *S. enterica* was recovered from individual CIU-03-AR-1, who was an adult male positioned on his right side.

#### 1.2. Ak-bauyr: AKY

*Country:* Kazakhstan
*Region:* Ulan district, Ustkamenogorsk
*Coordinates:* 49.6806°, 82.6833°
*Sample Date:* 1686–1538 cal BCE (2σ)
*Radiocarbon Date (AKY001):* 3344 ± 18 BP, 1686–1538 cal BCE (2σ)
*Excavation/Sample Provenance:* Investigations between 2020 and 2021.
*Contact People:* L. Djansugurova; Z. Samashev; L. Musralina; E. Khussainova

*Site Description:*
Ak-Bauyr is located 3 km north of the village of Besterek, Ulan district (East Kazakhstan region) on the slope of Mount Ak-bauyr and 38 km from Ust-Kamenogorsk, Ulan district. Between 2020 and 2021, a mound with “whiskers” was investigated at the Kyzyltas burial ground. The central feature is a circular stone mound measuring 10.5 m in diameter and 30–40 cm in height. Beneath it, an oval grave pit (3.4 m × 2.4 m, 60 cm deep) was identified. Extending southeast and northeast from the mound are curved stone alignments—referred to as “whiskers.” The northern alignment measures

approximately 140 m in length and 1.5-4 m in width; the southern is about 110 m long and 1.5-3 meters wide. Excavated materials include pottery fragments and scattered animal bones.

Excavations revealed two cultural layers (20–50 cm deep) containing stone architectural remains. In the southwestern sector, a rounded structure—possibly a dwelling—was identified next to accumulations of stones likely associated with farm buildings. Also, approximately twelve hearths were found. A particularly enigmatic find consists of three human skulls located within a small stone-enclosed area. The dating of these structures remains debated, with hypotheses pointing to either the Early Nomadic period or the Early Middle Ages. The mounds are primarily associated with solar and fire cults, as well as ancestor worship. Some researchers interpret the "whiskers" as possible astronomical markers designed to track solar movements.

- **AKY001 (Burial 1, cyst box 1):** *S. enterica* was recovered from a tooth from individual AKY001.

##### 1.3. Boyanovo: BOY

*Country:* Bulgaria
*Region:* Yambol Oblast
*Coordinates:* 42.2819°, 26.6294°
*Sample Date (BOY008):* Early Bronze Age, Yamnaya Culture, 2899-2697 cal BCE (2σ)
*Radiocarbon Date (BOY008, tooth, MAMS-47227):* 4213 ± 24 BP, 2899-2697 cal BCE (2σ).
*Excavation/Sample Provenance:* Excavated during archaeological rescue excavations in 2010.
*Contact Person:* Adéla Sobotkova

*Site Description:*
The excavation site is located on top of a limestone quarry in southeastern Bulgaria, ca. 2.7 km east of the Tundzha River and 2 km north of Boyanovo village, within the Yambol province. It is positioned at the southern edge of the Thracian Plan and the area is marked by east-west ridge-lines rising above the river valley, a landscape supporting agricultural settlements from the Late Neolithic and mound-building activity from the Early Bronze Age onward. The site lies within a forest-steppe ecotone, transitioning between Pontic coastal zones, steppe, and montane forest systems.

Excavations were carried out in 2010 as part of a rescue archaeology campaign prior to limestone quarry expansions. Of the three mounds, the first mound was constructed atop a prominent limestone outcrop as the largest mound (45 m in diameter and 4.8 m in height) and contained 20 burials with 23 individuals. The second mound yielded no burials. The third mound was the smallest of the cluster of three, measuring 23 m in diameter and 1.6 m in height. Mound 3 lay some 300m southeast of mound 1 and contained five individuals. Stratigraphic and material evidence, along with subsequent radiocarbon dating via DNA studies, confirms primary use during the Early Bronze Age (ca. 3300 BCE) with reuse in the Late Bronze Age, and later post-Roman or medieval intrusions. Twelve burials at the base of mound 1 were dated to the Early Bronze Age. One burial (#13) contained a Bronze Age ceramic and silver pendant while another (#7) contained a ceramic jug dating to EBA III (neither yielded viable DNA). Others were dated to EBA because of their location near or below #13 at the very centre of the mound in a rock cut grave (19, 20, and 21) or in nearby stone-lined pits (#13 and 17) and crouched position on side or back, covered by ochre (#18, 12, 9, and 7). Some crouched burials did not contain any ochre (#2 and 8) and are differentiated from other crouched burials (#4, 5, 6 , 10, and 15) only by darker grey-brown soil. Two of these burials (#5 and 15) contained LBA ceramics. Two supine, extended burials (#3, 14) were discovered near the surface of the mound and provisionally dated on the basis of the location and supine position to the post-Roman period. None of these late burials had any grave goods associated with them so the specific period is difficult to determine.

- **BOY008 (Mound 1, Grave 9):** *S. enterica* was recovered from an individual identified as male aged ca. between 60–65 years at death. The burial contained no grave goods, but the body was covered in ochre (consistent with ritual practices). Genetic analysis also associated the individual with populations of Pontic-Caspian steppe origin[1]. While the burial context suggests local continuity, the genetic data point to broader patterns of mobility.

##### 1.4. Binipati Nou: BPN

*Country:* Spain
*Region:* Tudons, Illes Balears
*Coordinates:* 39.9928°, 3.8906°
*Sample Date (BPN005):* Prototalaiotic, Iron Age, ca. 906–813 cal BCE (2 σ)
*Radiocarbon Date (BPN005):* 2716 ± 21 BP, 906–813 cal BCE (2 σ)
*Excavation/Sample Provenance:* Excavated between 1982 and 1983.
*Contact People:* Marta Díaz-Zorita Bonilla; Elena Sintes; Roberto Risch

*Site Description:*
The site is located on the western side of Menorca and is a Naveta with a boat-shaped ground plan. Radiocarbon analysis of human remains yielded ten dates, indicating that inhumations took place between approximately 1050 and 850 BCE, covering nearly the entire Prototalaiotic period. The site was used for collective burial, with the skulls and long bones of the deceased placed along the chamber walls, likely to make room for subsequent interments. Excavations conducted between 1982 and 1983 revealed the remains of around 50 individuals. Grave goods included typical Prototalaiotic pottery, bronze bracelets, bi-conical beads, and various ornaments, as well as bronze awls with bone handles and a bone button [2]**.**

- **BPN005 (MEN-125**): *S. enterica* was recovered from an infant individual (infant I, ca. 2 years old -1.30/5.8 year range).

##### 1.5. Brücken: BRC

*Country:* Germany
*Region:* Mansfeld-Südharz district, Saxony-Anhalt
*Coordinates:* 51.4442°, 11.1972°
*Sample Date (BRC017, BRC018, BRC034):* Early Medieval period, ca. 400–600 CE
*Radiocarbon Date:*

- *BRC017:* none, due to poor collagen preservation
- *BRC018: MAMS-63501:* 1616 ± 17 BP, 414–537 cal CE
- *BRC034: MAMS-63473:* 1577 ± 16 BP, 436–542 cal CE

*Excavation/Sample Provenance:* Excavated and documented in 2020 by Thomas Kubenz before the start of construction work.
*Contact People:* Jörg Orschiedt; Thomas Kubenz; Hellen Mager; Arnold Muhl

*Site Description:*
The early medieval burial ground at Brücken, located in the Mansfeld-Südharz district of Saxony-Anhalt, was fully excavated in 2020 in advance of planned construction. Situated approximately 1.3 km south of the modern village, the site covered 6525 $m^2$ and contained 76 graves with 77 individuals of varying ages and sexes. A nearby sacrificial pit held the remains of five horses, four cattle, and two dogs (mostly young males), analysed by Carola Oelschlägel.

The size of the cemetery is typical for early medieval Thuringian burial grounds, which rarely exceed 100 graves. Among the roughly 60 comparable sites in Saxony-Anhalt, Brücken is exceptional in having been fully excavated and comprehensively documented. This makes it an important dataset for demographic studies and for correlating typological and stylistic analyses with genetic and isotopic data on ancestry and mobility.

Grave goods, ranging from weapons and jewellery to simple tools (or, occasionally, their absence), reflect social differentiation and suggest both local Thuringian traditions and broader cultural contacts. Frankish tableware and Alamannic jewellery, for example, indicate links to the Rhineland and southwestern Germany. These finds suggest the cemetery was associated with a nearby manorial settlement.

Although the settlement has not yet been identified, it is presumed to lie within 500–1000 m of the cemetery, based on landscape analysis and regional settlement patterns. Historical sources from the Carolingian and Ottonian periods note the convergence of long-distance trade routes near Brücken, likely contributing to the site's regional importance and access to imported goods.

Based on preliminary results and grave good typology, the cemetery dates to ca. 460–540 CE, within the Germanic-Thuringian period. Skeletal preservation ranged from poor to moderate, limiting the accuracy of age and sex determinations. The population comprised approximately 29% subadults, 35% younger adults (20–40 years), and 36% older adults (40–60 years). Morphological and genetic data identified 32% of individuals as female and 29% as male. Dental pathologies—including calculus, caries, and periodontal disease—were present in over half the sample. Degenerative spinal changes affected 29% of older adults. Healed fractures were identified in 5% of postcranial bones and 2.5% of skulls. A unique case of intentional cranial modification was observed in a 45–55-year-old male—the only known male example of this practice in central Germany.

*S. enterica* was recovered from the three individuals **BRC017 (Ind. 13913:87:1)**, **BRC018 (Ind. 13913:50:1)** and **BRC034 (Ind. 13913:21:1)**.

- **BRC017** is a female adolescent individual, estimated to have died between 15–18 years.
- **BRC018** is a male child, approximately 6 ± 2 years old.
- **BRC034** is a female juvenile individual, approximately 11 ± 1.5 years old. Pathological examinations showed mild *Cribra orbitalia.*

#### 1.6. Chesh-Tyube: CHB

*Country:* Kyrgyzstan
*Region:* Talas Valley
*Coordinates:* 42.5184°, 72.2428°
*Sample Date (CHB002):* 725–404 cal BCE (2 σ)
*Radiocarbon Date (CHB002, tooth, MAMS-52916):* 2416 ± 21 BP, 725–404 cal BCE (2 σ).
*Excavation/Sample Provenance:* Site studied by I. K. Kozhomberdiev in 1960.
*Contact People:* A. Buzhilova; N. Berezina; L. Djansugurova; L. Musralina; E. Khussainova

*Site Description:*
The burial site is located in the Chesh-Tyube region on the left bank of the Kaindy River in the Manas region of the Talas district, Kyrgyzstan. The site was studied in 1960 by I.K. Kozhomberdiev[3].

The burial site consists of over 20 stone-earth mounds, eight of which were excavated. Six of them contained ground burials from the late Saka period, and the other two had catacomb burials of the Kenkol type. The first type of burial can currently be dated to the 4th–2nd centuries BCE, while the second type dates to the 3rd–5th centuries CE.

The grave goods for both types of burials were relatively poor and mainly consisted of ceramic ware.

- **CHB002***: S. enterica* was recovered from the lower right M3 of individual CHB002 (#622).

#### 1.7. Chleby: CHL

*Country:* Czechia
*Region:* Bohemia, Nymburk district
*Coordinates:* 50.2228°, 15.0894°
*Sample Date (CHL003)*: Bronze Age, Únětice culture (archaeological context), 2221–1895 cal BCE (2σ) (radiocarbon date)
*Radiocarbon Date (CHL003, pars petrosa, MAMS-40617):* 3623 ± 28 BP, 2221–1895 cal BCE (2σ)
*Excavation/Sample Provenance:* Excavated by the University of West Bohemia in 2016 (P. Krištuf).
*Contact Person:* Michal Ernée

*Site Description:*
In 2016, the University of West Bohemia in Pilsen conducted archaeological excavations at an Early Neolithic enclosure in Chleby, unearthing a multiple burial associated with the early stage of the Únětice culture. Chleby is located in the Nymburk district approximately 45 km east of Prague and 25 km north of Kolín.

Initially, a single child burial (skeleton 2035) was discovered and attributed to the Early Bronze Age Únetice culture. Subsequently, a burial pit (Ft 2036) containing skeletal remains of at least 15 individuals was uncovered. Within multiple stratigraphic layers, three skeletons in anatomical position, three sets of postcranial remains, four isolated skulls and a bone deposit were identified in the northern section of the burial pit.

Osteological analysis identified four female and seven male individuals. The age distribution primarily comprised adults (n=9). Additionally, two children, one juvenile, and three older adults were recorded. The individuals were most likely buried sequentially. As grave goods five vessels and animal bones were found [4].

- **CHL003 (skeleton 2039) –** *S. enterica* was recovered from a genetically male adult (ca. between 20–25 years). The skeleton was almost completely preserved and found in an anatomically correct position on the western side of the pit. The individual was laid down in a crouched position on the right side with a north-south orientation and facing east. The burial method as well as the radiocarbon dates indicate the early to middle Únětice culture.

#### 1.8. Copa: CPA

*Country:* Kazakhstan
*Region:* Karaganda region, Karkaraly district, Nurken village (Central Kazakhstan)
*Coordinates:* 43.5259°, 75.7886°
*Sample Date (CPA002):* Early Iron Age, Tasmola culture ($8^{th}$–$4^{th}$ centuries BCE)
*Radiocarbon Date (CPA002, tooth, MAMS-49763):* 2574 ± 21 BP, 796–567 cal BCE (AMS, 2 σ)
*Contact People:* A. Beisenov; L. Djansugurova; L. Musralina; E. Khussainova

*Site Description:*
Two burials were examined from the Copa site - a male individual from Kurgan 3 and a female individual from Kurgan 4. There is no evidence of grave goods or grave architecture.

- **CPA002 (Copa 4)** – *S. enterica* was recovered from a female individual, aged 25–35 years. The skull is brachycranial with a small longitudinal diameter and average width of the brain box. The frontal bone is broad, with both minimum and maximum widths classified as large to very large. The occipital bone is notably narrow, and the cranial base exhibits average dimensions. The facial skeleton is of medium vertical height and mesognathous in profile, with a combination of large facial angles and moderate alveolar projection. The zygomatic and facial widths are within the upper range on a global scale. The nose is wide and low (*platyrrhine*), with nasal bones moderately protruding. The orbits are wide and of average height, classified as mesoconchal. Facial flattening is observed in the upper tier, with moderate relief at the zygo-maxillary level. The *canine fossa* is minimally expressed.

#### 1.9. Fischamend: FSM

*Country:* Austria
*Region:* Bruck an der Leitha, Lower Austria
*Coordinates:* 48.1156°, 16.61278°
*Sample Date (FSM027):* ca. 1300–1600 CE
*Radiocarbon Date:* none
*Excavation/Sample Provenance:* Excavated in 2020 by the Novetus GmbH under direction of M. Binder.
*Contact People:* Kurt Alt; Nicole Nicklisch; Michaela Binder

*Site Description:*
Fischamend is located approximately 15 km south-east of Vienna in the district of Bruck an der Leitha (Lower Austria) and lies at the confluence of the Fischa and Danube rivers. In 2020 Novetus GmbH (directed by M. Binder) carried out archaeological excavations, uncovering the remains of the parish church of St. Stephan together with the cemetery. Archaeological evidence dates the church's construction in the late 12th century and its abandonment likely during the 17th century, possibly linked to the Thirty Years' War[5].

A total of 153 individuals were recovered from simple rectangular graves, often overlapping. Burials span the entire period of the church's use. Grave goods were rare (only present in 7.8% of the burials) and are interpreted as chronological markers rather than indicators of social status, consistent with medieval Christian burial practices. The buried individuals were primarily members of the local community. During the medieval and early modern periods, the economy of Fischamed was largely driven by agriculture and viticulture, supplemented by trade and transport, owing to the town's position along a major route connecting Vienna with Hungary and the eastern Habsburg territories [5].

- **FSM027 (Ind. 138, Obj. 163A, SE 411):** *S. enterica* was recovered from a male individual between 25–35 years of age at death. Only partial skeletal remains have been preserved, including the skull, cervical vertebrae, first thoracic vertebra, several ribs, the right shoulder girdle, and the right humerus. Despite the incomplete preservation the bone condition is excellent. The individual exhibited severe dental caries and degenerative changes in the cervical facet joints, likely resulting from repetitive physical stress.

##### 1.10. Grigorievka: GRY
*Country:* Kazakhstan
*Region:* Pavlodar
*Coordinates:* 52.7544°, 78.2095°
*Sample Date (GRY001, GRY004):* Bronze Age, Andronovo culture, ca. 1610–1510 BCE
*Radiocarbon Date:*
- *GRY001 (bone, MAMS-68688):* 3288 ± 19 BP, 1613–1507 cal BCE (2 σ)
- *GRY004 (bone, MAMS-68689)*: 3293 ± 19 BP, 1613–1509 cal BCE (2 σ)

*Excavation/Sample Provenance:* The burials were discovered during pipeline construction in 2020
*Contact People:* I. Merz; V. Merz; E. Usmanova

*Site Description:*
The burial ground is located in the village of Zhanakala (formerly, the village of Grigoryevka) on Pervogo Maya Street in the Pavlodar district, Kazakhstan. Zhanakala is located on the right valley side of the Irtysh River. During pipeline construction on April 19, 2020, two inhumation graves- provisionally designated as Grave 1 and Grave 2- were destroyed by an excavator. Within the trench, the edge of Grave 2 was partially visible. It was oriented west-east and measured approximately 1.2m.

The pottery assemblage of the burials is attributed to the late period of the Andronovo cultural and historical community. Radiocarbon dates indicate that the burials date from the end of the 17th to end of the 16th centuries BC.

- **GRY001 (Grave 1)**: *S. enterica* was recovered from an adult individual. The grave (Grave 1) was a single-inhumation containing the skeleton and a damaged ceramic vessel. Direct similarities to the decor of the vessel are not yet known, but each of the ornamental patterns is widespread in the Andronovo ornamental tradition.
- **GRY004 (Grave 2, Ind. 3)**: *S. enterica* was recovered from individual 3. This grave was located 15 m north from grave 1. From the grave filling, scattered skeletal remains from three adult individuals were collected, which, due to the high degree of destruction, are considered to represent a single burial. Eight fragments of pottery from four vessels were also found within the grave.
    - Vessel 1 (five fragments) is reconstructed as a jar-pot container. The vessel belongs to the Bishkul type of pottery of the Fedorovo Culture[6].
    - Vessel 2 is represented by a large fragment of the pot wall. The pot belongs to the Fedorovo type[6], which is found in the Kuznetsk basin, Tomsk Ob region, Baraba, Eastern and Central Kazakhstan, southern Trans-Urals, and the forest-steppe Tobol region[7].
    - Vessel 3 is represented by a fragment of the body of a pot. In terms of morphological features, it is close to item #1 from grave 289 of the Yelovsky complex-II. In general, pottery of this type is found in the Andronovo complexes of the Middle Yenisei, Kuznetsk Basin, Baraba forest-steppe, and Tomsk Ob region [7].
    - Vessel 4 is represented by a fragment of rim from a jar-type vessel. The ornamental pattern on the pottery is known mainly on the eastern periphery of the Andronovo world[7].

##### 1.11. Gars-Thunau: GTH

*Country:* Austria
*Region:* Gars am Kamp, district Horn, Lower Austria
*Coordinates:* 48.5871°, 15.6454°
*Sample Date (GTH003):* ca. 800–1000 CE (contextual date), Early Medieval
*Radiocarbon Date:* none
*Excavation/Sample Provenance:* Excavated in 1987 by the Institute for Prehistoric and Early Historic Archaeology at the University of Vienna.
*Contact People:* Doris Pany-Kucera; Margit Berner; Sabine Eggers

*Site Description:*
The early medieval cemetery at Thunau am Kamp, specifically on the Obere Holzwiese, is located in Lower Austria, north of the Danube. The site is part of a larger fortified hill settlement on the Schanzberg, which is unique in Austrian early medieval archaeology due to the size and the scope of the investigation. The graves were excavated primarily in 1987 and 1990 as part of a research project by the Institute for Prehistoric and Early Historic Archaeology at the University of Vienna[8,9].
The cemetery on the Obere Holzwiese is the largest early medieval burial ground north of the Danube in Lower Austria. It dates mainly to the Carolingian period ($8^{th}$–$9^{th}$ century) and reflects Christian burial practices such as extended supine inhumations in simple earth graves, with

occasional use of wooden coffins and stone structures. Some graves also included food offerings and material goods, such as dress items and ceramics, indicating complex social identities and burial customs.

Regarding the skeletal remains, two PhD theses were carried out, one on entheseal and joint changes and one on dietary reconstruction by stable isotope analysis.[10,11] Selected studies on the site have been published so far, e.g. on the distribution of tuberculosis among the population[12], and a case of malaria was recently described[13].

In the course of the HistoGenes project (Austrian academy of Sciences, HistoGenes project, n° 856453 ERC-2019-SyG), in total, 48 individuals (21 subadults, 27 adult individuals) from a middle, contiguous area of the Thunau cemetery were selected for aDNA sampling. Based on radiocarbon dates from some of those, the time frame covers ca. 770–990 CE[8].

- **GHT003 (NHM-OSTE 25019, Grave 64)**: *S. enterica* was recovered from a morphologically undetermined, genetically female child with an estimated age at death of 9–10 years[11]. The skeletal remains are partly preserved and the child was buried in supine position without grave goods. The cranial remains of the skeleton from grave 64 show endocranial lesions at the internal lamina (single and fine-branched vessel impressions and new bone formation), which are partly well organised. Moreover, there are porosities along the sagittal sinus, and enhanced depressions are visible left and right of the middle part. The individual shows slight orbital cribra, and porotic hyperostosis. The mandible displays porosities at the maxillar and mandibular alveolar ridge, as well as porosities along the mandibular foramen bilaterally.

  The postcranial remains reveal areas of whitish porous new bone formation, probably in remodelling status, which are especially visible at the femora. Remnants of these changes are present at the right medial ulnar shaft (*facies anterior*), and extensive distribution of new bone formation is visible medial of the *linea aspera*, mainly upper two thirds of the femur shafts and also at the dorsolateral aspect. The right tibia exhibits this new bone formation at the *facies posterior* and *medialis* (predominantly distal), and the left tibia demonstrates similar characteristics at these locations, albeit to a lesser extent. The left fibula displays new bone formation at the *facies lateralis* (the right fibula is not preserved). Porosities that may be increased are visible at the ventral aspect of two thoracic vertebrae. These skeletal lesions may indicate scurvy and/or anaemia.

##### 1.12. Hulín 1 - U Isidorka: H1I

*Country:* Czechia
*Region:* Moravia, Kroměříž district
*Coordinates:* 49.3169°, 17.4649°
*Sample Date (H1l004):* Bronze Age, Bell Beaker culture, 2465–2287 cal BCE (2 σ)
*Radiocarbon Date (H1l004, tooth, MAMS-48832):* 3883 ± 25 BP, 2465–2287 cal BCE (2 σ)

*Excavation/Sample Provenance:* Hulín 1 – U Isidorka. Excavated by the Archaeological Center Olomouc (ACO) during extensive rescue excavation between 2004 and 2005.
*Contact Person:* Jaroslav Peška

*Site Description:*
Between 2004 and 2005 the Archaeological Center Olomouc excavated a ca. 14 ha area along the D1 highway in the Kroměříž district (Moravia). The Hulín 1 – U Isidorka site is a prehistoric multi-cultural settlement on an elevation between the Moštěnka and Rusava rivers. The excavation revealed settlement remains and burials with around 750 finds from the Late Neolithic and Early Bronze Age[14]. Finds include habitation traces from the Nitra, Bell Beaker, Únětice, and Věteřov cultures as well as necropolises linked to the Moravian Corded Ware and Nitra cultures. A Bell Beaker necropolis contained 29 graves with 30 inhumations and four cremations. The grave goods confirmed cultural affiliation, including ceramics, jewelry, tools, and weapons. Of the 20 excavated graves, the majority contained inhumations, with only four cases presenting cremated remains. These burials were aligned along a northeast-southwest orientation. The individuals were typically deposited in a contracted posture, predominantly positioned laterally – eleven on the left side, and eight on the right. Eight of the interments yielded well-preserved skeletal material, enabling both sampling procedures and detailed bioarchaeological investigation[15].

- **H1l004 (Grave H 73)** – *S. enterica* was recovered from a genetically male individual between ca. 35–45 years (*maturus*). *S. enterica* was recovered from a tooth. The individual was buried in a crouched position on the left side, head towards north. Grave H 73 was the largest grave of the Bell Beaker cemetery (2.35 m x 1.71 m x 0.71 m) and contained a complete set of blacksmithing equipment, including anvils and hammers.

##### 1.13. Hauts de Lattes: HDL
*Country:* France
*Region:* Montpellier (Hérault), Occitanie
*Coordinates:* 43.57°, 3.9094°
*Sample Date (HDL004):* 671–820 cal CE (2 σ).
*Radiocarbon Date (HDL004, bone, MAMS-48716):* 1267 ± 20 BP, 671–820 cal CE (2 σ)
*Excavation/Sample Provenance:* The site was excavated in 2018 as part of rescue excavations directed by I. Daveau (Inrap).
*Contact Person:* Mathieu Ott

*Site Description:*
The Soriech farm complex was situated within an L-shaped walled enclosure, measuring ca. 316 $m^2$. It was located on the eastern slope of the Pérols hill, providing a strategic vantage point over the Lez valley and an ancient route connecting the trading port and bishopric of Maguelone on the Mediterranean coastline to the historic settlement of *Sextantio* (modern-day Castelnau-le-Lez). Archeological evidence dates its main occupation between ca. 650 and 750 CE. To the southern perimeter of the enclosure, a complex system of intersecting walls formed tight corridors and angular passageways. These features may have had a defensive function, restricting access or controlling movement within the site.

Two silos (SI20242, SI21752) were excavated in the southeast corner of the courtyard. Stratigraphic analysis indicates that both were dug and backfilled simultaneously, when the earthen wall separating them collapsed. The use of the twin silos changed at this stage, with the deposition of a body (SP21789) across the half-filled pits.

- **HDL004 (SP21789)** – *S. enterica* subsp. *enterica* was recovered from a mature adult. The remains of SP21789 were deposited directly across both silos, with the upper body in silo SI21762 and the lower limbs extending into silo SI20242. This level is not horizontal, and varies from 9.54 m above sea level under the spine to 9.90 m above sea level under the tibias.

  The body was laid on an irregular and disorganised layer of limestone blocks whose edges protruded into the body. In the north-eastern part of the structure, there was a large slab stuck obliquely into the sediment of the silo, which could initially have been used to seal it. This cover seems to have protected the individual's skull, but we remain cautious about this hypothesis, as the presence of the slab could be a coincidence linked to the final filling of the structure.

  The skeletal remains were poorly preserved, as mechanical excavations during the initial silo exposure destroyed a significant part of the body. Contact with limestone appears to have accelerated bone decay, turning much of the remains to powder.
  The individual was buried supine, with their head facing north and turned leftwards. The upper limbs were tightly flexed inward, and the knees were elevated, likely due to the uneven grave surface. Post-mortem joint disarticulation suggests the body decomposed in an empty space, likely within a biodegradable container (e.g. a cloth shroud or other soft wrapping), as no rigid container could fit the irregular pit. The skeletal analysis revealed degenerative joint disease and *cribra orbitalia* in both eye sockets.

##### 1.14. Hoštice 1: HO1

*Country:* Czechia
*Region:* Moravia
*Coordinates:* 49.5218°, 17.1585°
*Sample Date (HO1003):* Bronze Age, Bell Beaker culture, ca. 2465–2299 cal BCE (2 σ)
*Radiocarbon Date (HO1003, tooth, MAMS-48834):* 3898 ± 20 BP, 2465–2299 cal BCE (2 σ)
*Excavation/Sample Provenance:* Hoštice 1 Za Hanou. Excavated by the Archaeological Rescue Research in Brno under the direction of A. Matějíčková between 2002 and 2003 during extensive rescue excavations.
*Contact Person:* Jaroslav Peška

*Site Description:*
The Institute of Archaeological Rescue Research in Brno conducted extensive rescue excavations between the years 2002 and 2003 along the D1 highway between Vyškov and Mořice under the direction of A. Matějíčková. Excavated was a 1 ha area, uncovering 257 graves containing 143 individuals from the Bell Beaker culture[16,17]. Most individuals were interred in the typical

gender-specific flexed position, with exceptions including a multiple burial and four cremations in urns. Radiocarbon dating indicated the necropolis was used between 2480–2130 BCE, aligning with the entire regional Bell Beaker phase. The grave goods reflect this extended use period. Anthropological analysis revealed an equal male-to-female ratio, with approximately 40% of the individuals being children or adolescents[16,17].

- **HO1003 (Grave 68, Ind. H 900)** – *S. enterica* was recovered from an adult individual aged ca. 40–50 years. The osteological investigations indicated a female individual, whereas genetic testing identified a male sex. The skeleton was found in a right-sided flexed position, oriented south-north with the face directed eastward. The skull, right arm, and several smaller bones were displaced from their anatomical position.

##### 1.15. Ivanovice VI-Borůvka: IVB

*Country:* Czechia
*Region:* Moravia
*Coordinates:* 49.3055°, 17.0933°
*Sample Date (IVB002):* Bronze Age, Bell Beaker culture, ca. 2288–2141 BCE (2 σ)
*Radiocarbon Date (IVB002, tooth, MAMS-48833):* 3783 ± 21 BP, 2288–2141 cal BCE (2 σ)
*Excavation/Sample Provenance:* Ivanovice VI – Borůvka. Excavated between 2002 and 2003 as archaeological rescue excavations by the Institute for Archaeological Heritage Protection.
*Contact Person:* Jaroslav Peška

*Site Description:*
Between 2002 and 2003, rescue excavations were conducted in the Ivanovice na Hané area by the Institute for Archaeological Heritage Protection, under the direction of Bálek, Vitulová, Smíd, and Parma. Additional investigations followed in 2006 and between 2014 and 2016.

The earliest part of the site, Ivanovice na Hané 6 "Borůvka", comprises a Bell Beaker cemetery. Adjacent is Ivanovice na Hané 7 "Spravedlnost", which yielded finds from the Corded Ware culture, Bell Beaker culture, and both the Early and Late Bronze Age[18].

In Ivanovice 6, a 35 x 45 m cemetery with 23 graves, 22 skeletons, and one biritual burial (including an urn) was documented. The graves are deep and mostly circular to oval in shape. Possible settlement traces, such as postholes and undated pits, were also observed.

Some of the largest graves (806, 807, 812, 814) may have been originally covered by burial mounds, based on their size and spacing (2–4 m apart). The skeletal remains are generally well-preserved, with only minor surface damage due to soil pressure. The burial practices conform to Bell Beaker gender norms, with males placed on the left side facing north, and females on the right-side facing south[18,19].

- **IVB002 (Grave VI–812/02)** – *S. enterica* was recovered from a male adult individual (age between 25–35 years). The osteological examinations of the skull and pelvis revealed the sex and age of the deceased. The individual from Grave VI–812/02 was well-preserved, with 31 teeth remaining and signs of cribra orbitalia in the left orbit[20]. A mandibular molar (M37) was sampled for analysis. Radiocarbon dating places the burial between 2288–2141 cal BCE, aligning with the Bell Beaker period.

The skeleton lay in a flexed position on the left side, oriented northeast southwest. The burial was notably rich, potentially marked by a barrow and wooden enclosures. Grave goods included two copper daggers, three ceramic vessels, and a set of arrows behind the back.

##### 1.16. Koken: KKN

*Country:* Kazakhstan
*Region:* Abay Oblast
*Coordinates:* 49.8643°, 79.6001°
*Sample Date:* Bronze Age, 1875–1637 cal BCE (2 σ)
*Radiocarbon Date (KKN099):* 3436 ± 23 BP, 1875–1637 cal BCE (2 σ)
*Excavation/Sample Provenance:* Archaeological excavations began in 2019 by the Trans-Eurasian Exchanges: Contemporary Dialogues and Archaeological Inquiry (TEECA) Project.
*Contact People:* Paula Doumani Dupuy; Aidyn Zhuniskhanov

*Site Description:*
The site Koken is located in eastern Kazakhstan in the Kokentau Mountains, south of the present-day city of Semey. The area lies at the intersection of the Eurasian Steppe and the Inner Asian Mountain Corridor, a transitional zone in the prehistoric cultural landscape. The site is positioned along the banks of seasonal marshland and includes both a settlement and a cemetery, surrounded by steppe, mineral outcrops, and rich biodiversity. Radiocarbon dates from the site reveal intensive use during the entire Bronze Age. The cemetery yielded graves dated between ca. 1900–1400 cal BCE in the Middle Bronze Age, whereas the settlement layers show occupation spanning from 2840–906 cal BCE (Early to Late Bronze Age), based on AMS dating of charcoal, human bones, and faunal bones.

Excavations were conducted under the Trans-Eurasian Exchanges: Contemporary Dialogues and Archaeological Inquiry (TEECA) Project, a collaboration between Nazarbayev University (KZ) and the Margulan Institute of Archaeology in Almaty. Systematic surveys of the Kokentau Mountains in 2018 identified over 30 archaeological sites in the vicinity, including the site of Koken. Excavations at Koken then commenced in 2019. Approximately 70 graves were identified, of which 22 have since been excavated. Excavations of the settlement area have spanned over 200 $m^2$ revealing multiple building phases spanning the Stone Age to Ethnographic period[21].

- **KKN099 (KKBR19)** – *S. enterica* was recovered from an adult female individual between 50–70 years old at death. The individual was interred in a Middle Bronze Age grave cluster at the southern end of the cemetery. The burial had been looted and the skeletal remains were disarticulated, with no associated grave goods. Osteological analysis revealed signs of dental disease, vertebral and joint degeneration, as well as musculoskeletal stress markers. Lambdoid ossicles were noted bilaterally.

##### 1.17. Kamenice: KNC

*Country:* Albania
*Region:* Korça
*Coordinates:* 40.5329°, 20.7255°
*Sample Date:* Bronze Age to Iron Age

- *KNC041:* Iron Age, 767–543 cal BCE (2 σ)

- *KNC056:* Iron Age, 900–808 cal BCE (2 σ)
- *KNC095:* Iron Age, 767–541 cal BCE (2 σ)
- *KNC103:* Iron Age, ca.  850–750 BCE (contextual)
- *KNC108:* Iron Age, ca. 850–750 BCE (contextual)
- *KNC139:* Iron Age, 754–421 cal BCE (2 σ)
- *KNC181:* Late Bronze Age/Early Iron Age, 1108–929 cal BCE (2 σ)

*Radiocarbon Date:*

- *KNC041:* 2490 ± 18 BP, 767–543 cal BCE (2 σ)
- *KNC056:* 2699 ± 18 BP, 900–808 cal BCE (2 σ)
- *KNC095:* 2484 ± 18 BP, 767–541 cal BCE (2 σ)
- *KNC139:* 2460 ± 18 BP, 754–421 cal BCE (2 σ)
- *KNC181:* 2851 ± 19 BP, 1108–929 cal BCE (2 σ)

*Excavation/Sample Provenance:* Excavated between 2000 and 2007.
*Contact Person:* Lorenc Bejko

*Site Description:*
The Kamenicë tumulus is located in south-eastern Albania, on the southern edge of the Korça Basin - a well-researched region that has contributed significantly to the reconstruction of the prehistoric development of Albania and the Western Balkans. The tumulus was originally part of a larger group of burial mounds, many of which have been completely destroyed in the last 80 years due to intensive agricultural use.

Systematic excavations were carried out between 2000 and 2007 following repeated looting in the late 1990s. 401 burials with a total of 425 individuals have been documented. Only two thirds of the mound has been excavated (one third is preserved in situ) and an unknown part has been damaged by looting; it is estimated that the tumulus originally contained around 600 burials.

The earliest burial - a central grave - dates to the beginning of the 16$^{th}$ century BCE (Middle Bronze Age according to local chronology). The original cemetery was surrounded by two concentric stone circles of medium size, with the first grave in the centre. Over a period of almost seven centuries, the tumulus developed into an artificial, hemispherical mound with a diameter of approximately 26 m and a maximum height of 3.1 m. Each new burial led to the expansion of the mound through fresh earth mounds, occasionally through small to medium-sized stones and often through wooden structures.

In the second half of the 8$^{th}$ century BCE, there is evidence of a transformation: the tumulus was remodelled into a large, elongated cemetery about 70 m long and 40 m wide. During this phase, only stones of different sizes were used to construct the new section, which surrounded the original mound, particularly on its southern and eastern sides.

Over 270 burials were documented during the first utilisation phase (16$^{th}$ to mid-8$^{th}$ century BCE). These are unevenly distributed across the Middle and Late Bronze Age, with the majority falling into the Early Iron Age (10th to mid-8th century BCE). The second phase comprises 131 burials; however, only around half of this section has been excavated, with the remainder preserved for future research.

The two phases differ not only in the structure of the tumuli, but also in terms of burial structures, spatial organisation and the material culture of the grave goods. The last burials date to the end of the 6th century BCE.

- **KNC041 (AL-KAM-041-V140, Grave 140)** – *S. enterica* was recovered from a young adult female individual, dated to the second half of the 8$^{th}$ century BCE (radiocarbon date: 767–543 cal BCE). The burial is attached to a stone lined grave of an adult male. The female burial KNC041 was interred with seven grave goods. The assemblage includes a pair of gold earrings or head ornament (found in only 18% of contemporary female burials) and a bronze button (unique find for females in this period), the remaining items correspond to typical indicators of female identity at Kamenicë.
- **KNC056 (AL-KAM-056-V028, Grave 28)** – *S. enterica* was recovered from a middle aged, adult male individual. It is radiocarbon dated to between 900–808 BCE (Early Iron Age II) and belongs to the first phase of the site. The grave belongs to the more richly equipped burials among the male interments of the Early Iron Age II period with nearly four times the average quantity of grave goods. Grave goods contain a.o. matt-painted cantharos, an iron spearhead and knife – these are typically associated with male identity during the Iron Age at Kamenicë and in the wider region.
- **KNC095 (KAM-102-V141, Grave 141)** – *S. enterica* was recovered from a young female adult individual, dated to the second half of the 8$^{th}$ century BCE. The burial is attached to the stone-lined grave of an adult male. The burial contains seven grave good items.
- **KNC103 (KAM-110-V192, Grave 192)** – *S. enterica* was recovered from an adult female individual. The burial is a stone-lined grave covered with mid- and small-sized stones. It is spatially differentiated from the cluster of KNC041 and KNC095. There is no radiocarbon dating of the individual, but the stratigraphy indicates the time period of the late 8$^{th}$ century BCE. Nine grave goods were found – pottery, ornaments, cloth accessories, and a tool associated with female identity during the second phase of the site. Worth noting is a bronze applique (decorating a belt buckle of some sort) which compares with similar objects found in the western Balkans in similar burial contexts (Glasinac, in today's Bosnia-Herzegovina). This is the only such item from Kamenicë.
- **KNC108 (KAM-115-V215, Grave 215)** – *S. enterica* was recovered from an adult female individual. The burial is located in a stone-lined grave next to an adult male (Grave 182). Like KNC103 no radiocarbon date is available, but it can securely be dated to the beginning of the second phase of the site (late 8$^{th}$ to early 7$^{th}$ centuries BCE); the date is based on stratigraphy and artefacts analysis. Five artefacts are associated with this burial, all characteristic of female identity during the site's second phase. These include a matt-painted amphoriskos, iron tools, a dress pin, and a single gold earring—the latter linking this individual to KNC041. Notably, female individuals KNC041, KNC095, KNC103, and KNC108 show similarities in age, date, and grave good composition. Each burial includes amphoriskoi, iron pins, and tools, differing primarily in the type of jewelry or ornaments.
- **KNC139 (KAM-146-V6)** – *S. enterica* was recovered from a 25–30-year-old male individual. The remains of the individual were discovered outside of its primary context, removed by looting activities on the site. Radiocarbon dating of a bone dates the individual to the second phase of the development of the site. No grave goods can be linked to the individual.

- **KNC181 (KAM-182-V179, Grave 179)** – *S. enterica* was recovered from an adult male individual and dates to the Late Bronze Age and Early Iron Age I (first phase of the site). The grave consists of thick layers of decayed wood above and under the skeleton, indicating the use of a well-constructed wooden coffin. No grave goods were found in the burial.

##### 1.18. Kenkolsky burial site: KNL

*Country:* Kyrgyzstan
*Region:* Talas Valley
*Coordinates:* 42.51837°, 72.2429°
*Sample Date (KNL019, KNL025, KNL026):* Iron Age (Central Asia)
*Radiocarbon Date:*

- *KNL019 (tooth, MAMS-52900):* 1824 ± 19 BP, 132–318 cal CE (2 σ)
- *KNL025 (tooth, MAMS-52895):* 1712 ± 19 BP, 257–408 cal CE (2 σ)
- *KNL026 (tooth, MAMS-49765):* 1851 ± 22 BP, 128–238 cal CE (2 σ)

*Excavation/Sample Provenance:* Excavation between 1938–1939 by A. N. Bernsham, between 1956–1957 and 1960 by I. K. Kozhomberdiev and between 2001–2002 by B. E. Amanbaeva.
*Contact People:* A. Buzhilova; N. Berezina; L. Djansugurova; L. Musralina; E. Khussainova

*Site Description:*
The burial site is located on the left bank of the Kenkol River, near the village of Tash-Aryk in the Talas region of Kyrgyzstan. It was introduced into scientific circulation by A.N. Bernsham, who first conducted excavations there in 1938-1939, followed by further research in 1956-1957 and 1960 by I.K. Kozhomberdiev, and again in 2001-2002 by B.E. Amanbaeva.

The Kenkol burial site contains about 100 kurgans, scattered in several groups without any systematic arrangement. Approximately 60 kurgans have been excavated. Most of the structures are earth mounds containing burials in catacombs, predominantly dating to the 3$^{rd}$–5$^{th}$ centuries BCE. There are also a small number of stone-earth mounds with burials in ground pits, dating to the 5$^{th}$–2$^{nd}$ centuries BCE, located in the eastern part of the burial site. The skulls bear traces of ring-like deformation, which is linked to the area of the Aral Sea[22].

The catacombs in Kenkol had relatively long dromos (passageways) perpendicular to the burial chamber, often blocked with stone. The burial chambers were quite spacious. Beneath large burial mounds, there were usually single burials, often in wooden coffins. Small mounds contained catacomb burials of a collective nature, representing family tombs. The catacomb burials at the Kenkol site yielded rich grave goods. In male burials, bone fittings for bows of Hunnic type, iron arrowheads, fragments of swords and daggers, knives, iron bits, and buckles were found, along with wooden bow saddles, etc. In female burials, there were ornaments in a polychrome style, earrings, pins, mirrors, including Chinese imports, beads, cosmetic sticks made of antimony, and spindles. Items of clothing, including silk, were also of interest. Almost all burials contained ceramic ware, a significant portion of which was decorated with a characteristic carved wavy pattern. In addition, wooden tables, bowls, cups, cradle remnants, fragments of baskets, and more were found.

*S. enterica* was recovered from the teeth of 3 individuals:

- **KNL019 (Ind. 700/K34)**: Lower right M3
- **KNL025 (Ind. 706/K28)**: Upper left M2
- **KNL026 (Ind. 709/K19)**: Upper left M2

##### 1.19. Kolín, area I-7: KO1

*Country:* Czechia
*Region:* Central Bohemia
*Coordinates:* 50.0283°, 15.2078°
*Sample Date (KO1037):* Early Bronze Age, Úně̌tice culture, 1927–1767 cal BCE (2 σ)
*Radiocarbon Date (KO1037, CRL 21_864):* 3521 ± 19 BP, 1927–1767 calBCE (2 σ)
*Excavation/Sample Provenance:* Excavated during rescue excavations between 2008 and 2010 led by R. Šumberová.
*Contact Person:* Michal Ernée

*Site Description:*
Led by Šumberová, the excavations were carried out between 2008 and 2010, prior to the construction of a bypass road. Kolín, located ca. 50 km east of Prague and south of the Elbe River. Archaeological evidence points to continuous occupation from the Neolithic onward. Settlement and burial features represent a range of cultural groups, including the Funnel Beaker, Baden, Řivnáč, Globular Amphora, Únêtice, and Hallstatt cultures, as well as the Corded Ware culture and the Early Medieval period[23].
An Early Bronze Age Únětice culture inhumation cemetery with 70 burials in two groupings was excavated. The entire cemetery was analysed by researchers in various fields and published[24,25].

- **KO1037 (Grave 65)** – *S. enterica* was recovered from a female individual with an osteological age of 40 years or older. The rectangular single grave was located slightly away from the other graves, aligned southwest-northeast, and featured substantial stone lining along the north and south walls composed of large, flat slabs. The individual was buried in a flexed position on the right side, oriented southwest. The burial was secondarily disturbed, with skeletal elements displaced up to 50 cm from their original position. A mandibular molar (M37) was sampled and analysed. Radiocarbon dating places the burial between 1927–1767 cal BCE, corresponding to the classic phase of the Únětice culture. In addition, isolated foci of osteomyelitis (bone marrow inflammation) and inflammation of the fossa intercondylaris on the thigh bone were detected.

##### 1.20. Kiprino-II: KPI

*Country:* Russian Federation
*Region:* Shelabolikhinsky District, Altai Krai
*Coordinates:* 53.5339°, 82.0923°
*Sample Date:* Andronovo culture, Bronze Age, 16$^{th}$–14$^{th}$ centuries BCE
*Radiocarbon Date:* Not dated.
*Excavation/Sample Provenance:* Discovery of the burial ground in 1983 by Y. F. Kiryushin
*Contact People:* Alexey Tishkin; Y.F. Kiryushin

*Site Description:*
The Kiprino-II soil burial ground was discovered by Y.F. Kiryushin in 1983. It is located in Shelabolikhinsky District of Altai Krai (Russia), near the village of Kiprino. Burials of ancient individuals were discovered on sand dunes within the floodplain of the Ob River. Preliminary analysis indicates that the burial ground is associated with the Andronovo culture of the Bronze Age (16$^{th}$–14$^{th}$ centuries BCE), based on the orientation and positioning of the bodies and the presence of typical ceramic vessels.

- **KPI001 (Grave 2, Sample no. 4395)** – *S. enterica* was recovered from a male individual, who died between 20–30 years. The individual was buried in a hunched position, on the right side with his head to the south-west. According to the paleoanthropological analysis, this individual is close to the Andronovo variant of the Proto-European type, probably mixed with the local West Siberian ancient population (definitions of physical anthropologist Dr K.N. Solodovnikov). The sample was submitted to the Cabinet of Anthropology of Tomsk State University and examined by Professor A. A. Tishkin.

##### 1.21. Kara-Saz: KSZ

*Country:* Kyrgyzstan
*Region:* Ton District (Issyk-Kul)
*Coordinates:* 41.7492°, 76.6152°
*Sample Date (KSZ005):* 5$^{th}$ to 2$^{nd}$ centuries BCE (356–169 BCE radiocarbon date)
*Radiocarbon Date (KSZ005, tooth, MAMS-52620):* 2182 ± 18 BP, 356–169 cal BCE (2 σ)
*Excavation/Sample Provenance:* Archaeological research was conducted between 1953 and 1955 by A. K. Kibiriv.
*Contact People:* A. Buzhilova; N. Berezina; L. Djansugurova; L. Musralina; E. Khussainova

*Site Description:*
The Kara-Saz burial ground is situated near Kyzyl-Ompol Mountain, at the outlet of a small valley in southwestern Pre-Isyk-Kul, within the Ton district of Kyrgyzstan's Issyk-Kul region. The site was excavated between 1953 and 1955 under the direction of A. K. Kibiriv. The excavated cemetery comprises ca. 100 stone-and-earth mounds (kurgans), spanning multiple chronological phases. Most of the larger burials are associated with the Saka period (5$^{th}$–2$^{nd}$ centuries BCE). The smaller mounds, containing upright stone stelae, are dated to the Turkic period (6$^{th}$–early 10$^{th}$ centuries CE). Six of these mounds were excavated and contained inhumations in ground-level graves, mainly single burials and with the exception of one collective burial of three individuals. The deceased were buried in extended supine positions, the head oriented toward the west or northwest. The accompanying grave goods included hand-formed ceramic ware, iron knives, bronze earrings, rings, beads, stone spindle whorls, and more.

Stone-covered pit graves can be a characteristic feature of the Central Tien Shan region. Slab-roofed graves are emblematic of the pre-Sun (Iszedon) tombs, generally dated to the 4$^{th}$–3$^{rd}$ centuries BCE. Grave goods like gold-foil ornaments, iron knives, and textile-impressed ceramics show connections to the Usun phase of the Sak-Usun period (1$^{st}$ century BCE–1$^{st}$ century CE).

- **KSZ005:** *S. enterica* was recovered from the lower left M2 of KSZ005 (Kara-Saz 602)

##### 1.22. Majaky: MAJ

*Country:* Ukraine
*Region:* Oblast Odessa
*Coordinates:* 46.4122°, 30.2731°
*Sample Date (MAJ022):* Yamnaya culture, Eneolithic, 3329–3022 cal BCE (2 σ)
*Radiocarbon Date (MAJ022, tooth, MAMS-63966):* 4451 ± 17 BP, 3329–3022 cal BCE (2 σ)
*Excavation/Sample Provenance:* Excavation by the Odesa Archaeological Museum
*Contact People:* Igor Bruyako; Igor Manzura

*Site Description:*
The archaeological complex near the village of Mayaki is located on the left bank of the lower reaches of the Dniester River, at the southern outskirts of the village (Belyaevka district, Odessa region, Ukraine). It is located on the third river terrace. The southern ledge of this terrace, occupied by the settlement of the Usatovo culture and burials of later periods, is partially bounded by an ancient gully in the east and the floodplain of the Dniester River in the west and south. In plan, this ledge is a separate cape, noticeably towering above the floodplain. The main part of the archaeological complex is the settlement and cemetery of the Usatovo culture. In addition to the objects of this culture, materials of different historical periods have been discovered on the territory of the site. Over the years, Early Eneolithic, Bronze Age, and archaic "Greek-Scythian" burials, an extensive Sarmatian cemetery, and a field of ritual pits from the first centuries CE have been studied here.

In the general plan, the territory of the site can be inscribed in a conventionally delineated rectangle with dimensions of 600 × 200 m, stretched in the direction of southeast-northwest. The settlement of the Usatovo culture occupies a small outlying area of 130×40 m. About two-thirds of its area was destroyed by erosion processes. To date, only the northern part with an area of 0.52 hectares has been preserved. The studied remains of the settlement are a kind of labyrinth of six or seven deep ditches which are connected to each other and follow in different directions. According to the totality of features, this part of the archaeological complex is considered as a ritual centre, where various religious rites were periodically performed, and ceremonies and feasts were arranged within the framework of prestige-gift relations.

The cemetery associated with the settlement is 200 m to the north-west and occupies an area of about 300 × 100 m. The burial ground belongs to the mound-flat type, although due to many years of ploughing, it is not always possible to determine the real nature of some burial complexes. During excavations, 47 burials and several cult pits of the Usatovo culture were discovered. All primary graves in the barrows belong to the Usatovo culture. Secondary burials in the barrows belong to different periods of the Bronze Age starting from the Yamnaya culture and later. Barrow 1 with a height of 1 m and a diameter of 35–50 m was situated at the southern edge of the cemetery. It was explored in 1974. Primary burial 9 belongs to the Usatovo culture. Burials of the Yamnaya culture, the Late Bronze Age and later periods were allowed into the mound later. On the surface of the mound there were stones, probably from a destroyed cromlech. A ditch with a diameter of 35 m and a depth of 2.3 m was dug around the primary mound. The structures belong to the Usatovo culture.

- **MAJ022 (Kurgan 1, Grave 19)** – *S. enterica* was recovered from a male adult individual. The skeleton was laid in a crouched position on its back, with the head facing west. His arms were stretched out along the body, the legs bent, and the knees turned to the right. Grave 19 of the Yamnaya culture was inserted into the burial mound of the Usatovo culture. The burial pit had an irregular oval shape (1.7 × 1.1 × 0.95 m) and was covered with stone slabs.

##### 1.23. Maguelone: MGL

*Country:* France
*Region:* Villeneuve-lès-Maguelone (Hérault), Occitanie
*Coordinates:* 43.5332°, 3.8616°
*Sample Date:* 5th–8th century CE
*Radiocarbon Date (MGL008, bone, MAMS-55363):* 1280 ± 19 BP, 672–774 cal CE (2 σ)
*Radiocarbon Date (MGL011, tooth, MAMS-5509):* 1589 ± 19 BP, 427–542 cal CE (2 σ)
*Excavation/Sample Provenance:* Excavations and archaeological research since 1998 by the CNRS. Excavations in 2019 and 2020 by L. Tarrou (Inrap), C. Raynaud (Cnrs) and B. Ode (SRA Occitanie).
*Contact Person:* Claude Raynaud

*Site Description:*
Located on the Mediterranean coast, approximately ten kilometers south of Montpellier, the former islet of Maguelone served as the seat of a bishopric from the 6th to 16th centuries, as evidenced by the Romanesque cathedral built at its summit. Most of the remains unearthed during archaeological research conducted by the CNRS and Inrap since 1998 date to Late Antiquity and the Early Middle Ages. The predominance of amphorae highlights the scale of commercial activity along the coast, marked by abundant imports from across the Mediterranean.

From the 4th to 7th centuries, archaeological evidence spans almost the entire islet, covering approximately twenty hectares. Observations indicate a primary settlement at the summit, where numerous materials have been recorded, including but not limited to tiles, cut stones, marble, and mosaics. Elsewhere on the islet, concentrations of slag indicate a glass- and iron-ware production area. In the southwest, at a location known as “Port Sarrazin”, the abundance of amphora fragments suggests the existence of a port facility, where small boats likely transferred cargo after crossing the ancient Grau de Maguelone, a narrow channel now blocked in the coastal barrier.

During the same period, hastily buried bodies found outside the established burial grounds point to the presence of individuals marginalised by the community, likely reflecting the isolation of those suffering from contagious diseases.

- **MGL008 (MAG19, SP2559):** *S. enterica* was recovered from individual SP2559. The individual was placed directly into an unlined pit, whose contours were indistinct due to the identical nature of the fill and surrounding sediment. The position of the body could indicate that the pit was too small for the individual. The radiocarbon date suggests a deposition in a neglected area, possibly linked to the abandonment of nearby buildings.
  The body was tightly constrained, with flexed lower limbs, forward-tilted head, and compressed thorax, indicating that the grave pit was too narrow. The upper limbs were folded across the pelvis, and skeletal alignment shows the lower limbs were bent eastward.

Mechanical disturbance damaged much of the lower skeleton: only parts of the feet and pelvis remain intact, while distal elements such as metatarsals and phalanges are absent.

- **MGL011 (MAG20, SP4003):** *S. enterica* was recovered from individual SP4003. The individual was carefully placed within a mixed container built out of wood, stone slabs, and tiles, following a northeast-southwest axis. It was inserted into a quadrangular pit with vertical sides and a flat base. The construction technique and radiocarbon date indicate that the burial occurred in the 5th–6th century CE, during a phase of intensive occupation and active building development in the northern sector of the site.

##### 1.24. Nepluyevsky: NEP

*Country:* Russian Federation
*Region:* Kartalinsky district; Chelyabinsk region
*Coordinates:* 52.8826°, 60.1149°
*Sample Dates:* NEP018: SPb_2958; 3495 ± 55 BP;  1959–1636 cal BCE (2 σ)
NEP019: MAMS-45116; 3432 ± 27 BP; 1874–1630 cal BCE (2 σ)
NEP021: SPb_2961; 3521 ±  45 BP; 2008–1698 cal BCE (2 σa)
*Excavation/Sample Provenance*: Excavation Institute of History and Archaeology, Urals Branch of the Russian Academy of Sciences, Ekaterinburg, 2015–2017, license 2015-633, 2016-905, 2017-649 (S.V. Sharapova), together with Goethe University, Institute of Archaeological Sciences, Frankfurt am Main, Germany (E. Stolarczyk, A. Stobbe, R. Krause)
*Contact People:* Rüdiger Krause; Svetlana Sharapova; Astrid Stobbe; Eliza Stolarczyk; Marina Karapetian

*Site Description:*
The Srubnaya culture  (ca. 1900–1200 cal BCE) belongs to the so-called post-Sintašta period (Sintašta culture ca. 2010–1780 cal BCE) and thus to the Late Bronze Age (LBA)[26–29]. Its distribution extends across the forest steppe and steppe areas of eastern Ukraine along the Kama region and the banks of the Belaya and Ural rivers to Trans-Ural and western Kazakhstan. In the Trans-Ural area there is a strong interaction between the Srubnaya and Alakul' cultures (Andronovo family), which is regarded as the Srubnaya-Alakul' phase. The Nepluyevsky necropolis is also part of this phase, as Alakul' elements can be seen in the pottery decoration alongside the typical Srubnaya ornamental patterns.

The kurgan necropolis is quite large and consists of 38 kurgans[30]; during the 2015–2017 field campaigns,  three kurgans were excavated. Two small kurgans (Kurgans 5 and 9) yielded only non-adult burials, while under the mound of the biggest,Kurgan 1, individual primary and double burials of adults and children were recorded. Besides these, possible cenotaphs and structures for food offerings were placed within the funeral ground. The material analysed here was sampled from individuals buried in Kurgan 1. Kurgan 1 is characterised by a single-phase alignment. The relative dating of Kurgan 1 was initially determined on the basis of the ceramic typology of the vessels and assigned to the Srubnaya culture, a determination that was subsequently confirmed by numerous AMS radiocarbon dates, and thus belongs to the Late Bronze Age.

Funeral chambers were furnished with stone plates and removed sub-soil clay. This construction was then placed into the lower level of the kurgan mound. A study of the paleosol supports the hypothesis that the burial area initially functioned as a flat burial ground and was later covered with a kurgan mound. The most impressive interments, marked with rock-clay funeral expenditure, were mapped outside the geometrical centre of the kurgan and contained male and female skeletal remains. There is no specific order to the location of the juvenile burials. Double burials occurred among all age categories, i.e. juveniles, young adults, and older adults. The set of grave goods includes votive ceramic vessels (none of which appear to have been used), rings, bracelets made of bronze, and bronze temporal pendants plated with gold foil.

**Individuals infected with *Salmonella enterica* from Kurgan 1 of the Nepluyevsky necropolis**

- **NEP018**: Grave 25, 25–35 year old female. Calibrated date BCE (2σ range (95.4%)): 1873 to 1635. Intact. Location: the northernmost burial in kurgan 1. The grave goods included two ceramic vessels found to the east of the skull without any other items. The individual was lying on her left side in a crouched position (northwest-southeast), with the arms bent at the elbow, hands on the chin, and legs bent at the knees. The arm bones were placed one on top of the other, with the right resting on the left. There was an incompletely healed fracture of the diaphyseal part of the left ulna[31]. The grave pit was possibly filled with earth after the burial and covered with slabs.

- **NEP019**: Grave 26, 14–16 year old female. Calibrated date BCE (2σ range (95.4%)): 1875 to 1660. Intact. Location: Northeast sector, 2 m southeast of burial 25. The inventory consists of two ceramic vessels near the elbow bones and the knees, two relatively intact bronze bracelets (one on each arm) and fragments of at least two others. Also, several beads were located under the left shoulder blade and in the wrist area (a part was also found when the soil was sieved). Fragments of two bronze temporal pendants plated with gold foil were located on the right side of the lower jaw and on the left side under the jaw. This was the most richly furnished grave within Kurgan 1. The individual was lying on her left side in a crouched position (north-south). The right arm was raised towards the torso and bent at the elbow, the left arm bent at the elbow and lying slightly apart, the left hand near the individual's face,and the legs bent at the knees. The grave pit was covered with slabs.

- **NEP021**: Grave 30, 18–22 year old male. Looted. Calibrated date BCE (2σ range (95.4%)): 1879 to 1689. Located in the north-eastern sector, 2.6 m east of the zero reference point. The remains were found without anatomical classification in the various grave fill layers. The state of preservation is satisfactory, as nearly all skeletal elements are intact. However, some of the bones, including the skull, long bones, sacrum, ribs, and pelvis, were dislocated to one side, which may have occurred during the robbery. The grave goods consist of only a few fragments of pottery. Excessive strain on the spine was detected[31]. No slabs covered it.

Burials 25, 26, and 30 from Kurgan 1 are synchronous interments located in the eastern side of the ground. Females were placed on the periphery and males close to the central part. All are associated with the Srubnaya-Alakul phase, which was present in the region from ca. 1900–1600 BCE. The inferred family tree at Nepluyevsky spans three generations [32]. NEP019 was unrelated to the individuals in this extended pedigree; NEP018 was inferred to have married into this family, as her son and his father were both found at this site, while she was unrelated to anyone else in the burial. NEP021 was also part of this extended family, and was the cousin of NEP018’s son.

##### 1.25. Petting: PET

*Country:* Germany
*Region:* Bavaria, District of Traunstein near lake Waging
*Coordinates:* 47.9128°, 12.8162°
*Sample Date (PET001, PET002):* late 6th century CE
*Radiocarbon Date:* none
*Excavation/Sample Provenance:* Excavated between 1991 and 1993 by the Bavarian State Office for Monument Preservation (Bayerisches Landesamt für Denkmalpflege – BLfD). Stored at the SNSB, State Collection for Anthropology Munich (SAM).
*Contact People:* Michaela Harbeck; Brigitte Haas-Gebhard

*Site Description:*
The site of Petting, located approximately 1 km south of the southern shore of Lake Waging, was the focus of large-scale archaeological excavations conducted by the Bavarian State Office for Monument Preservation (Bayerisches Landesamt für Denkmalpflege – BLfD) between 1991 and 1993. Excavations took place in the area "Am Mühlfeld", designated for construction.

A total of 721 burials were uncovered, allowing for an almost complete investigation of the cemetery. Following the excavation, the finds were transferred to the BLFD, where they underwent conservation as part of the early medieval burial restoration program. The human remains were later transferred to the SNSB, State Collection for Anthropology Munich. The initial use of the cemetery has been tentatively dated to the second quarter of the middle third of the 6th century CE. [33,34]

- **PET001 (Grave/Ind. 342) and PET002 (Grave/Ind. 343)** – *S. enterica* was recovered from two individuals. According to osteological examinations, the individual from Grave 342 (PET001) is more likely to be classified as male (also according to the grave goods). The age can be estimated at 20–25 years. The individual from Grave 343 (PET002) is estimated to be between 7 and 9 years old, based on dental eruption. Due to the lack of skeletal maturity, an osteological determination of sex is not possible.

  Both individuals are part of a simultaneous triple burial together with burial 344 (which is likely a woman) dated to the late 6th century CE. Individual 343 was interred between and partially atop the others. The assemblage is notably sparse compared to contemporary graves. These factors could suggest a rapid burial, the body position of individual 342 further indicates a probable secondary grave disturbance.

##### 1.26. Plaošnik (Ohrid): PLO

*Country:* North Macedonia
*Region:* : Municipality of Ohrid
*Coordinates:* 41.1188, 20.7914
*Sample Date:* 800–500 BCE (contextual date). No direct radiocarbon dates have been produced for PLO010.

*Radiocarbon Date:* None
*Excavation/Sample Provenance:* Excavated between 2007–2014 under the auspices of the Institute for Protection of Monuments of Culture and Museum Ohrid and Dr. Pasko Kuzman.
*Contact People:* Daniela Heilmann; Aleksandra Papazovska; Tatjana Stojoska Vidovska

*Site Description:*
Plaošnik is a historically and culturally significant archaeological site with a long-standing historical importance, located on a plateau between Samuil's Fortress and the Church of St. John at Kaneo in Ohrid. In 893 AD, Saint Clement of Ohrid founded a monastery here, constructing the Church of St. Panteleimon over the remains of an earlier Early Christian basilica. Excavations have taken place since the 1940s at Plaošnik; between 2007 and 2014 the excavations uncovered an Iron Age cemetery.

- **PLO010** (PLA-321-2008): *S. enterica* was detected in a male individual with an estimated age at death of 15–17 years.

##### 1.27. Polwica: PLW

*Country:* Poland
*Region:* Polwica (site 5), Gmina Domaniów, Oława County, Lower Silesian Voivodeship
*Coordinates:* 50.9°, 17.1833°
*Sample Date:* Bronze Age, Únetice culture, 2021–1773 cal BCE (2 σ)
*Radiocarbon Date (PLW005, tooth, MAMS-52811):* 3576 ± 23 BP, 2021–1826 cal BCE (2 σ); *(PLW005, fibula, Poz-181986)*: 3560±35 BP, 2021-1773 cal BCE (2 σ)
*Excavation/Sample Provenance:* Excavated in 1998 and 1999 in connection with the construction of the A4 motorway, commissioned by the Institute of Archaeology and Ethnology of the Polish Academy of Sciences
*Contact People:* Mirosław Furmanek; Agata Hałuszko

*Site Description:*
The site is located approximately 30 km southeast of Wrocław in the Silesian Plain. It was excavated between 1998 and 1999 by the Institute of Archaeology and Ethnology of the Polish Academy of Sciences, Wrocław Branch, by a team of employees of the Archaeological Museum in Wrocław. Settlement and burial features of diverse time periods were discovered at this site, including the Neolithic period, the Únêtice culture, the Lusatian Urnfield culture, the La Tène culture, the Przeworsk cultures, and the Early Medieval period [35–37]. Among the discovered features, four middens were found, which contained mixed animal and human remains.

One of these features was pit number 3263, from which samples were obtained for genetic studies. The artefactual chronology of the pit is related to the Lusatian Urnfield culture (Late Bronze Age); however, the AMS 14C dating of the human remains (fragment of a child's fibula shaft) indicates a connection with the Únetice culture (Early Bronze Age): Poz-181986; 3560 ± 35 BP (2σ 2021–1773 cal BCE). Two clusters of Únêtice culture burials were recorded within the excavated area. One contained three and the other five graves. The small number of burials suggests that these cemeteries were severely damaged in later periods. Presumably, the skeletal human remains were placed in the small

(0.8 m x 1.34 m x 0.38 m) garbage pit secondarily, and their original context should be associated with the cemetery of the Únêtice culture.

- **PLW005 (Ft. 3263, Ind. 2)** – *S. enterica* was recovered from a subadult individual. Anthropological examinations could not identify the sex of the individual due to the high fragmentation of the bones and the young age. The age at death is estimated to be between 2–4 years, with a significant difference between the age estimated based on the rate of development of the dentition and the postcranial skeleton. Except for enamel hypoplasia no other pathologies were identified.

##### 1.28. Stříbrnice 1 - Lopaty: S1L

*Country:* Czechia
*Region:* Moravia
*Coordinates:* 49.3283°, 17.2458°
*Sample Date (S1L002):* Bronze Age, Corded Ware culture, 2574–2468 cal BCE (2 σ)
*Radiocarbon Date (S1L002, tooth, MAMS-48835):* 4003 ± 25 BP, 2574–2468 cal BCE (2 σ)
*Excavation/Sample Provenance:* Stříbrnice 1 – Lopaty. Excavated during rescue excavations by the Institute of Archaeological Heritage Brno (ÚAPP) between 2002 and 2003.
*Contact Person:* Jaroslav Peška

*Site Description:*
Between 2002 and 2003, the Institute of Archaeological Heritage Brno conducted rescue excavations along the D1 motorway route from Brno to Kroměříž and Lipník nad Bečvou. Archaeological investigations, led by Matějíčková and Peška, aimed to document prehistoric and historic remains prior to highway construction. At the site Stribrnice 1 – Lopaty, approximately 400 m from modern day Stribrnice, cemeteries of the Moravian Corded Ware and Bell Beaker culture were uncovered. A total of 14 Corded Ware and 84 Bell Beaker burials were documented, alongside four Bell Beaker settlement features. The remains were located on an elongated, elevated ridge between two watercourses, sloping southward into the Haná River valley[38].

- **S1L002 (Grave H92)** – *S. enterica* was recovered from a male individual. The grave was located at the periphery of the excavation area, approximately 40–50 m from the main burial cluster. The body lay in a right-flexed position, oriented west-east with the head facing east. Post-depositional disturbance of the skeleton was noted. Grave goods included Corded Ware vessels near the head and feet, an axe, a split adze behind the head, and animal bones placed in front of the face. A sample from the pars petrosa yielded a radiocarbon date of 2574–2468 cal BCE, placing the burial within the Corded Ware cultural horizon.

##### 1.29. Sapar-Kharaba, Tsalka: SAP

*Country:* Georgia
*Region:* Kvemo Kartli, Tkalka Municipality
*Coordinates:* 42.1939°, 42.1939°
*Sample Date (SAP004):* Late Bronze Age, $15^{th}$–$14^{th}$ centuries BCE
*Radiocarbon Date (SAP004, tooth, MAMS-59513):* 3061 ± 24 BP, 1408–1235 cal BCE (AMS, 2σ)
*Excavation/Sample Provenance:* Excavated between 2003 and 2005.

*Contact People:* Lia Bitadze; Nino Tavartkiladze

*Site Description:*
The Saphar-Kharaba burial ground is located in the Tsalka district, north of the village of Sphar-Kharaba in the Trialeti region. The cemetery covers an area approximately 1500 m x 500 m and dates to the 15$^{th}$–14$^{th}$ centuries BCE. Excavations conducted between 2003 and 2005 uncovered 115 pit graves, typically surrounded by cromlechs constructed from large basalt stones. Most graves were oriented north to east, with a single burial place at the cromlech's centre. The deceased were generally laid on their right or left side, facing north. Grave goods predominantly included ceramic vessels (thin, black-burnished pottery with polished decoration and coarse-grained, brownish wares adorned with relief bands). Additional artifacts included Near Easter-style daggers, scabbards, arrowheads, lancet-like weapons with bone handles, pyramidal stones, symbols of authority, and Mitannian cylinder seals of the "Common Style".

- **SAP004 (Grave 75)**: *S. enterica* was recovered from a male individual with a R1b1a1b Y haplogroup and evidence of steppe ancestry[39].

##### 1.30. Sirmium 55: SIA

*Country:* Serbia
*Region:* Sremska Mitrovica, Vojvodina
*Coordinates:* 44.9798°, 19.6095°
*Sample Date (SIA006):* ca. 300–500 CE
*Radiocarbon Date:* None.
*Excavation/Sample Provenance:* Cemetery excavated under the direction of Miloje Vasić in 1976/1977.
*Contact People:* Vujadin Ivanišević; Nataša Miladinović-Radmilović; Dragana Vulović; Ivan Bugarski

*Site Description:*
Sirmium is located in Sremska Mitrovica, strategically situated along the Sava River. The site was a Roman city in the Province of Pannonia and one of the most prominent cities of the Roman Empire in the Balkans. It functioned as an imperial residence and an important administrative and military headquarters in the 4$^{th}$ century CE. During the first half of the 4$^{th}$ century, emperors resided within their courts in the city, which boasted important facilities such as a mint, offices, as well as ceremonial spaces. The chronicles of Ammianus Marcellinus and Zosimus report the existence of a palation in Sirmium. The city also featured a circus, which has been located during excavations. Sirmium was also an important centre of Christianity in Late Antiquity.

Site 55 is located in the garden area at Palanka Street No. 63 in Sremska Mitrovica (Sirmium). Protective archaeological research was carried out in 1976 and 1977 on an area of 850 m², under the direction of M. Vasić. Then, in the so-called Eastern necropolis of Sirmium, the graves from the 1$^{st}$–4$^{th}$ centuries were discovered[40]. During the investigation of the place inside and around the Basilica of St. Irenaeus, more than 80 graves with three burial levels were discovered, the chronology of which could only be established relatively. The graves mostly belonged to the 4$^{th}$ century. A smaller number of graves were chronologically older than the basilica, so e.g. grave No. 68 belonged to a cremation from the 2nd century. Also, both grave No. 54 and the grave with frescoes were older than the basilica. It is certain that two levels of burial have been established at graves Nos. 38 and 44, where grave No. 44 is older. Three levels were found at graves Nos. 57, 81, and 82. The oldest is grave No. 57, then No. 81, and the youngest is grave No. 82. Two levels of graves that are certainly contemporary with the basilica were found in the northeast corner of the basilica, where grave No. 25 was "redone into two graves (Nos. 13 and 16)". Osteological material of human origin from the graves, which are chronologically older than the basilica, was not available for anthropological analysis[40].

Intensive burials continued in the spacious eastern necropolis in the 4th century, especially in the early Christian period. Numerous finds of graves and tombs from the end of the 19th and beginning of the 20th century are known thanks to I. Jung, who noted over forty graves, tombs, and tombstones from the Chikas stream to the east[41].

The deceased were buried in wooden coffins or in brick tombs, and exceptionally also in sarcophagi. 59 fragmentary inscriptions and tombstones and 11 fragments of mensa, circular and square, were discovered. In the immediate vicinity of the altar area, a marble slab was discovered with an inscription that identified the church with the martyr and the first historically proven bishop of Sirmium, St. Irenaeus. While the discovered inscription confirmed Bishop Irinej for the first time, in P. Milosevic's opinion it did not resolve the issue of the location of the martyrdom and the grave of the bishop, because it can still be assumed to be in the Roman necropolis and an important church centre on the right bank of the Sava[41].

- **SIA006 (Grave 86/1977, Ind. 1)** – *S. enterica* was recovered from an adult, female individual (ca. 50 years) with well-preserved but incomplete cranial and postcranial remains. Paleopathological analysis revealed osteoarthritis on the left calcaneus, while other bones showed no pathological changes. Dental analysis indicated a mixed degree of dental wear and moderate to severe periodontitis.

  The grave is located in the eastern necropolis of Sirmium and constructed of brick set on edge, with a tile-mortar base. It was oriented east-west, with the head to the west. The grave was situated within the apse of the Basilica of St. Irenaenus (335 CE). J. Guyon and M. Jeremić suggest that graves 86/1977 and 85/1977 may be part of a cemetery that predates that of the basilica[42].

##### 1.31. San Juan de Loarre: SJL

*Country:* Spain
*Region:* Huesca, Pre-Pyrenean outer mountain range
*Coordinates:* 42.314°, -0.6257°
*Sample Date:*
SJL010 and SJL019 are genetically identical from human aDNA analyses, so likely originate from the same individual.

*Radiocarbon Date (SJL010):* 4059 ± 21 BP, 2836–2489 cal BC (2 σ)
*Radiocarbon Date (SJL019):* 4137 ± 23 BP, 2872–2586 cal BC (2 σ)

*Radiocarbon Date (SJL020):* 4751 ± 22 BP, 3634–3383 cal BC (2 σ)
*Excavation/Sample Provenance:* Archaeological Rescue Excavations on 26 and 30 November, and 10 and 11 December 2007 conducted by Ma Victoria Pastor Sánchez and Diana Vicente Checa.
*Contact People:* Belén Gimeno Martínez; Marivi Pastor; Diana Vicente Checa

*Site Description:*
The sepulchral cave is located at the foot of Loarre Castle, in the Pre-Pyrenean outer mountain range of Huesca, at an elevation of 980 m. It was discovered in 2007 following a report of exposed human bones. Archaeologists Ma Victoria Pastor Sánchez and Diana Vicente Checa conducted an emergency excavation between November and December 2007, with official support from local and regional heritage authorities.

The cave measures approximately 2.5 m x 1.5 m and likely represents the collapsed remains of a larger cavity, used exclusively for collective burials during the Final Neolithic to Chalcolithic period, as confirmed by radiocarbon dating. The stratigraphy revealed a single anthropogenic layer, with highly fragmented human remains scattered throughout, showing no individualised burials but some evidence of anatomical groupings. A minimum of 47 individuals were identified, including both adults and children, with no age- or sex-based selection. Bodies were likely deposited in quick succession, without time for full decomposition. Grave goods included beads and pendants made of stone, shell, bone, and boar tusk, flint tools, fragments of handmade pottery, and charcoal traces (possibly linked to cremation-related practices). The cave was exclusively used as a burial site. Its isolated and hidden location supports interpretations of a non-visible burial site, disconnected from daily settlement life[43].

*S. enterica* was recovered from three tooth roots of two individuals:

- **SJL010 (individual 4) (later identified as identical to SJL019)**
- **SJL019 (individual 14) (later identified as identical to SJL010)**
- **SJL020 (individual 17)**

##### 1.32. Sármellék-Száraz eleje: SME

*Country:* Hungary
*Region:* Western Transdanubia
*Coordinates:* 46.7384°, 16.9152°
*Sample Date (SME005):* Late Copper Age Baden culture; 3520–3366 cal BCE (2σ)
*Radiocarbon Date (SME005, bone (rib), SUERC-106970):* 4653 ± 24 BP, 3516–3366 cal BCE (2 σ)
*Excavation/Sample Provenance:* Excavations between 2021–2022 by the Göcseji Museum in Zalaegerszeg
*Contact People:* Mária Bondár; Piroska Rácz; Dániel Gerber; Balázs G. Mende

*Site Description:*
The archaeological site of Sármellék-Száraz eleje is located in western Hungary, in Zala County, on a slope rising from the marshy valley of the Zala River. Excavations were carried out between 2021–2022 by the staff of the Göcseji Museum in Zalaegerszeg, prior to the construction of the M76 motorway and a connecting junction. A large area of more than 13 hectares, intensively inhabited in

many archaeological periods from the Early Neolithic to the Middle Ages was investigated, including settlement features (pits, food storage pits, cremation and inhumation burials, mass graves) of the Late Copper Age Baden Culture[44,45].

- **SME005 (Ft. 81, Ind. 81/1)** – *S. enterica* was recovered from individual 81/1. According to the anthropological examination the deceased was a woman aged 20–29 years. The bony remains of individual 81/1 consist of a fragmentary and incomplete skull with mandible and an incomplete and fragmentary skeleton. There is an injury on the left middle ulnar shaft, a well-healed fracture which was accompanied by injury to the interosseous antebrachial membrane, indicated by a small exostosis and the uneven surface of the left radius. Each of the bones of the left forearm shows a slight periosteal reaction. A healed fracture of the right first metacarpal was also observed, with significant shortening of the bone and osteomyelitis. The remaining lower limb bones (both femora and the right tibia and fibula) also show periosteal reaction without any observable traumatic event. The parietal bones show porotic hyperostosis. A dental disorder, peg-shaped upper lateral incisors could be observed in SME005[46].

This research was initially part of an NRDIO-supported project (NKFI K-128413) "Complex Analyses of the Late Copper Age Burials of the Carpathian Basin". We would like to thank István Eke for granting us the right to process the finds, and for providing us with the excavation documentation.

##### 1.33. Sion, Petit-Chasseur: SPC

*Country:* Switzerland
*Region:* Sion, Valais
*Coordinates:* 46.2319°, 7.3508°
*Sample Date:* Late Neolithic or Bell Beaker cultural horizon; 4213 ± 25 BP (2899–2697 BCE; IntCal20, OxCal v4.4; ETH131355) to 3711 ± 24 BP (2199–2029 BCE; IntCal20, OxCal v4.4; ETH131360)
*Radiocarbon Date:* None
*Excavation/Sample Provenance:* Discovered in 1961; excavated 1961–1969 (O. J. Bocksberger); 1971–1973 (A. Gallay). Further work: 1987–1988 (S. Favre and M. Mottet); 1992 (M. Besse); 2003 (M. Mottet).
*Contact People:* Jocelyne Desideri; Déborah Rosselet-Christ

*Site Description:*
The site of Petit-Chasseur, located in the heart of the town of Sion in Valais, is one of the most significant Neolithic sites in Switzerland and a key reference for Alpine prehistory. Discovered in 1961 during urban development works, the site has yielded a remarkable cultural sequence spanning from the Middle Neolithic to the Second Iron Age[47].

The earliest levels reveal Middle Neolithic habitation, including domestic structures and animal enclosures dated between 4500 and 3600 BCE. These occupations are characterised by cereal cultivation, goat herding, household-based craft activities, and Chamblandes-type funerary cysts, often associated with juvenile burials[48].

The Petit-Chasseur site stands as one of the most important megalithic necropolises of the Late Neolithic in Switzerland and Western Europe. This exceptional funerary complex, composed of twelve monuments including dolmens and cist graves, documents a continuous and evolving sequence of mortuary practices spanning from the Late Neolithic to the Early Bronze Age (3200–1600 BCE), with a significant Bell Beaker phase in between. Excavations carried out from the 1960s through the late 1980s have established a detailed stratigraphic and chronological framework for the site's development. Several major phases of occupation have been identified in the necropolis.

The initial occupation of the megalithic area corresponds to the beginning of the Late Neolithic (ca. 3100 BCE) and is marked by the construction of dolmen MXII, a structure with triangular foundations and lateral antennae. Its funerary chamber contained the remains of 115 individuals, making it the most informative burial for this period in the region[49]. Occupation of the site continued with the construction of a second dolmen with triangular foundations and lateral antennae - dolmen MVI - which presents a more complex occupation history spanning approximately eight centuries, from the Late Neolithic to the Bell Beaker period (ca. 3000–2200 BCE). Built after dolmen MXII, this monument underwent distinct phases of use. The first, attributed to the Late Neolithic, corresponds to the construction and initial occupation of the structure, although the original placement of the burials is difficult to reconstruct due to later disturbances. These were caused by a second phase during which Bell Beaker groups cleared the funerary chamber to accommodate their own burials[50]. The Bell Beaker groups adopted a hybrid approach. Alongside the reuse of dolmen MVI, they constructed their own monuments - three side-entrance dolmens (MI, MV, and MXI) and six small funerary chambers (MII, MIII, MVII, MVIII, MIX, and MX).

The subsequent phase of activity, attributed to the Early Bronze Age, is characterised by the construction of a circular pit adjacent to the outer wall of dolmen MVI. It was used for secondary cremations of human remains originating from one or more earlier burials. Bell Beaker material recovered from within the pit suggests a potential link with this cultural horizon, potentially explaining the absence of human remains in several cists and dolmens conventionally attributed to this group. The pit contained the cremated remains of at least 91 individuals and may also have included individuals dating from the Late Neolithic. Archaeological evidence suggests that the contents of the pit were deposited in a single, intentional event[51].

The site’s final funerary use occurred in the Second Iron Age (La Tène), as evidenced by seven graves aligned according to a standardised orientation and furnished with rich grave goods, including weapons and ornaments[47].

More than 200 individuals were buried in these collective graves. A wide range of archaeological material has been recovered: abundant ceramics, flint and rock crystal tools (including blades, arrowheads, circle segments, and cores), a polished stone industry with numerous arrowheads, ornaments made of silver, gold, and various seashells, archer’s wrist guards, V-perforated buttons, spindles, polishers, and a copper awl. Most striking, however, are the 31 richly decorated anthropomorphic stelae, which have brought international recognition to the site[52].

Petit-Chasseur stands out for its exceptional stratigraphic depth, quality of archaeological finds, and extensive documentation. It remains a pivotal site for understanding Neolithic and Bronze Age societies in the Alpine region and beyond. Since its discovery, it has generated a wealth of academic research and continues to be a focus of interdisciplinary study and institutional collaboration.

- **SPC020** (FI_05_MVI JD 022 FN/BB): This sample was recovered from the Early Bronze Age circular pit at Petit-Chasseur, which contained disarticulated human remains. In this case, the sample corresponds to a permanent lower left first molar, whose precise chronological attribution remains uncertain, though it may be associated either with the Late Neolithic or the Bell Beaker cultural horizon. It can therefore be placed during the 3rd millennium BCE, within a chronological range spanning from 4213 ± 25 BP (2899–2697 BCE; IntCal20, OxCal v4.4; ETH131355) to 3711 ± 24 BP (2199–2029 BCE; IntCal20, OxCal v4.4; ETH131360). Ongoing genetic analysis of the tooth also indicates that this individual was male.

This research was initially carried out as part of a project supported by the FNS (project 10521F-205059/2022-2025, PI J. Desideri) "Towards a renewed vision of alpine agropastoral societies through the analysis of diets, lifestyles and settlement dynamics", an interdisciplinary research project that uses a multi-proxy approach to examine lifestyles, health status and the evolution of diets and subsistence habits, as well as population dynamics, mobility and kinship.

##### 1.34. Tavan Tolgoi: TAV

*Country:* Mongolia
*Region:* Ongon sum, Sukhbaatar aimag
*Coordinates:* 45.357°, 113.1421°
*Sample Date:* Early Mongol period: site dates starting 1048–1255 CE; ending 1175–1347 CE
*Radiocarbon Date:* None
*Excavation/Sample Provenance:* Excavations conducted in 2004 by Department of Anthropology and Archaeology, National University of Mongolia
*Contact Person:* Erdene Myagmar

*Site Description:*
The Tavan Tolgoi site ('Five Holy Hills') is located in southeastern Mongolia and has been excavated by the Department of Anthropology and Archaeology (University of Mongolia) since 2004. The burial grounds lie on the southern slopes of a central hill, ca. 47 km southeast of the district centre, within a historically significant region situated between Karakorum and Dadu, the former capitals of the Mongol Empire and Yuan Dynasty. During excavations in 2004 and 2005 ca. 20 graves were found, indicating high-status burials. The grave goods consist of several elaborate offerings, such as gold ornaments, ceremonial objects, and horse burials with gilded saddles. Radiocarbon dating of human bones, textiles, and coffin wood yielded calibrated dates ranging from ca. 1030–1280 CE, aligning with the period of the Great Mongol Empire and fitting with the lifetime of Genghis Khan and the rise of the Golden Horde lineage[53].

- **TAV007 (Grave 2, AT-619):** *S. enterica* was recovered from a young adult male individual (ca. 30 years old). The individual was found in Grave 2 in a supine position, with the right arm placed beneath the torso, grasping a silk-wrapped golden *Jins* – an emblem traditionally associated with elite social rank. Osteometric analysis of the skeletal remains yielded a stature estimate of 169.8 cm.

The carbon dates of this site are published in Youn et al. (2007)[54].
Stable isotope analysis of this site is published in Fenner et al. (2014)[53].

##### 1.35. Thebes: TBS

*Country:* Greece
*Region:* Thebes, Boeotia (Central Greece)
*Coordinates:* 38.3310°, 23.3630°
*Sample Date (TBS012):* Late Roman/Early Byzantine Period, 247–376 cal CE (2 σ)
*Radiocarbon Date (TBS012, tooth, MAMS-55010):* 1743 ± 16 BP, 247–376 cal CE (2 σ).
*Excavation/Sample Provenance:* Excavations between 2011 and 2017 under the direction of Stephanie Larson and Kevin Daly and the local Greek Ephorate of Antiquities.
*Contact People:* Stephanie Larson, Kevin Daly, Maria A. Liston, Alexandra Charami, David R. Scahill

*Site Description:*
The site is located on the Ismenion Hill, southeast of the ancient city of Thebes in Central Greece, and has been an important topographical and cultural place for over three millennia. Due to its elevated position, it was a significant place for funerary, religious, and later domestic and industrial activity, with occupations attested from the Late Bronze Age through the Byzantine period. Archaeological evidence indicates that the hill was in use as early as the 15th century BCE, functioning as one of the three principal Mycenaean chamber tomb cemeteries in Thebes. By the 8th century BCE, it became the site of a prominent sanctuary to Apollo Ismenios, featuring three successive temples: an early Geometric structure, an Archaic Doric temple, and a later Doric kiosk, likely unfinished, whose foundations are still visible today. In Late Antiquity, the hill was transformed into a Christian and later Byzantine cemetery, in use from the 3rd to 12th centuries CE. This burial function may be connected to the nearby Church of the Evangelist Luke. From the 11th century onward, light domestic or industrial use is also attested.

The site was excavated in the early and middle years of the 20th century. More recent excavations were carried out between 2011 and 2017 by a team from Bucknell University (Pennsylvania, U.S.) under the direction of Stephanie Larson and Kevin Daly and in collaboration with the local Greek Ephorate of Antiquities.

- **TBS012 (Ind. 11, Grave 19):** *S. enterica* was recovered from a male adult (ca. 35–50 years). Despite the poor state of skeletal preservation, the cranial and pelvic joint morphology indicate the sex and age estimation. The skull showed nasal bone loss consistent with leprosy (rhino-maxillary syndrome). Additional hand and foot bones in the grave also display leprous changes. Graves 19 and 20 differ from others at the site by showing no evidence of secondary manipulation or exhumation. This pattern may reflect a sudden epidemic.

##### 1.36. Targap: TGP

*Country:* Kazakhstan
*Region:* Almaty
*Coordinates:* 43.3303°, 75.8432°
*Sample Date (TGP001):* Saka culture, 376–203 cal BCE (2 σ)
*Radiocarbon Date (TGP001, tooth, MAMS-49764):* 2226 ± 19 BP, 376–203 cal BCE (2 σ)

*Excavation/Sample Provenance:* Excavations by R. K. Sherbaev in 2018.
*Contact People:* L. Djansugurova; L. Musralina; E. Khussainova

*Site Description:*
Targap is a village located in the Zhambyl District of Almaty Oblast, Kazakhstan, and is administratively part of the Samsinskiy rural district. It lies approximately 40 km west of the village of Uzynagash. The area is notable for the presence of burial mounds dating to the Early Iron Age Saka culture.

- **TGP001:** *S. enterica* was recovered from an individual from mound 1.

##### 1.37. Tesárske Mlyňany: TML

*Country:* Slovakia
*Region:* Nitra
*Coordinates:* 48.3319°, 18.3544°
*Sample Date (TML028):* ca. 400–600 CE
*Radiocarbon Date:* none
*Excavation/Sample Provenance:* Rescue excavations in 2002 and 2003.
*Contact People:* Matej Ruttkay; Mária Krošláková

*Site Description:*
The site is located in southwestern Slovakia, ca. 3 km southwest of the Zlaté Moravce. The cemetery is situated on a gentle, southeasterly slope in the Gočol field. It lies about 400 m west of the original meandering bed of the Žitava River. Rescue excavations have examined 98 graves. It is the largest burial site from the 5$^{th}$–6$^{th}$ centuries in Slovakia north of the Danube.

The excavated burial pits exhibit a slender oval to nearly rectangular plan and are mostly oriented west to east with the head to the west. Structural elements such as traces of wooden chests, grave hollows, and stone slab coverings were identified in many cases. A significant number of graves displayed secondary disturbances, but no clear evidence of robber shafts was observed. These patterns are consistent with 5$^{th}$ to 6$^{th}$ century grave "plundering" practices common in the Middle Danube region. Artificial cranial deformation was confirmed in eight individuals, reinforcing cultural links with Germanic or Hunnic traditions.

Due to the secondary disturbance of the graves, grave goods were relatively sparse, but included beads, iron knives, earrings, belt buckles, fibulae, belt fittings, bone and shell pendants, iron arm rings, tweezers, and combs. Pottery was almost entirely absent, which is unusual for graves of this period.

The site is attributed to a Suebian population substrate with East Germanic or Hunnic influences. It represents an important addition to the archaeological record of the post-Attilan transition and the early migration period in the Middle Danube region. Alongside evidence from recently discovered settlements, it supports the continued Germanic presence in parts of southwestern Slovakia, particularly in the Žitava and Hron regions, until the arrival of early Slavic groups at the end of the 5$^{th}$ and beginning of the 6$^{th}$ centuries.

- **TML028 (Grave 44):** *S. enterica* was recovered from a female individual who died between 17–19 years[55]. The individual was buried in a rectangular grave pit with rounded corners and the body was oriented west-east with the head to the west. Only the lower legs remained in anatomical position; the femurs were found displaced above them. Bones from the upper body, including arms and a skull fragment with teeth, were heaped in the western part of the grave. Associated finds included green and black glass beads, a fragment of a bronze plate ornament, and small bronze pieces. The skeletal remains were poorly preserved: the cranium was missing, with only seven loose teeth recovered, and the postcranial skeleton was severely eroded. Osteological features indicate a juvenile individual (17–19 years old), likely female based on pelvic morphology, with weak entheseal development and a generally gracile postcranial skeleton.
  Preliminary investigations date the grave to the 5$^{th}$ to first half of the 6$^{th}$ centuries.

### 2. Methods

##### 2.1 Sampling and Screening

Teeth were sampled for each individual presented here for ancient DNA analysis (see **Table S7** for details of which teeth were sampled and specific library protocols used). DNA was extracted as described by Dabney et al. (2013)[56], and either single or double stranded libraries were prepared as described in Gansauge et al. (2020)[57] (for single stranded) or Meyer and Kircher (2010)[58] (for double stranded libraries). Some libraries were treated fully with Uracil DNA Glycosylase (UDG) (to mitigate the impact of postmortem damage), while others were treated with an inhibitor after 30 minutes to retain some signature of post-mortem damage to allow authentication (UDG-half). Some libraries were not UDG treated. Details of specific library protocols, UDG treatment, and sampling strategy are presented in **Table S7**.

Low-depth Illumina shotgun sequencing on an Illumina HiSeq4000, NextSeq500, HiSeq2500, or NovaSeqX (see **Table S7** for sequencing-level metadata) was performed to screen these individuals for ancient host and pathogen DNA. The HOPS pipeline was used to assess the evidence for ancient pathogen DNA [59]. Those with strong evidence for an *S. enterica* infection (i.e. reads assigned specifically to *S. enterica* subsp. *enterica*, with strictly declining edit distance and evidence for post-mortem damage) were brought forward for in-solution capture following Vågene et al. (2018)[60] and further analysis. The captured libraries were more deeply sequenced using either paired or single-ended sequencing (see **Table S7**).

##### 2.2 Data Processing

**Sequencing Data**

Raw, demultiplexed FASTQ files were processed and aligned to the *S. enterica* serovar Paratyphi C RKS4594 reference genome (accession: NC_012125.1) using nf-core/eager v2.5.1 [61]. Sequencing reads were trimmed to remove sequencing adapters using AdapterRemoval v2.3.2 [62], and paired-end reads overlapping by ≥11 bp were merged. Collapsed and single-end reads were aligned to the Paratyphi C reference genome using bwa aln v0.7.17-r1188 with loose mapping parameters (-l 16, -n 0.01) to account for errors due to post-mortem deamination [63]. Aligned data was filtered to remove duplicates using Picard's MarkDuplicates v2.26.0 [64], and filtered for a minimum read length of 34 and mapping quality of 37. In order to mitigate the impact of post-mortem deamination on variant calling, 5 bp from the 5' end of single-stranded non-UDG sequences and 2 bp from the 3' end were trimmed using BamUtils v1.0.15; for double stranded untreated libraries, 5 bp were trimmed from both ends; for partial UDG treated sequences, 2 bp from each end were trimmed[65]. Where relevant, sequencing data from different libraries were merged into sample-level BAM files, and reads were "soft-clipped" - i.e. the first and last two base calls for each read had their quality scores reduced to 2. Read length distributions and post-mortem damage were quantified using DamageProfiler v0.4.9[66].

Single nucleotide variants were called using GATK's UnifiedGenotyper v2.0-35-g2d70733[67], with ploidy set to 2 to allow for the investigation of mis-mapping and potential contamination, and using the option EMIT_ALL_SITES. SNPs were then called for analysis with a dataset of published ancient and modern genomes (**Table S4**) , using the tool MultiVCFAnalyzer v0.87-alpha[68], excluding conserved and repetitive regions as in Vågene et al. (2018)[60]. A minimum base quality of 30, read depth of 3, and support of 90% was required for basecalling.

In order to estimate the impact of cross-mapping and contamination on each sample, that is, overall levels of heterozygosity in non-masked regions of the genome, VCFs were filtered for sites passing UnifiedGenotyper quality filters, with a minimum read depth of 3. Overall contamination levels were estimated by calculating the mean minor allele support across the genome. This was also evaluated for non-transition sites, to calculate the lower bound of contamination estimates. The total number of non-reference and non-reference heteroplasmic sites was also calculated, to assess the impact of contamination and/or mismapping across the genome. Additionally, average heteroplasmy in 10kB windows (sliding across the genome in 1kB steps) was calculated to assess the distribution of heteroplasmic sites across the genome.

**Background Metadata**

In order to assess our ability to draw any conclusions from the changing distribution of *Salmonella* serovars in time and space, we visualised an anonymised representation of the sample set used for screening. This included any non-cranial skeletal elements sampled from Eurasia within the last ten thousand years, which had been screened in an automated pathogen screening pipeline before the 31st of January 2024 (the cutoff date for inclusion in this study). The screening data were split into 2500-year time bins (8000–4000 BCE; 4000–2500BCE; 2500–1000BCE; 1000BCE–500CE; 500CE-present), and latitude and longitude coordinates were jittered within a 2 degree range of their reported locations. Kernel density estimation was performed over the geographic range of Eurasia, using the R packages raster v. 3.6.30 and spatstat v. 3.3.2, and plotted using a land mask downloaded from the R package pastclim v. 2.2.0 [69–71].

##### 2.3 Maximum Likelihood Phylogenetic Analysis

For initial placement in phylogenetic trees, a SNP alignment of representative modern *S. enterica* subsp. *enterica* genomes and quality-filtered ancient *S. enterica* genomes with a minimum coverage of 3X was generated using MultiVCFAnalyzer v0.87-alpha[68], using a minimum required read depth of 3, and a minimum of 90% reads supporting a call, and with masking as in Vågene et al. (2018)[60]. This SNP alignment was filtered for maximum site-level missingness of 5%, and a maximum-likelihood phylogenetic tree was constructed using IQTREE v. 1.6.12, using *S. enterica* subsp. *arizonae* as an outgroup. ModelFinder was used to find the best-fitting substitution model, and 1000 ultra-rapid bootstraps were used to assess support [72–74]. This procedure was repeated for each of the low-quality samples individually to estimate their placement in the tree.

In order to investigate relationships within the Para C lineage more closely, a SNP alignment was created using representative modern genomes and ancient genomes over 3X from the Para C lineage, as above, but using the Birkenhead lineage as an outgroup.

**2.4 Bayesian Phylogenetic Analysis**

Bayesian phylogenetic analysis, implemented in BEAST2 (v2.7.5)[75], was used to investigate the timings of different evolutionary events within the Para C lineage, as well as to investigate changing bacterial population sizes through time. A more stringent coverage filter was used for this analysis, to optimise data quality while still maintaining sufficient diversity to represent our Para C dataset. A SNP alignment was created from variant calls from representative modern Para C samples and ancient samples with radiocarbon dates or secure contextual dates (in the case of historical samples), with at least 50% of their genome covered at 4X, using MultiVCFAnalyzer as above but with a minimum of 4 reads and at least 90% of the total reads supporting a call. This SNP alignment was filtered for a maximum of 5% missingness, and a maximum likelihood tree was constructed using IQTree, as above.

Radiocarbon dates for ancient samples were recalibrated using OxCal v4.4 with default parameters and the IntCal20 calibration curve [76,77]. A root-to-tip regression was performed on the Para C lineage maximum likelihood tree using the tool TreeTime [78], with median radiocarbon dates or reported sampling dates as tip dates.

The full genome alignment from MultiVCFAnalyzer and maximum likelihood tree were used to infer potential recombination using the tool gubbins v3.3.5[79]. The alignment was masked to remove potentially recombinant regions, SNP sites were extracted using the tool snp-sites, and the number of constant A, T, C, and G nucleotides were counted using snp-sites v2.5.1[80]. This SNP alignment was filtered to remove sites with more than 5% missingness. Strict and relaxed lognormal clock models were compared using nested sampling, as implemented in the NS package for BEAST2 [81], and a relaxed lognormal clock model was chosen.

Independent BEAST runs were set up using BEAST Model Test to select a substitution model[82], with a relaxed lognormal clock model and under a Coalescent Bayesian Skyline tree prior. Uniform tip date priors were used for all ancient samples, with the minimum and maximum at the extremes of the 2 sigma range for their radiocarbon date. Tip date priors for modern samples were set to a range of one year around their reported sampling dates. A monophyletic constraint was placed on all Birkenhead and non-Birkenhead (i.e. Para C lineage) samples. Each run was set up to have a total chain length of 1500 million, with a pre-burnin of 1.5 million. The sensitivity of our inference to the specified priors was assessed by running analyses that only sampled from the prior, and comparing the 95% HPD of parameters of interest between runs using sequence data and those sampling from priors. This comparison is shown in **Figure S16**.

Once BEAST runs had converged, log and tree files were combined using LogCombiner v2.7.6, with a 10% burnin. The underlying tree was estimated using conditional clade distributions as implemented in BEAST2's TreeAnnotator [83]. This tree was then visualised and explored using the R packages ggtree v3.13.0 and treeio v1.26.0 in R v4.3.3 [84,85].

##### 2.5 Pseudogenisation and Adaptation

###### Data Processing

Adapter trimmed, collapsed sequencing reads from ancient samples were realigned to an *S. enterica* pangenome reference, filtered to include only genes which were covered 100% by the capture probes used in data generation, as in Key et al. (2020)[86]. BWA mem with the clipping penalty disabled ( `-L 0` ) was used for alignment, in order to correctly align reads overlapping with the ends of genes in the pangenome reference.

BAM files were filtered to remove PCR duplicates using Picard v1.140, and ancient files were "softclipped" using a custom filtering script [87] (i.e. the base quality scores at the ends of reads were reduced to 2, so that bases more likely to be impacted by post-mortem deamination would not be used in genotype calling). For fully UDG treated libraries, 2 bp at each end were softclipped, for half-UDG treated data this was increased to 5 bp, and for non UDG treated data this was increased to 7 bp. Bam files were then merged to sample level, indel realignment was performed using GATK v2.0-35-g2d70733 [67], and bams were filtered for a minimum read length of 34 and minimum mapping quality of 37 using samtools v1.3.

Variants were then called using GATK's UnifiedGenotyper v2.0-35-g2d70733[67], with a minimum base quality of 30 required, and with the options `-glm BOTH -gt_mode DISCOVERY -out_mode EMIT_ALL_SITES` in order to call both SNPs and indels. A ploidy of 2 was used to allow filtering based on read depth and percentage support at variant sites as above.

The raw VCFs were then filtered based on read depth at each variant site using a custom script. Variants were retained if they had a coverage within two standard deviations of the mean coverage calculated from the Paratyphi C alignment, with an absolute minimum support of 3 reads, and with a minimum of 90% of reads at a site supporting a call.

The presence of genes within each sample was inferred based on the percentage of the gene covered at 1X, as calculated by bedtools coverage v2.25.0 [88]. A gene was classified as present if ≥90% was covered at ≥1X.

Modern data was processed and filtered in an identical manner, with the exception of the softclipping step.

###### SNP Annotation

VCFs for all samples were merged, invariant sites were removed, and multiallelic variant sites were atomised using bcftools norm (v1.20) [89]. Variants were annotated using SnpEff v5.2c, with a custom database created from the pangenome reference file, using the bacterial and plant plastid codon table [90].

Pseudogenising mutations were defined as those with a "HIGH" impact from the SnpEff annotations; i.e. those which are predicted to cause frameshift or nonsense mutations. Initial exploration of pseudogenisation rates showed that samples with mean Paratyphi C coverage < 5X and with fewer than 3750 genes covered ≥90% had unreliable pseudogene rate estimates, due to the impact of poor coverage on both SNP and indel calling. Therefore, these samples were excluded from downstream analysis. The modern dataset was restricted to a representative subset of genomes for serovars with known host specificity (see **Table S4**).

**Pseudogene rates and association with host specificity**

Per-genome pseudogenisation rates were calculated by counting the number of genes covered ≥90% at 1X with at least one pseudogenising mutation; this was normalised by the total number of genes "present" (*i.e.* the total number of genes covered ≥90% at 1X). The structure of shared pseudogenes in the dataset was explored by plotting a heatmap of shared pseudogenes between genomes, ordered by hierarchical clustering implemented in the R package pheatmap v1.0.12; this structure was also explored by performing principal component analysis on the shared pseudogene count matrix via the stats package (v4.3.3) in R.

Pearson's correlation coefficient between pseudogenisation rate and sample date was calculated for ancient genomes over 5X using the stats package (v4.3.3) in R [91].

Association between host specificity and pseudogenisation was first explored in the modern dataset by performing a chi-squared test for each gene, as implemented in R. This association analysis was restricted to modern genomes, as the host specificity of ancient genomes is unclear. The list of genes with an uncorrected p-value of 0.05 was then assessed to see if there was any enrichment for gene ontology terms relative to the pangenome background, using goatools v1.2.3[92]. GO annotation of the pangenome was performed using InterProScan v5.72-103.0[93]. Multiple testing was accounted for in GO enrichment analysis by applying a Benjamini-Hochberg (FDR) correction, as implemented in goatools.

**SNPPAR: convergent pseudogenisation**

Convergent pseudogenisation events in host-adapted genomes were investigated using the tool SNPPARv1.0 [94]. A maximum-likelihood tree was constructed using a SNP alignment from data aligned to the Paratyphi C reference genome, with a 95% partial deletion filter (as in **2.3 Maximum Likelihood Phylogenetic Analysis**). Annotated SNPs for analysis were restricted to the "core genome" - *i.e*. genes covered ≥90% in ≥95% of samples in the dataset - to mitigate the impact of missingness on analysis. This tree file, along with annotated SNPs were used as input to SNPPAR, which was used to place mutations at different nodes in the tree. Private mutations were excluded from analysis, to mitigate the impact of errors in SNP calling, whether from post-mortem deamination, contamination, or alignment errors. Each variant was then annotated based on the percentage of revertant descendant nodes in the tree, and variants with ≥5% offspring nodes carrying a revertant (i.e. ancestral) allele were filtered out. Each node in the tree was annotated as host generalist, host specific, or ancestral, based on the host specificity of offspring nodes. Nodes ancestral to both ancient and host-adapted genomes were annotated as being host-adapted, as it was assumed that if a mutation was fixed in both ancient and modern host-adapted genomes, it likely was relevant to host adaptation.

Recombination is a source of genomic variation in *S. enterica*; however, we wished to exclude the possibility that observed convergent pseudogenes were due to more recent recombination between already host-adapted strains. In order to do this, we excluded sites with identical mutations at different nodes in the tree. A parallel analysis retaining these mutations resulted in largely similar results.

For each gene, the number of independent pseudogenising, missense, and synonymous mutations were summarised. In order to identify genes with excessive pseudogenisation relative to a putatively neutral background mutation rate, the expected number of pseudogenes in a gene was calculated as the number of synonymous mutations in that gene, normalised by the total number of identified synonymous mutations, scaled by the total number of identified pseudogenising mutations. An analogous calculation was done for missense variation. Genes where this difference (termed "delta pseudo") was in the 97.5 percentile or above, were kept as genes with "excessive pseudogenisation". This percentile was chosen as the distribution of "delta missense" was approximately normal, so we kept anything outside of the middle 95% of genes; as genome reduction is the primary driving force behind the evolution of bacterial pathogenicity [95], we then kept those with an excess of pseudogenes relative to synonymous mutations, rather than those with an excess of synonymous mutations, which we expect to reflect conserved genes. This subset of genes were then filtered to include just those with at least one convergent pseudogenisation event between two host-adapted lineages, and to exclude any with pseudogenes in host generalist lineages.

In addition, the full list of genes with at least one convergent pseudogenisation event between two host-adapted lineages was analysed, to investigate whether there were any functional categories enriched relative to a background list of all genes included in the analysis, using the tool goatools, as before [92]. When this list was restricted to mutations occurring ancestral to all sequences in these lineages, no terms passed multiple testing correction.

The R packages treeio v1.26.0, data.table v1.16.2, tidyverse v2.0.0 and ggtree v3.13.0 were used in the analysis and visualization of convergent pseudogenes [84,85,91,96,97].

**2.6 Salmonella Pathogenicity Islands**

In order to assess the presence and absence of salmonella pathogenicity islands (SPIs) 1–10, all ancient and modern genomes over 3X mean coverage were aligned using nf-core/eager[61] (as above) to the *Salmonella enterica* Typhi CT18 reference genome (accession: NC_003198.1), which contains these pathogenicity islands. The linear reference genome rather than the pangenome was used for alignment in order to use a reference which conserved the order of these genes in relation to each other. Coverage across genes and pathogenicity islands of interest was calculated using bedtools coverage[88]. The proportion of these loci covered at least once was visualised using the R package pheatmap[98].

In addition, ancient samples were aligned to the Paratyphi C pSPCV virulence plasmid (accession: NC_012124.1) using nf-core/eager[61], as above (**2.2 Data Processing**).

Sequence variation within the differentially present pilin locus in SPI-7 was also assessed. Variants were called and filtered as in the pseudogenisation section, and the resulting VCF was filtered to include just the pilin locus and individual samples with a maximum of 25% per-individual missingness. This VCF was converted to a multiFASTA file using a custom python script, as in Jackson et al. (2024) [99]. This multiFASTA was aligned using ClustalW, as implemented in SeaView v5.0.5 [100,101]. The relationship between pilin sequences was investigated by calculating the raw distance between pairs of sequences (i.e. the number of nucleotide differences between pairs), using the dist.dna function from the ape package (v5.8.1) in R, and using a pairwise deletion approach to disregard sites where one or both samples had missing data [102]. A maximum likelihood tree was constructed from the full pilin sequence alignment and visualised using ggtree [84,103]. Non-synonymous variation at this pilin locus was also assessed using SnpEff with the pre-computed Typhi CT18 database [90].

##### 2.7 Modern Genome Processing

Modern *S. enterica* assemblies were processed similarly to the ancient data. This was done by splitting assemblies into "pseudo-reads" of length 100, with a 1 bp slide across assemblies. These reads were then aligned to either the Paratyphi C, Typhi CT18, or pangenome reference (as appropriate for the analysis used) using either bwa aln (or bwa mem for the pangenome reference), with identical parameters as for the ancient data, and using the same filtering steps. No softclipping was performed, as no post-mortem damage would be observed.

### 3. Quality Control: Authenticity of ancient fragments; authenticity of genome reconstruction and impact of mismapping.

A number of different metrics were assessed in order to confirm that the recovered *S. enterica* genomes were authentic and of reasonable quality for analysis. First, in order to confirm that these genomes came from ancient DNA fragments in the libraries and not modern contaminants, post-mortem deamination patterns and fragment length were assessed. Then, in order to confirm that the mapping reads came from an *S. enterica subsp. enterica* source, evenness of coverage and edit distributions for each alignment were investigated. Finally, the impact of mismapping and post-mortem damage on variant calling was assessed by investigating the proportion and distribution of heteroplasmic sites across the reference genome for each sample.

##### 3.1 Ancient DNA Authentication

In order to authenticate the ancient origin of these pathogen genomes, post-mortem damage patterns and mean and median aligned read lengths were assessed using DamageProfiler, implemented through nf-core/eager [61,66] (see **Methods 2.2** for details). These parameters were estimated separately for each library sequenced.

5' C to T misincorporation rates are presented in **Figure S1,** and 3' G to A misincorporation rates are presented in **Figure S2.** Most samples display expected damage patterns for their library preparation and UDG treatment type. Libraries from the sample KNL025 displayed some slightly elevated misincorporations at the 3 prime end of their reads: this was flagged as a potentially problematic sample.

Read length distributions are displayed in **Figure S3.** The majority of samples have very short mean read lengths, consistent with ancient DNA libraries. A small number of samples had longer average read lengths (CPA002, HDL004, KPI001, NEP021, and MGL011) - these all represent very well-preserved samples in terms of coverages retrieved, and as these all had reasonable post-mortem damage patterns, they weren't considered a concern. KNL025 and KNL019 show some excess of unpaired reads in their non-UDG treated single stranded libraries relative to double-stranded libraries.

##### 3.2 Alignment Authentication

In order to confirm that the aligned ancient reads belonged to ancient *S. enterica* and not some other, related taxon, the evenness of coverage and distribution of distances between the reads and reference genome was assessed. This is summarised in **Table S1**, where the percentage of the genome covered between 1–5X is summarised, and a summary statistic for the edit distance distribution (as described in Hübler et al. (2019)[59]) is also presented.

##### 3.3 Heteroplasmy and mismapping impact on SNP calling.

In order to assess heteroplasmy (and therefore the impact of contamination on our analyses), each site in the genome with a minimum depth of coverage of 3 which was not masked during SNP calling (as described in Vågene et al. (2018)[60]) was assessed. The number and proportion of sites affected, the average minor allele support genome-wide (with and without transitions, which are more likely to be impacted by post-mortem damage), and the average minor allele support at heteroplasmic positions are presented in **Table S8**. For samples with mean coverage <5X, an average minor allele support of 33% generally represents one discordant read. Additionally, the distribution of these heteroplasmic sites were assessed by calculating the average heterozygosity in 100kB rolling windows, with a step size of 10kB (**Figure S4-7)**. The proportion of heteroplasmic sites was generally low, with the highest proportion estimated in the lowest coverage samples. This is due to the minimum read depth filter on our SNP calls, which (in general) is much higher than the mean expected coverage for these samples, so shows some bias towards sites more likely to be impacted by mismapping. One exception to this is the sample KNL025, which had been flagged as potentially problematic due to generally high misincorporation rates when assessing the impact of post-mortem damage. Additionally, MAJ022 and ABF001 (both with coverages ≥3X) had a relatively high proportion of sites impacted by heteroplasmy (≥1% sites). For both of these samples, heteroplasmic sites are distributed evenly across the genome, rather than concentrated around certain peaks, which would be closer to the expected distribution for mismapped environmental reads (as seen in other samples: see e.g. TAV007 for a high quality example). When the analysis is restricted to transversions only, MAJ022 more closely resembles a "clean" genome, while ABF001 still has reasonably high levels of heteroplasmy across the genome. This suggests a potential mixed-strain infection for ABF001, or contamination from a close enough relative of *S. enterica* Paratyphi C that mismapping across the entire genome (rather than conserved regions) was possible. For MAJ022, post-mortem deamination may still have had an impact, despite the extensive trimming and data cleaning prior to analysis.

All samples were impacted to some extent by mismapping, as measured by heteroplasmy. In general, this impacted high coverage (≥5X) samples much less than lower coverage samples. Any samples with greater than 0.3% estimated contamination rate were excluded from any analyses requiring high-quality SNP calls.

### 4. Candidate genes from convergent pseudogenisation analysis

Of the 26 candidates for pseudogenes adaptive to host specificity, 11 are pseudogenised in the Para C lineage; of these, eight are pseudogenised in a subset of the ancient genomes. These genes include a lipoprotein (possibly a bacterial alpha-2-macroglobulin), *yciW, yegE, dkgA, yqjG, scsB, phoN,* a putative DNA/RNA non-specific endonuclease and a putative cell wall-associated hydrolase

*phoN: Acid Phosphatase*
We replicate the association between phoN pseudogenisation and host adaptation found in Key et al. (2020)[86]. The product of *phoN* has been shown to be recognised as an antigen by mice infected with *S.* Typhimurium [104], and it has also been shown to be a target antigen in human *S.* Typhi infections [105]. *phoN* is regulated by the phoP/Q two component system, which is an important regulator of virulence in a range of Gram negative bacteria [106].
Here, we were able to identify where in the Para C lineage this pseudogene was acquired (at the MRCA of serovars Typhisuis and Paratyphi C), and dated this acquisition to 890-510 BCE (95% HPD) using BEAST2.

*dgkA: 2,5-diketo-D-gluconic acid reductase A*
The gene *dgkA* is also part of the phoP/Q regulon in *S.* Typhimurium [106], and was pseudogenised in the Paratyphi C lineage (480 -230 BCE) as well as in Abortusequi. PhoP and PhoQ are a two-component system, present across many bacterial genera, which respond to low Mg2+ levels, acidic pH, increased osmolarity and short-chain fatty acids, relevant to the changing environments found during the infection process [107]. Differences between the PhoP/Q regulon in *S.* Typhi and *S.* Typhimurium have been observed previously [108]. This includes the observation that *phoN* is not regulated in the same way in Typhi as in Typhimurium, hinting that changes in this regulatory network are important in the evolution of host specificity.

*scsB: Suppressor for copper-sensitivity B*
Another candidate pseudogene observed in our dataset was *scsB*, pseudogenised in modern and ancient Choleraesuis genomes (470 BCE-40CE) , as well as in modern Paratyphi A genomes. This is part of the scs operon, involved in resistance to copper toxicity and redox stress in *S.* Typhimurium [109]. Ladomersky et al. (2017) show that macrophages induce copper toxicity in *S.* Typhimurium infections in mice [110], highlighting the importance of copper stress in host-pathogen interactions.

*yegE: diguanylate cyclase/phosphodiesterase*
The gene *yegE* is pseudogenised in both Abortusequi and Abortusovis serovars, as well as being pseudogenised in the subset of ancient genomes from Kamenice (KNC), Albania (MRCA 1470-950 BCE), which form a clade basal to modern Para C serovars. Cyclic diguanylate is a messenger molecule, often involved in the regulation of bacterial motility and adherence [111]. Although this specific gene appears to be poorly studied in *S. enterica*, a study of diguanylate cyclases and phosphodiesterases in *E. coli* have shown that *yegE* deletion mutants show defects in adherence to

bladder epithelial cells [112]. Although it is not clear exactly what impact this pseudogenisation event would have on the phenotype of these strains, there is some evidence that curli biosynthesis, and hence rdar ("red, dry and rough") colony formation would be impacted, which implies increased anthroponotic transmission, as in modern-day invasive non-typhoidal salmonella [113,114].

*yqjG: glutathione-dependent reductase*

*yqjG* is a glutathione-dependent reductase, pseudogenised in modern and ancient genomes in the Choleraesuis lineage (10-260 CE). This gene has previously been reported to be highly up-regulated in *S. enterica* on rotting plant material, and was also found to be upregulated in *S.* Enteritidis in a chicken intestinal infection [115], likely due to increased oxidative stress. This suggests that this gene is relevant for intestinal colonisation, which may be less relevant for invasive serovars [116].

*STM1940: putative cell wall-associated hydrolase*

The cell wall-associated hydrolase *STM1940* was found to be pseudogenised in serovar Typhisuis (MRCA 910-2020 BP) and was found to be pseudogenised in all ancient representatives of the Paratyphi C lineage (MRCA 550-290 BCE). Although this gene does not appear to be well characterised, it has also been identified as being associated with invasive lineages of *S. enterica* in a study of invasive non-typhoidal salmonella Typhimurium in Malawi [117]. This suggests that, despite limited experimental characterisation of this gene, its pseudogenisation is relevant for the evolution of invasive salmonella infection, both today and in the past. Some work characterising the expression of this gene shows that it tends to be upregulated in macrophages, and that its expression is regulated by the PhoP/Q two component system [118].

*STMMW_25491; STMMW_13371; yciW*

As well as these three candidates with clear roles in host-pathogen interactions, it is not fully clear how *STMMW_25491*, *STMMW_13371* and *yciW* contribute to host adaptation. *STMMW_25491* encodes a putative lipoprotein, which was annotated as a bacterial alpha-2-macroglobulin by KEGG orthology, hypothesised to be involved in defence against host antimicrobial strategies [119]. *STMMW_13371* is some kind of endonuclease, with an unknown role, while yciW is a cytoplasmic protein of unknown function, which is upregulated in iron-deficient conditions [120], and has been shown to been disrupted in *S.* Typhimurium U288 strain S01960-05, which was isolated in pigs from the UK [121]. Interestingly, this strain is associated with reduced human infections; the pseudogenes observed in our dataset occur at nodes ancestral to animal-adapted strains (Typhisuis and Abortusovis), although it also appears to be pseudogenised in Paratyphi C.

*ddlA: D-alanine--D-alanine ligase A*

The gene *ddlA* was pseudogenised in both Typhisuis and Paratyphi A genomes. This gene is involved in peptidoglycan biosynthesis, and the resulting structure is a target of the antibiotic vancomycin [122]. Although the presence of D-Ala-D-Ala in peptidoglycan is extremely conserved [123], this pseudogene was previously reported in Paratyphi A [124].

*fdoG: Formate dehydrogenase O, alpha subunit*

*fdoG* was pseudogenised in serovars Typhisuis and Abortusequi; this pseudogene has also been reported in some Paratyphi A genomes [124]. This gene is important for gut colonisation in serovar Typhimurium; however, this is less relevant for host-specific, invasive serovars [125].

*yhiI: Auxiliary transporter membrane fusion protein*

The gene *yhiI* is pseudogenised in modern Choleraesuis genomes (with the exception of 57_1_CS), as well as a serovar isolated from harbour porpoises (labelled Fulica-like), and would have been acquired between 450-1210 CE. An insertion at this locus has been shown to be relevant to enteric fever, but it is unclear whether this disrupts the *yhiI* gene [126].

As well as these Para C lineage pseudogenes, we also observe excessive pseudogenisation in genes in other host-adapted lineages.

*dacD: D-alanyl-D-alanine carboxypeptidase*

The gene *dacD* is pseudogenised in serovars Abortusovis, Abortusequi and Typhi. This is a penicillin binding protein with poorly defined functions, but has been implicated in penicillin resistance and its deletion resulted in increased biofilm formation in *S. enterica* and *E. coli* [127]. Like DdlA, DacD is also involved in peptidoglycan remodelling; it was suggested that its role in beta lactam resistance may be (indirectly) related to this [127]. However, its role in peptidoglycan remodelling appears to be relatively minor; in an *E. coli* dacABC triple mutant, only 5% D-alanyl-D-alanine carboxypeptidase activity is observed (relative to wild type), suggesting that pseudogenisation of this gene may not have a large impact on phenotype [128].

*STMMW_00881_1: sulfatase*

The gene labelled "*STMMW_00881_1*" in the pangenome shows high sequence identity with a sulfatase-like ester hydrolase. There is not much literature on this, and the gene is annotated as a pseudogene in the pangenome, suggesting that the loss of this gene may not have much of a phenotypic impact.

*slrP: leucine-rich repeat protein SlrP*

Convergent pseudogenisation was identified in the leucine-rich repeat protein *slrP*, in serovars Typhi, Paratyphi A, and Abortusequi. SlrP is an effector protein with ubiquitin ligase activity, translocated into eukaryotic host cells by *Salmonella* type 3 secretion systems [129]. It was first identified in a screen for genes impacting host-specificity in *S.* Typhimurium: slrP mutants show defects in colonisation of murine Peyer's patches [130]. It interferes with a human chaperone (ERdj3), and interferes with antigen presentation by dendritic cells [131,132]. This pseudogene was also identified in all ancient genomes (including the Birkenhead lineage), but this was then filtered out as potential recombination.

It is upregulated by PhoP under SPI-2 inducing conditions, emphasising the role of the phoP/Q system in virulence and host adaptation in *S. enterica* [133].

*dkgA 2,5-diketo-D-gluconic acid reductase A - paralog in Para C*

A homolog of *dkgA* (2,5-diketo-D-gluconic acid reductase A) was also pseudogenised in serovars Abortusequi and Typhi. Although these genes were annotated separately in the reconstructed pangenome, it is likely that they perform a similar function, and are likely both regulated by the PhoPQ system.

*dmsA: dimethyl sulfoxide reductase subunit A*

Both serovars Paratyphi A and Typhi have independent pseudogenes in the gene *dmsA*; there are also a subset of genomes from serovars Gallinarum and Abortusequi which carry a *dmsA* pseudogene. This is involved in anaerobic metabolism and is important in *S.* Typhimurium gut colonisation [134]. Nuccio and Bäumler (2014) show that the degradation of anaerobic metabolic networks is a common motif in the evolution of host restriction in *S. enterica*[135] .

*gspE: Type II secretion system protein*

GspE is a type II secretion system protein in *S. enterica* [136]. It has ATPase activity, and allows the formation of a bacterial pilus via the export of pilin proteins [137]. It is pseudogenised in serovars Fulica and Abortusovis, as well as in a subset of serovar Gallinarum genomes.

*STMMW_00181: chitinase*

*STMMW_00181* encodes a chitinase, and is pseudogenised in serovars Typhi and Abortusovis, and in a subset of serovar Gallinarum. Chitinases have been shown to aid *S.* Typhimurium to invade the mouse small intestine; however, the authors also demonstrate that chitinase-deficient mutants are capable of surviving in systemic infections [138].

*malT: HTH-type transcriptional activator*

malT acts as a transcription factor of genes involved in maltose metabolism [136]. This gene has been shown to be extremely upregulated under anaerobic shock conditions in transcriptomic experiments in *S.* Typhimurium [139]. In addition, in an experiment assessing the proteome of epithelial-cell invading *S.* Typhimurium, it was shown that maltose utilisation is extremely downregulated during epithelial cell invasion [140]. It was pseudogenised in serovars Abortusovis and Typhi.

*mrdA: Penicillin-binding protein 2*

The gene *mrdA* seems to be pseudogenised in serovars Abortusovis and Typhi. However, in previous transposon mutagenesis experiments [141,142], it was shown to be essential in both serovars Typhi and Typhimurium. The mutations identified here were a premature stop in both *S.* Abortusovis (p.Trp81*) and *S.* Typhi (p.Gln390*). Perhaps despite these predicted pseudogenisation events, there is still some low levels of translation, as in Feng et al. (2022)[143]; we stress that our identification of gene disruption events in both modern and ancient genomes is purely based on genomic analysis, and further experimental work could help resolve both what the functional impact of these disruption events were, and whether or not pseudogenes were actually formed.

*STMMW_27361; STMMW_39741; STMMW_37801; STMMW_27701*

There are also a number of pseudogenes for which there is little literature, and where the impact on the evolution of host specificity and invasiveness is unclear. These are summarised briefly below. The IS3 family transposase STMMW_27361 is pseudogenised in serovars Paratyphi A, Typhi and Abortusequi. STMMW_39741 is a putative hydrolase pseudogenised in serovars Abortusequi and Abortusovis, as well as in a subset of Paratyphi A genomes. STMMW_37801 is annotated as an aldose 1-epimerase family protein, and is pseudogenised in serovars Abortusovis and Fulica.

STMMW_27701 is annotated as a transcriptional regulator, and is pseudogenised in serovars Typhi and Abortusovis.

#### 5. Supplemental Figures

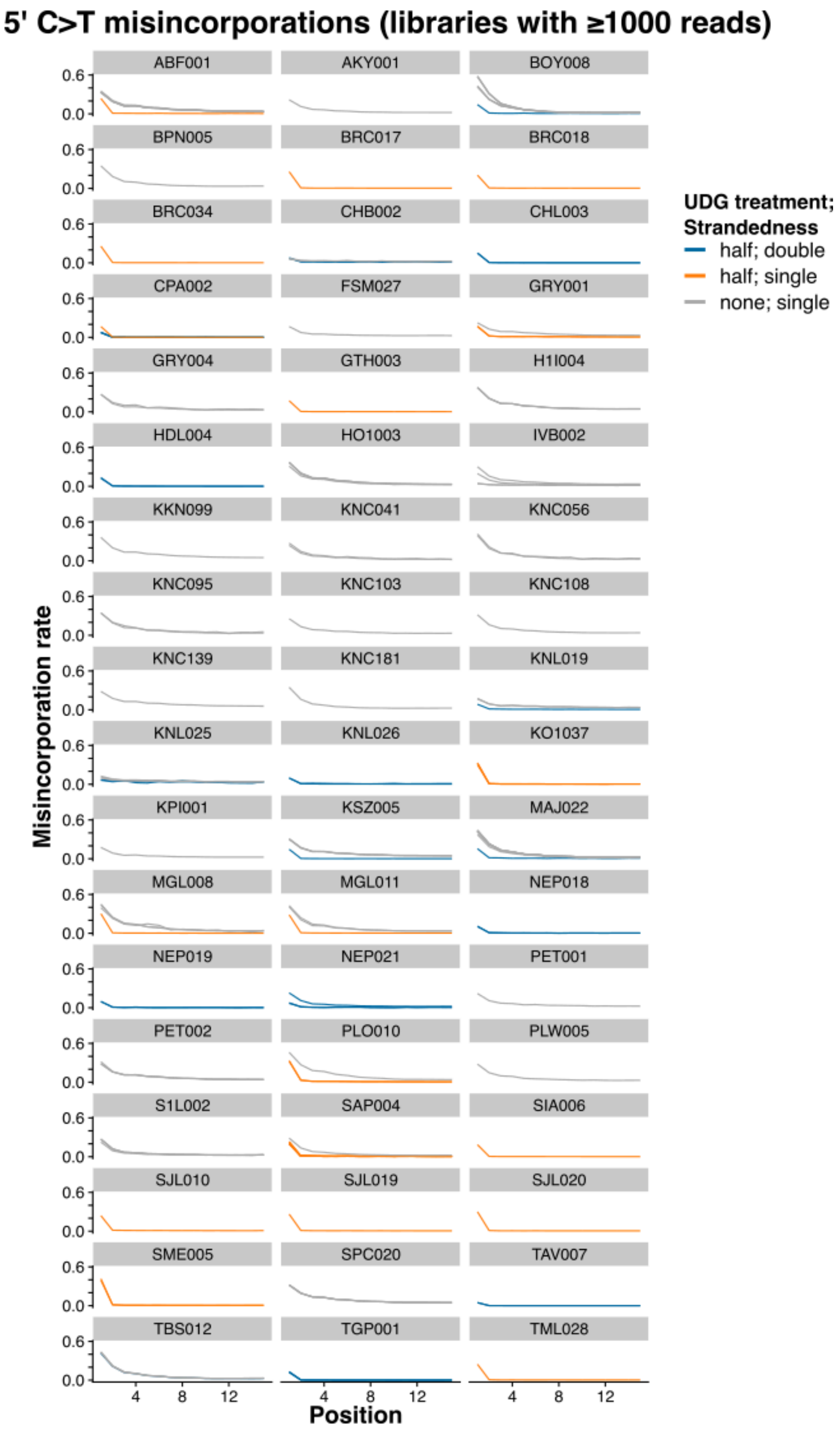

**Figure S1. 5’ C to T misincorporation rates.**

Misincorporation rates are plotted separately by library type, for libraries with ≥ 1000 reads mapping to the *S. enterica* genome: double-stranded half-UDG is blue, single-stranded half-UDG is orange and single-stranded non-UDG is grey.

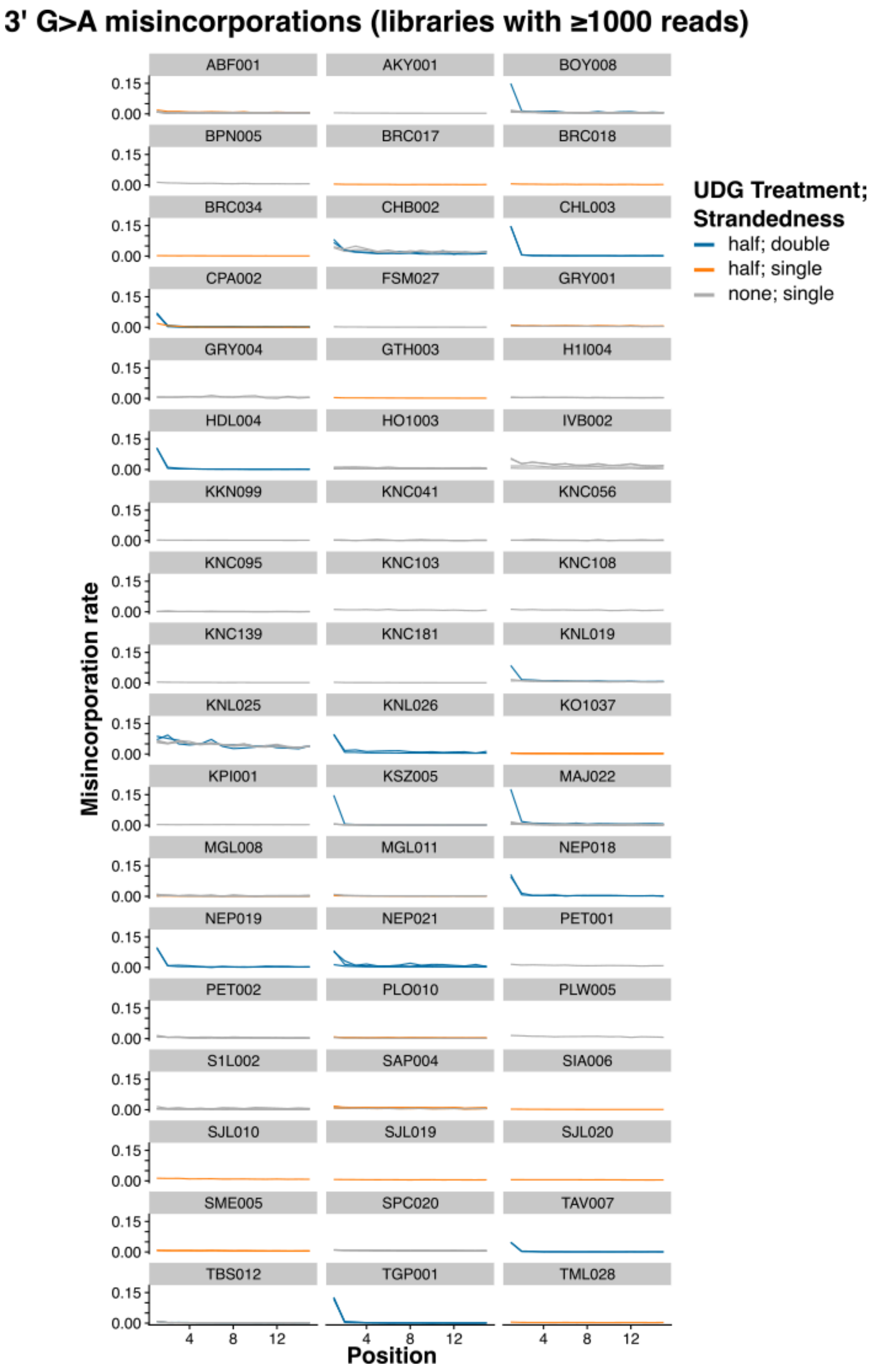

**Figure S2. 3' G to A misincorporation rates.**

Misincorporation rates are plotted separately by library type, for libraries with ≥ 1000 reads mapping to the *S. enterica* genome: double-stranded half-UDG is blue, single-stranded half-UDG is orange and single-stranded non-UDG is grey.

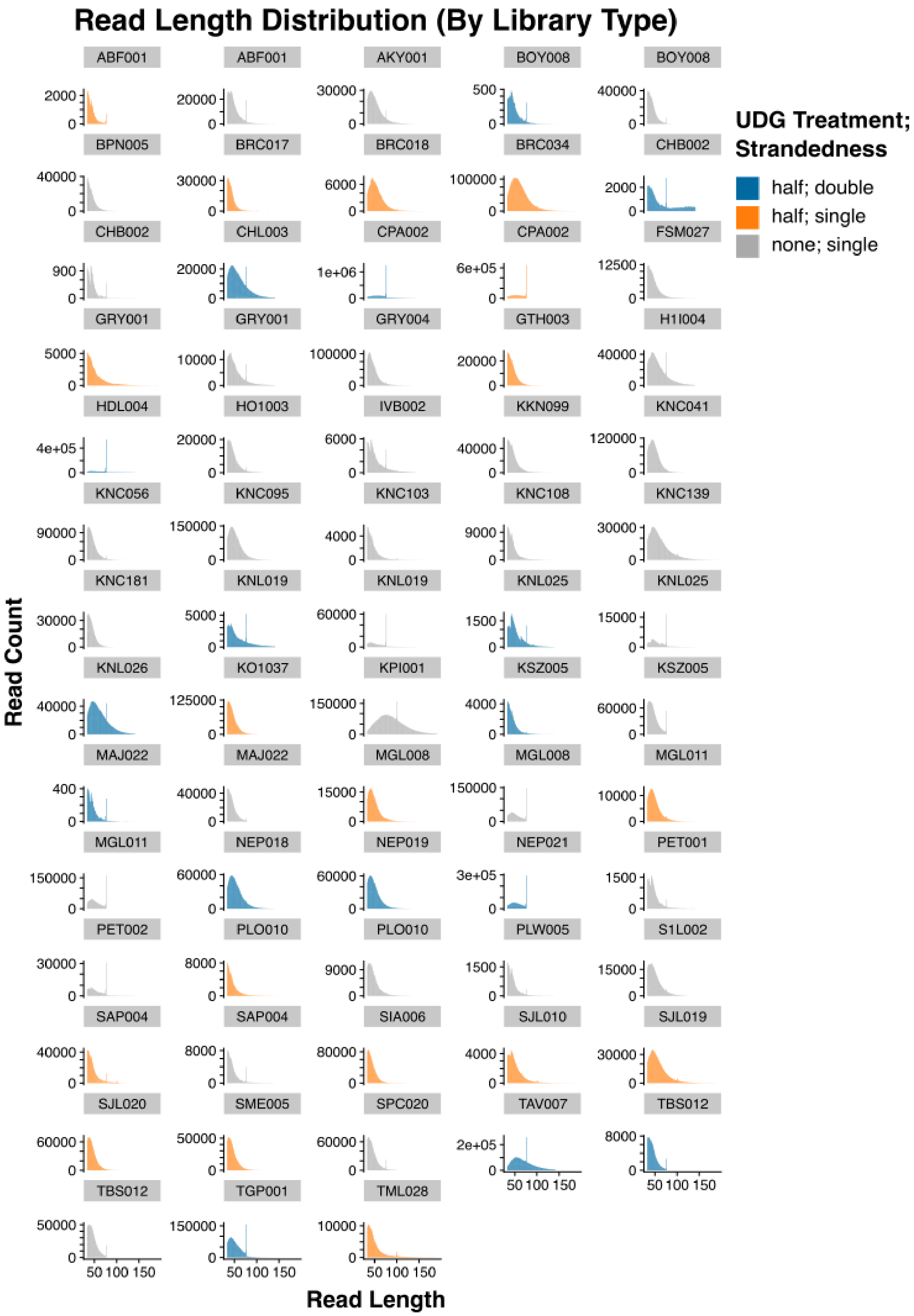

**Figure S3. Read length distribution for each sample and library type.**
Barplots are coloured by library type: double-stranded half-UDG is blue, single-stranded half-UDG is orange and single-stranded non-UDG is grey.

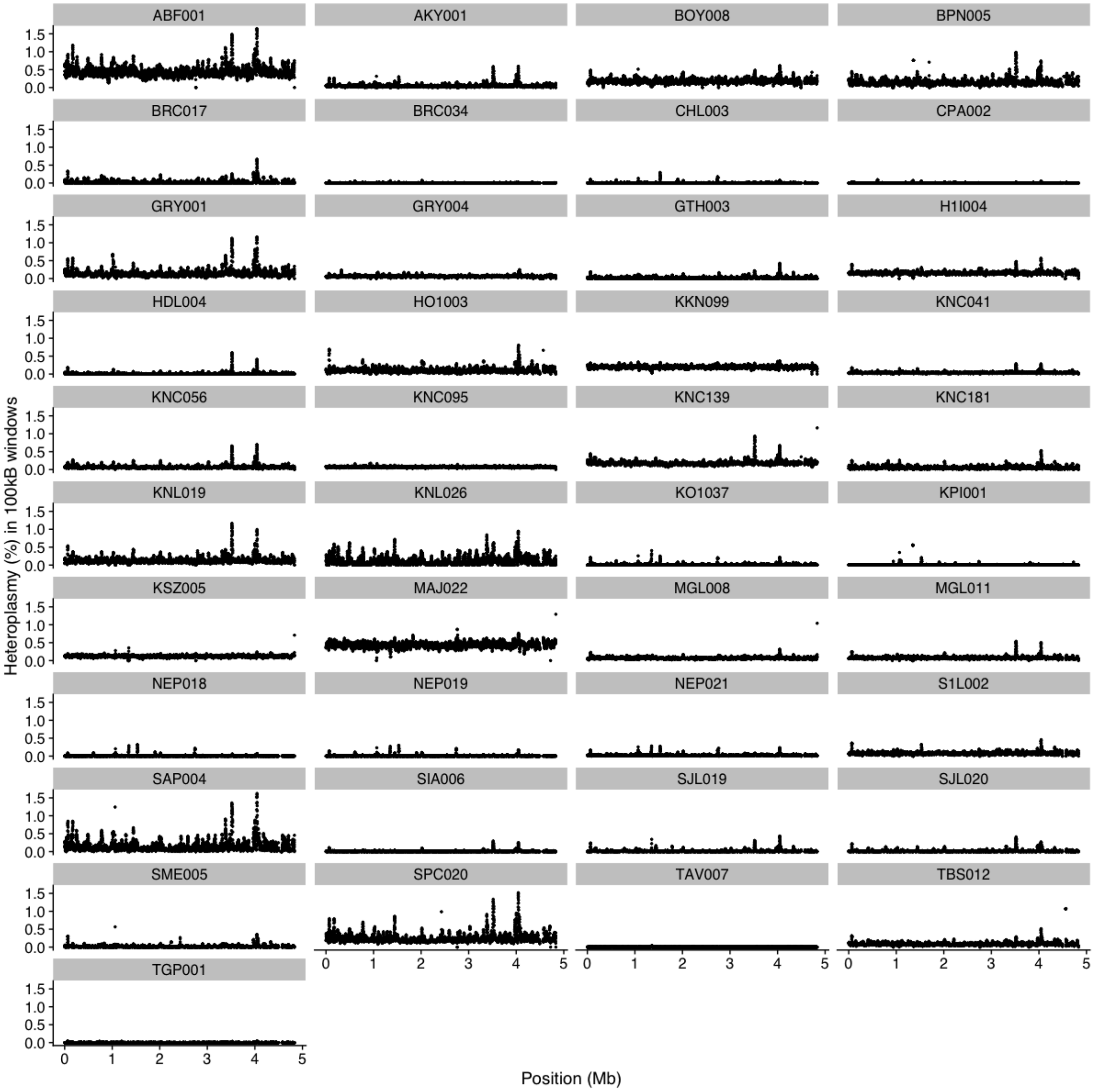

**Figure S4. Estimated heterozygosity in 100kB windows with 10kB step size for samples ≥3X.** Includes possible post-mortem deamination. The percentage heterozygosity in 100kB windows along the genome is plotted for each individual.

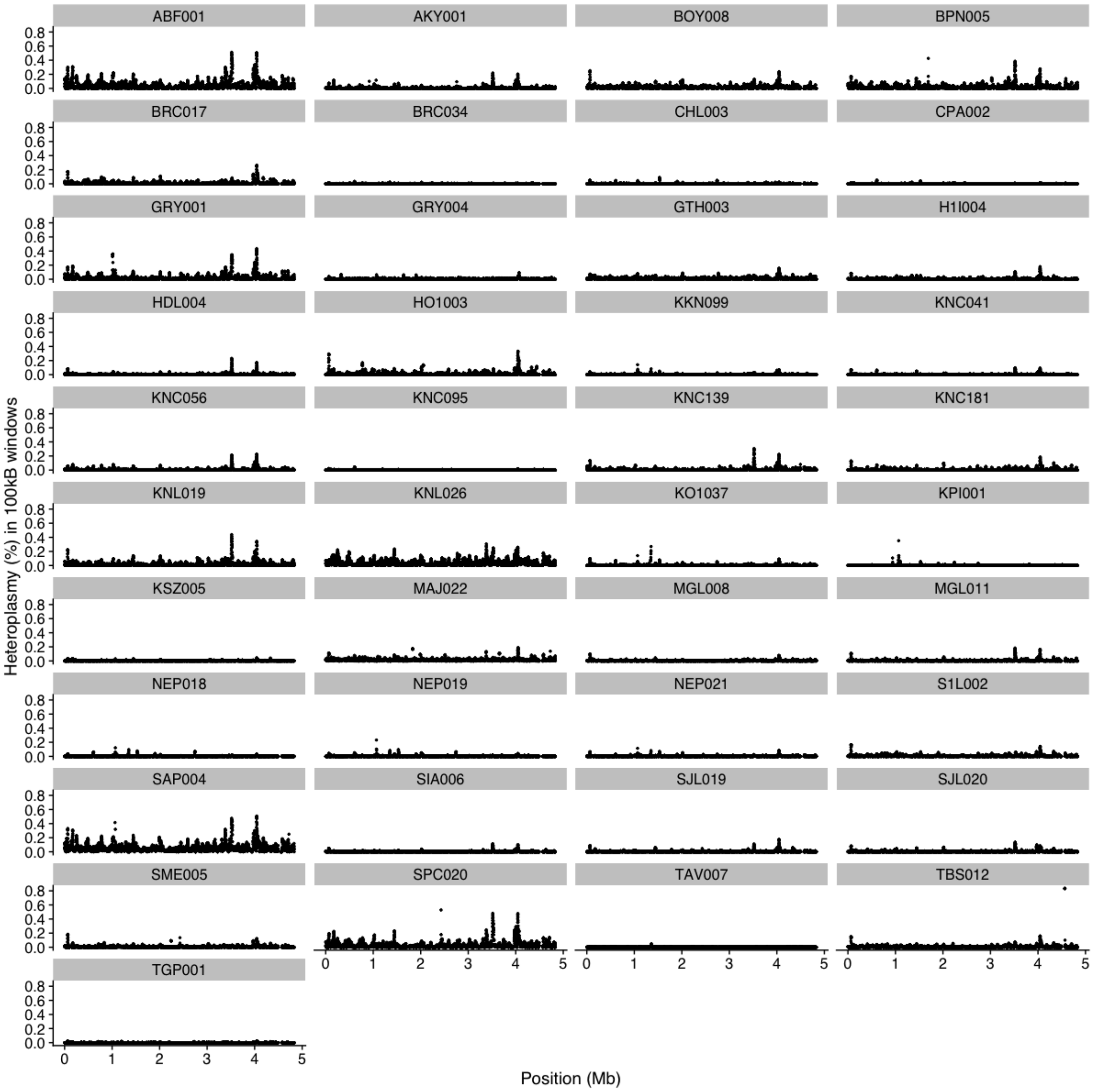

**Figure S5. Estimated heterozygosity in 100kB windows with 10kB step size for samples ≥3X (without possible post-mortem deamination).**

The percentage heterozygosity in 100kB windows along the genome is plotted for each individual.

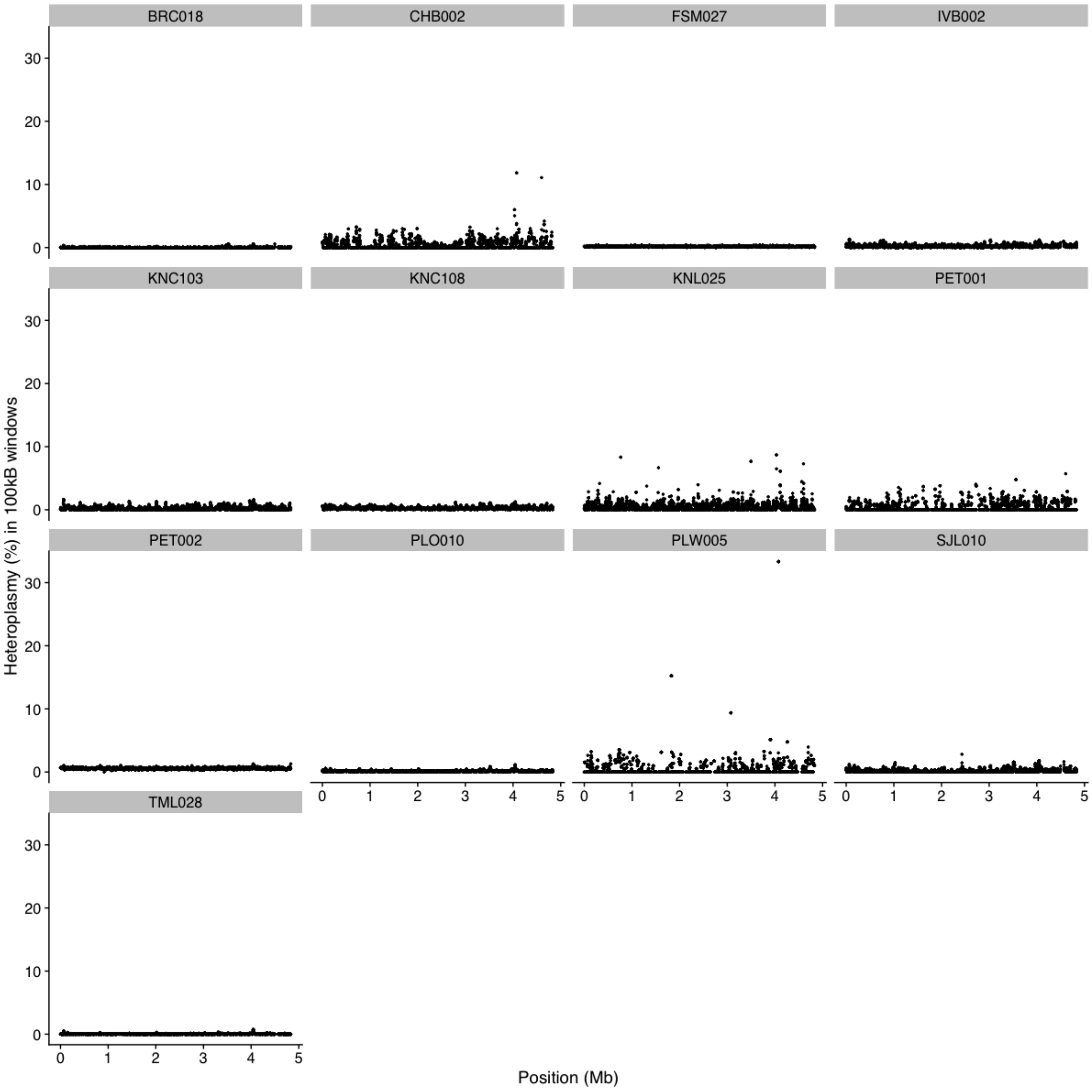

**Figure S6. Estimated heterozygosity in 100kB windows with 10kB step size for samples <3X.** The percentage heterozygosity in 100kB windows along the genome is plotted for each individual.

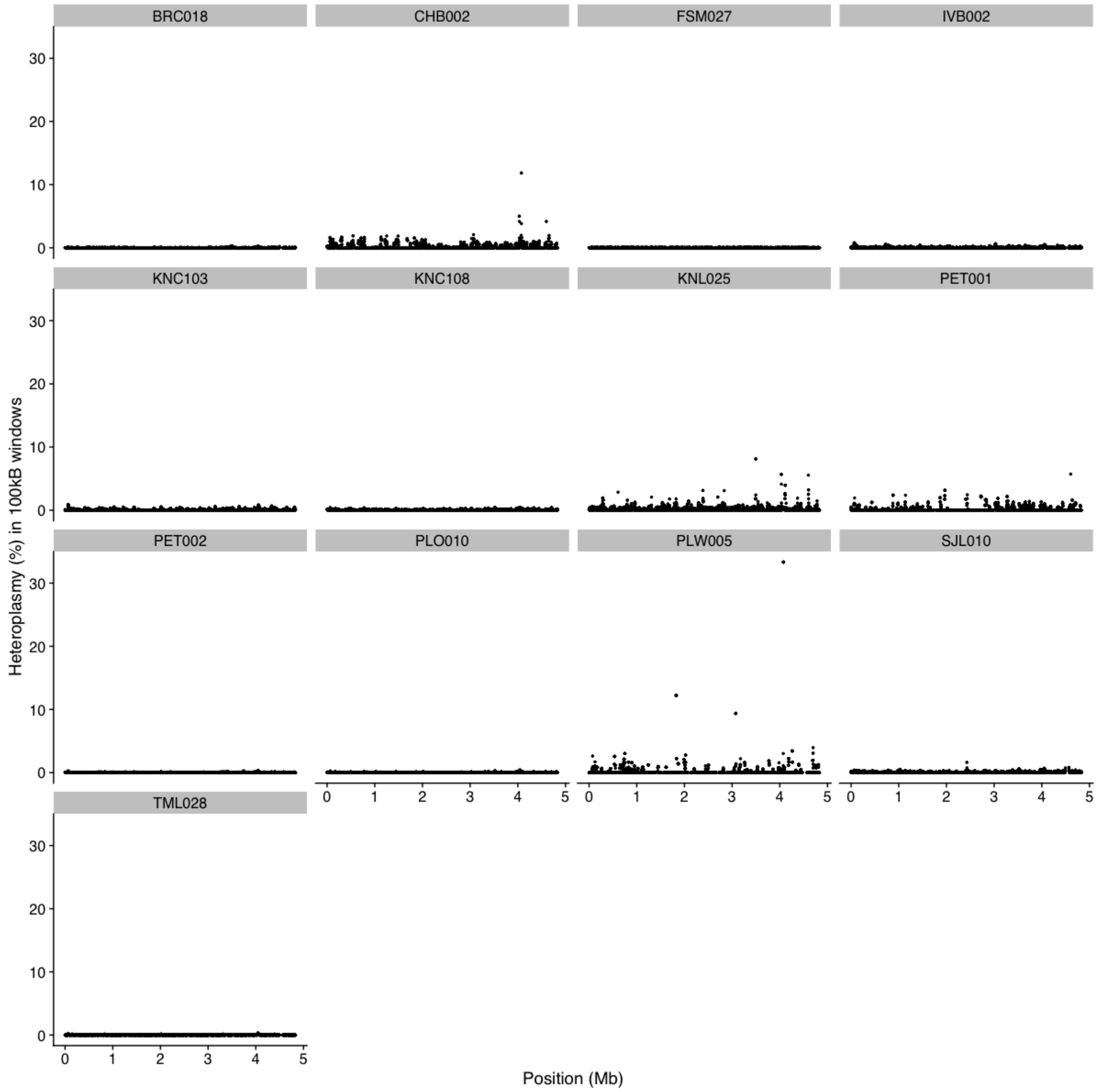

**Figure S7. Estimated heterozygosity in 100kB windows with 10kB step size for samples <3X (without possible post-mortem deamination).**

The percentage heterozygosity in 100kB windows along the genome is plotted for each individual.

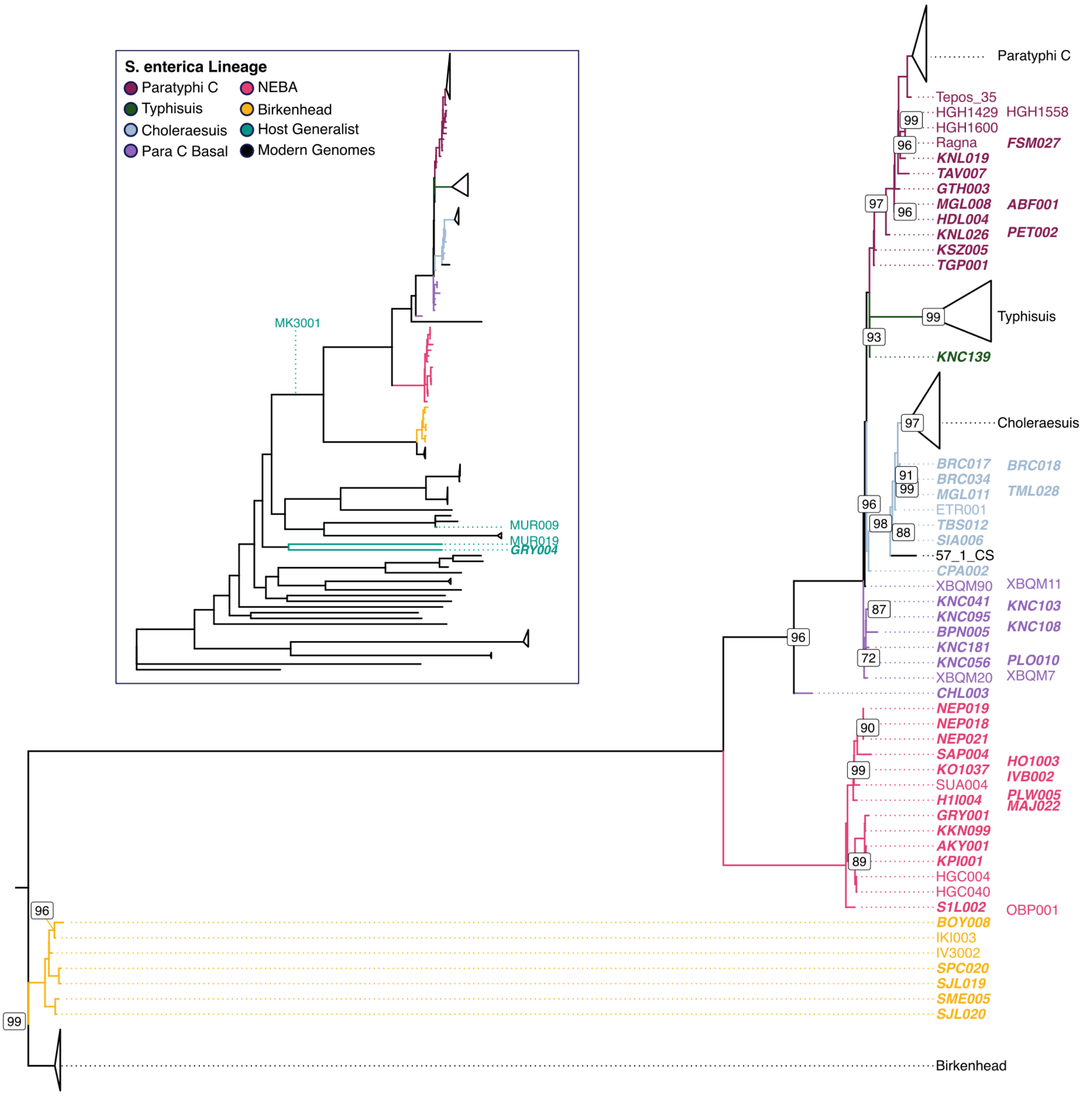

**Figure S8. Placement of ancient Samples in Para C and wider *S. enterica* tree.**

The majority of ancient genomes fall into the Para C lineage. A maximum likelihood tree showing these relationships is represented here; nodes with bootstrap support < 100 are labelled, and samples excluded due to quality issues (i.e. high levels of heteroplasmy or with <50% covered ≥3X) were placed individually, and their approximate placement is depicted. New samples are in a bold italic typeface. The colour represents the Para C sub-lineage that each sample belongs to; these are: Birkenhead (yellow); NEBA (pink); basal to the rest of Para C (purple); Choleraesuis (blue); Typhisuis (dark green); Paratyphi C (maroon). The full *S. enterica* tree is inset to show the placement of non-Para C ancient genomes (in green).

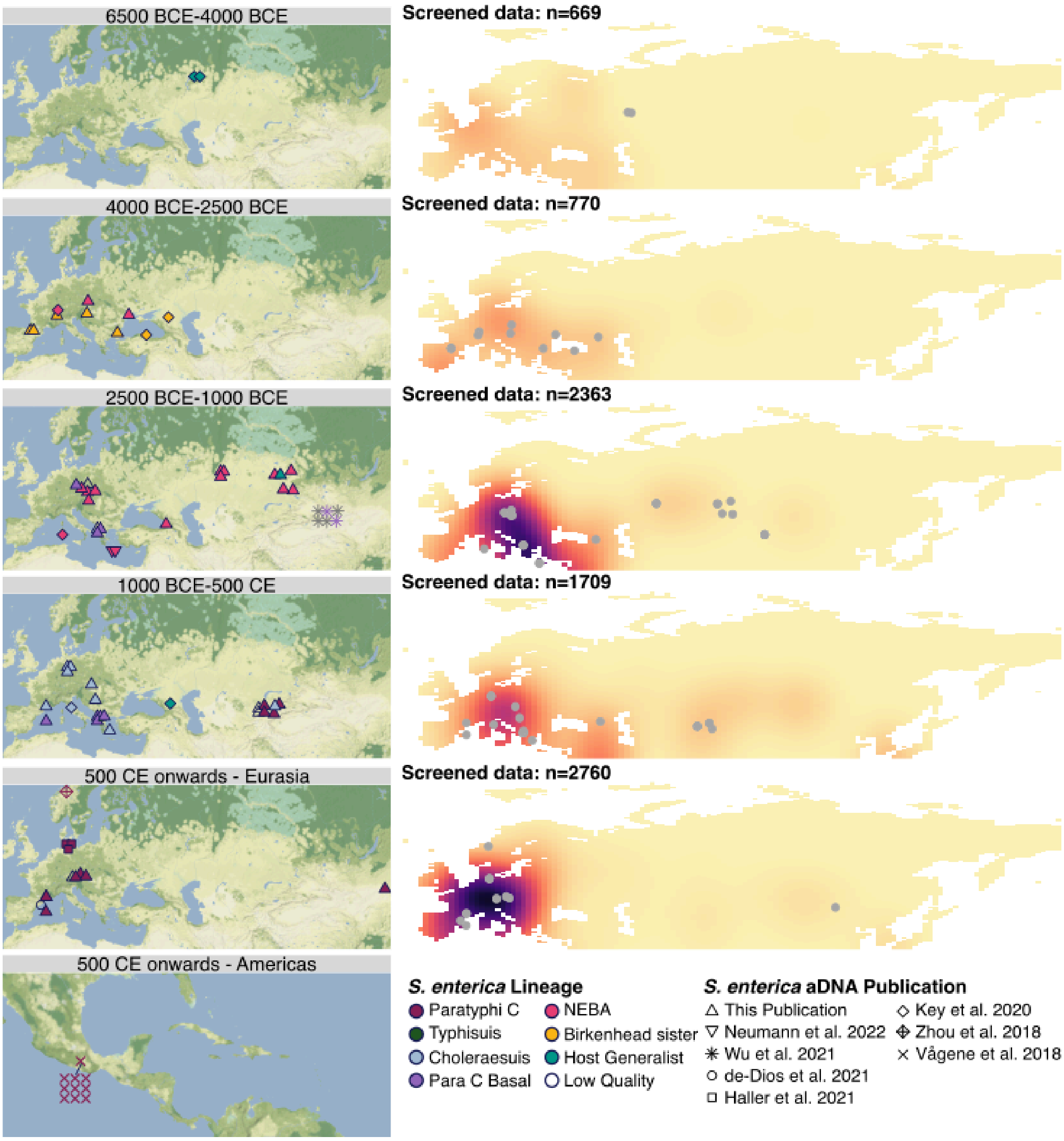

**Figure S9. Non-cranial screening density.**

Density of sampling for ancient pathogen screening from the Max Planck Institute for Evolutionary Anthropology (Leipzig) in 2500 year time windows with *S. enterica* detections marked as grey points. Darker colours indicate greater sampling density; density is normalised across all panels. Total sample numbers for each time period are also marked. The left-hand column shows lineage assignments for each detection. These lineages are: Birkenhead sister lineage (yellow); NEBA (pink); basal to the rest of the Para C lineage (purple); Choleraesuis (blue) Typhisuis (dark green); Paratyphi C (maroon); non-specific host generalists (turquoise).

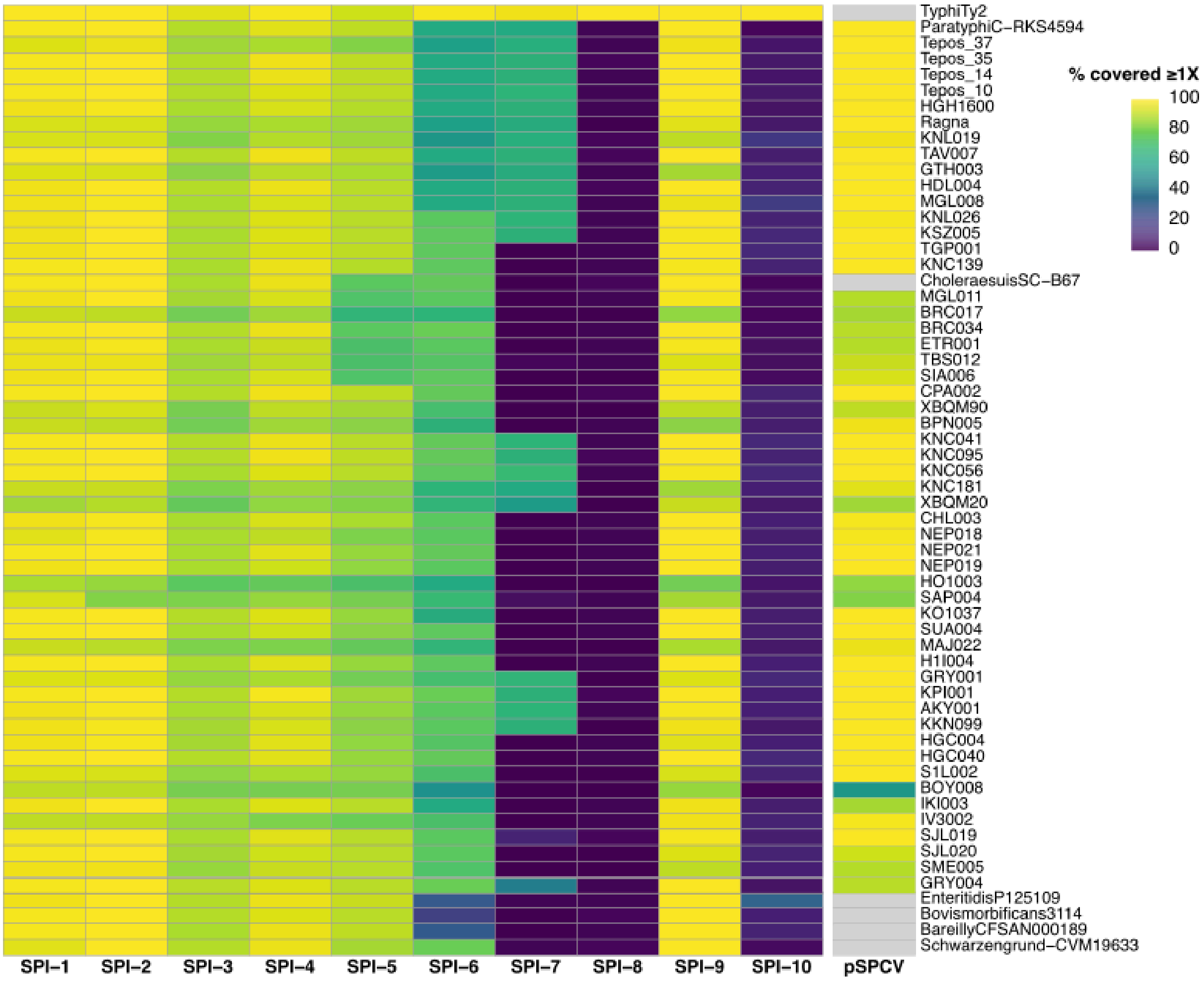

**Figure S10. Presence and absence of Salmonella Pathogenicity Islands (SPI1-10) and Paratyphi C virulence plasmid (pSPCV).**

For each genome ≥3X, the percentage of each pathogenicity island (or virulence plasmid) covered ≥1X is presented in the heatmap. For modern genomes, the presence of the virulence plasmid was not assessed, as data came from chromosomal assemblies.

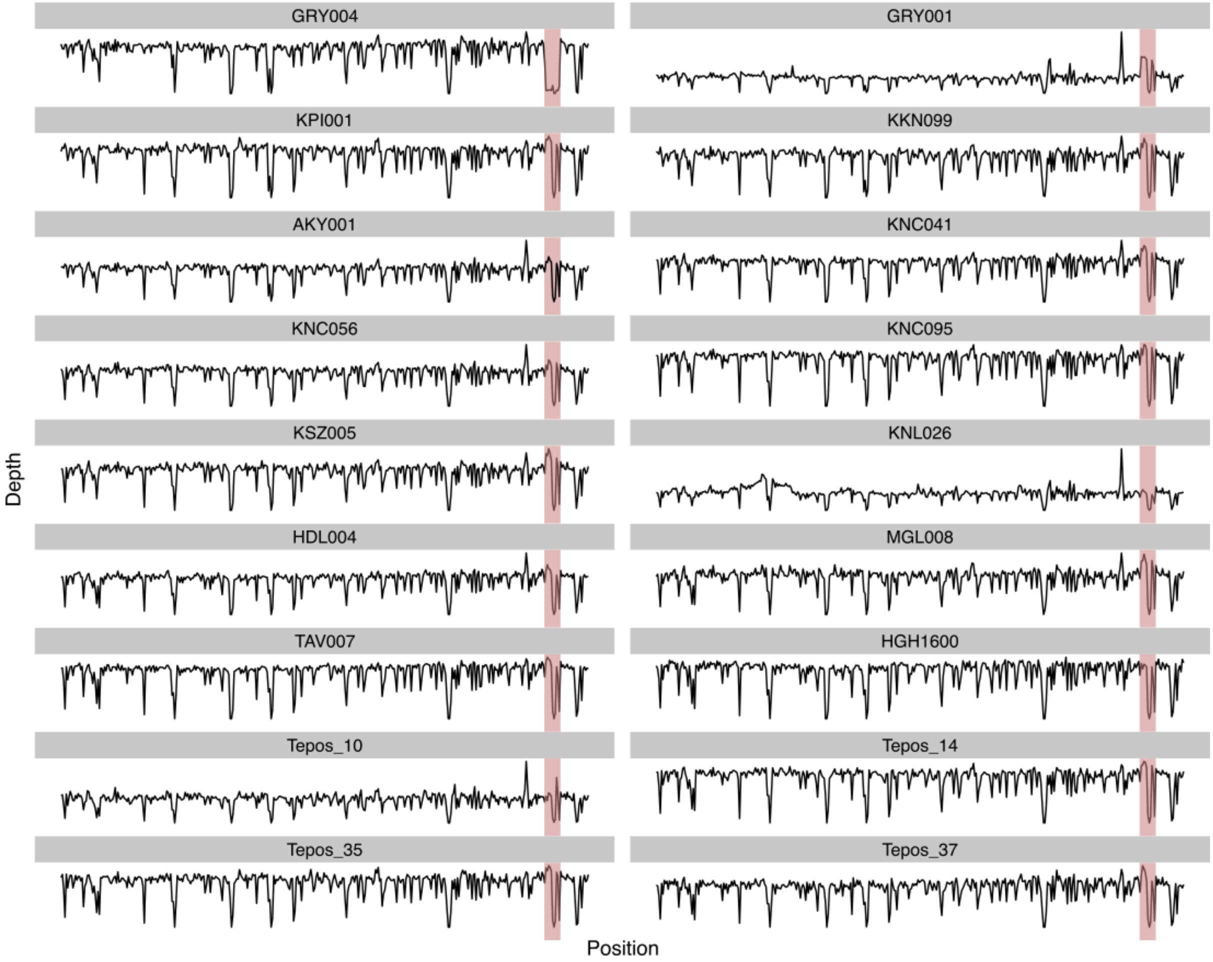

**Figure S11. Comparison between genome-wide coverage with SPI-7 coverage.**

Mean coverage (Depth) across the Typhi CT18 reference genome (calculated using qualimap) is plotted for each SPI-7 positive sample used in analysis, along with GRY001 and GRY004. The SPI-7 region is highlighted.

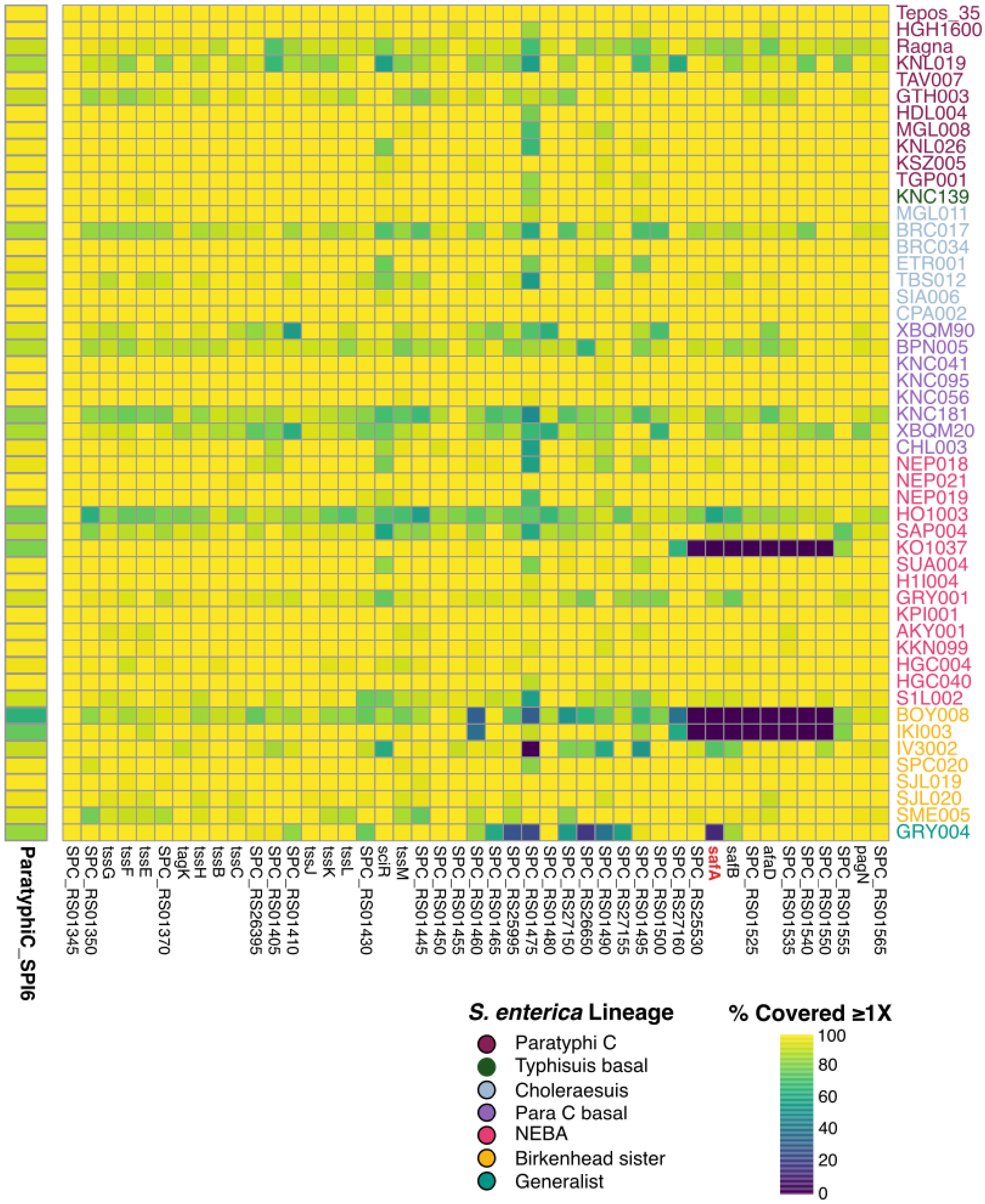

**Figure S12. Assessment of SPI-6 Presence from Paratyphi C Alignment.**

Percentage of the entire SPI-6 region and individual genes covered when ancient data is aligned to the Paratyphi C reference. Lighter colours indicate that a greater proportion of the gene is covered. Ancient cases are coloured by *S. enterica* lineage. The gene *safA* is highlighted in red.

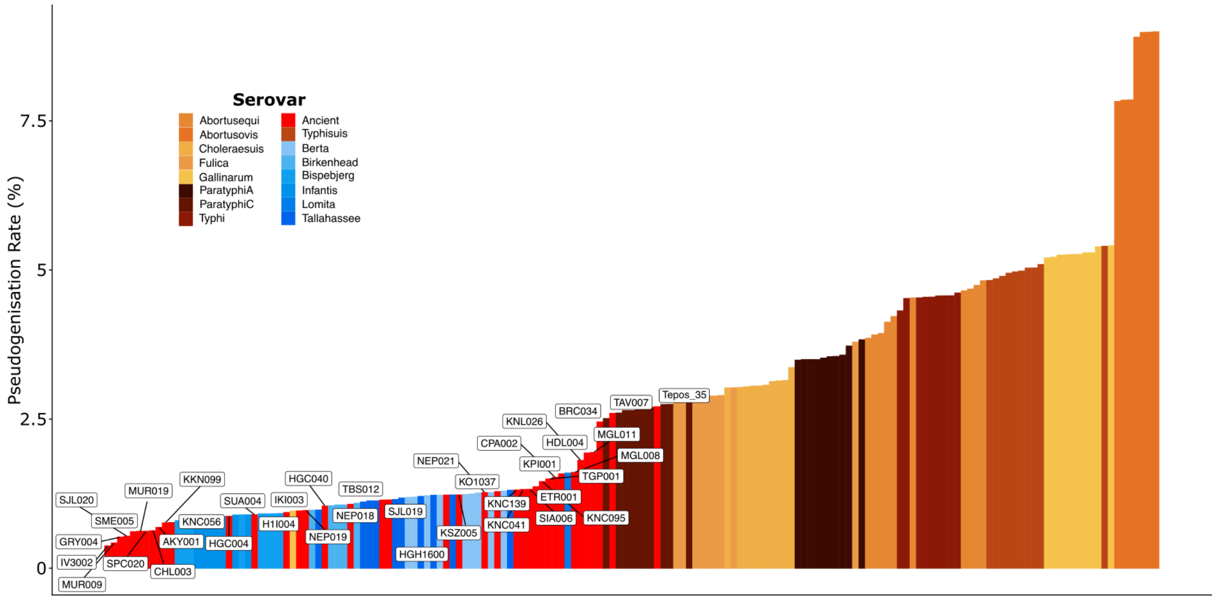

**Figure S13. Pseudogenisation rates are associated with host specificity.**

Pseudogenisation rates in all samples over 5X with a minimum of 3750 genes covered >90% at 1X. Ancient genomes are coloured in red and labelled with sample IDs; host generalist strains are coloured in a blue palette, and host specific strains are coloured in an orange-brown palette.

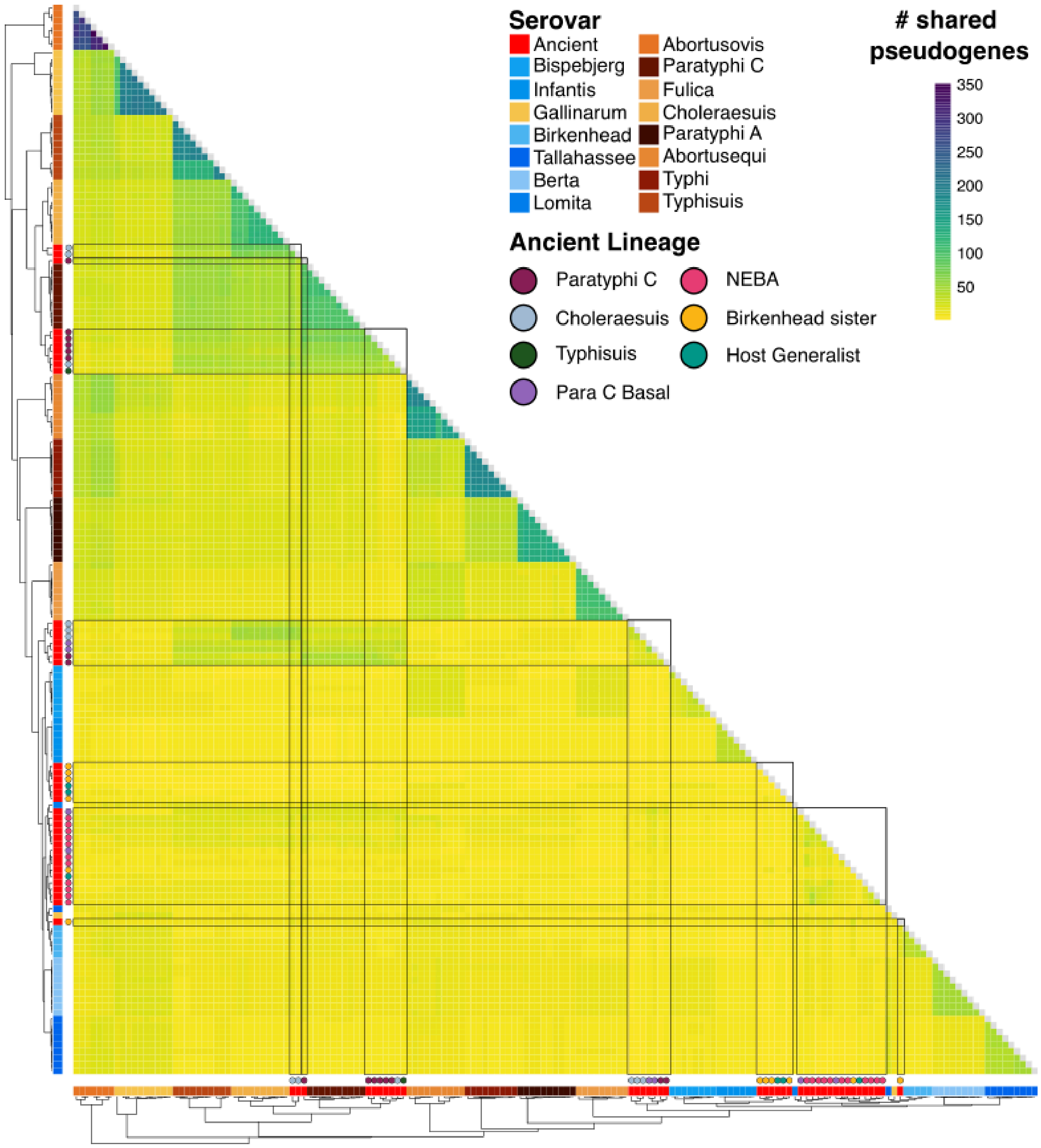

**Figure S14. Heatmap of shared pseudogenes.**

Modern serovars are annotated to the side of the heatmap, and ancient samples are additionally coloured by phylogenetic lineage. Hierarchical clustering was performed on a matrix of shared pseudogene counts, which is plotted on the heatmap. Darker colours represent a greater number of shared pseudogenes. Plotted with pheatmap package.

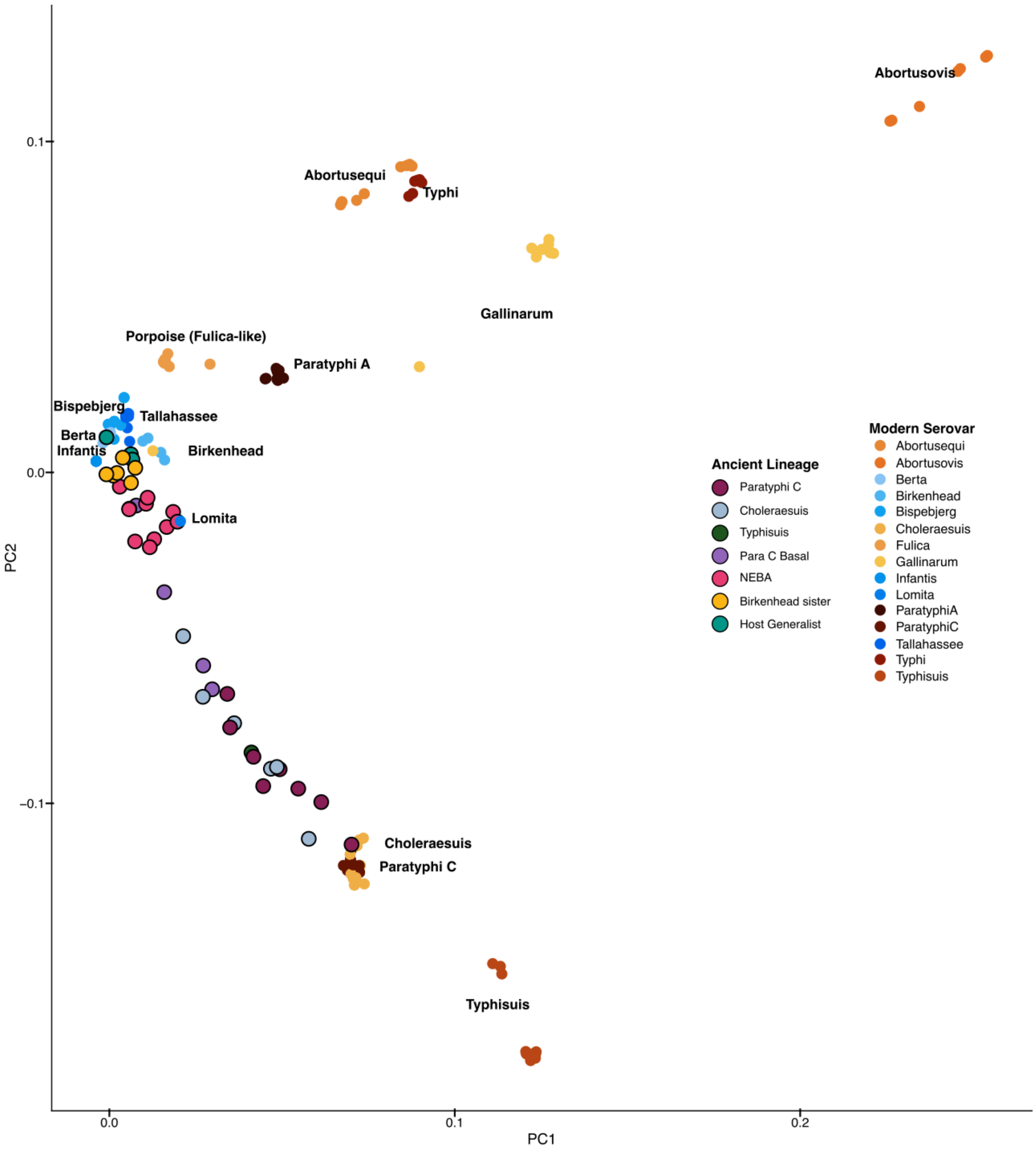

**Figure S15. Principal component analysis of shared pseudogene counts.**

Principal component analysis was performed on the shared pseudogene count matrix. Individual points represent individual genomes, and are coloured by either modern serovar (smaller points) or ancient lineage (larger points, outlined in black). There is a gradient of ancient samples between modern host-specific lineages (largely represented by older, likely host-generalist genomes) to modern Choleraesuis and Paratyphi C genomes. Plotted using ggplot2.

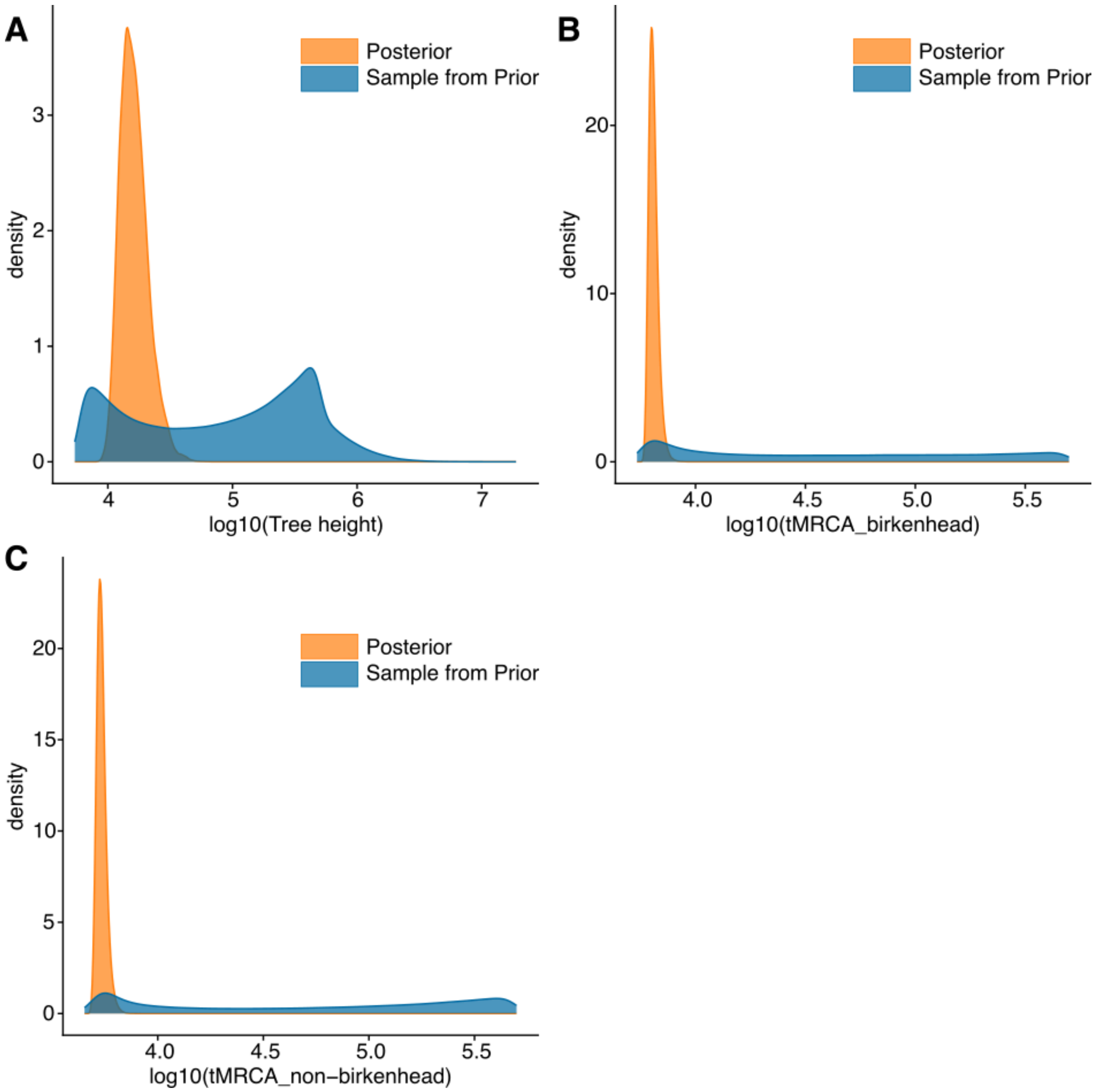

**Figure S16. Assessment of impact of priors on TMRCA inference.**

Density of estimated TMRCAs for **A.** the full Para C tree **B.** the Birkenhead lineage only **C.** All non-Birkenhead genomes**.** Posterior distributions are coloured orange, estimates from sampling from the prior are coloured blue. The x-axis is on a $\log_{10}$ scale, in order to be able to visually compare the two distributions.

41. Милошевић, П. *Археологија и историја Сирмијума*. (Матица Српска, Нови Сад,

2001).

49. Favre, S. & Mottet, M. *Le Site Préhistorique Du Petit-Chasseur (Sion, Valais) : 9. Dolmens M 12 et M 13. Approche Des Différents Niveaux Préhistoriques*. (Cahiers d'Archéologie Romande, Lausanne, 2011).

50. Bocksberger, O.-J. *Le Site prehistorique du Petit-Chasseur, Sion, Valais: Catalogue et planches*. 151 (Bibliothèque historique vaudoise, 1976).

51. Gallay, A. *Autour du Petit-Chasseur: l'archéologie aux sources du Rhône, 1941-2011*. (Editions

Errance, 2011).

52. Gallay, A. Chefs de guerre et matrones: Stèles et statues-menhirs: Chefs de guerre et matrones. *L'Archéologue: revue d'archéologie et d'histoire* **104**, 12–13 (2009).

53. Fenner, J. N., Tumen, D. & Khatanbaatar, D. Food fit for a Khan: stable isotope analysis of the elite Mongol Empire cemetery at Tavan Tolgoi, Mongolia. *J. Archaeol. Sci.* **46**, 231–244 (2014).

54. Youn, M. *et al.* Dating the Tavan Tolgoi site, Mongolia: Burials of the nobility from Genghis Khan's era. *Radiocarbon* **49**, 685–691 (2007).

55. Jakab, J. Tesárske Mlyňany, časť Tesáre nad Žitavou Poloha Gočol. in *Výskumná správa 18134/13. Dokumentácia AÚ SAV, v. v. i.* (Nitra, 2013).

63. Li, H. & Durbin, R. Fast and accurate short read alignment with Burrows-Wheeler transform.

*Bioinformatics* **25**, 1754–1760 (2009).

64. Picard. http://broadinstitute.github.io/picard/.

87. Cassidy, L. M. *filter_bam.py*. github.com/LCLabTCD/aDNA_data_processing/ (2023).

88. Quinlan, A. R. BEDTools: The Swiss-Army Tool for Genome Feature Analysis. *Curr. Protoc. Bioinformatics* **47**, 11.12.1–34 (2014).

89. Danecek, P. *et al.* Twelve years of SAMtools and BCFtools. *Gigascience* **10**, giab008 (2021).
90. Cingolani, P. *et al.* A program for annotating and predicting the effects of single nucleotide polymorphisms, SnpEff: SNPs in the genome of Drosophila melanogaster strain w1118; iso-2; iso-3. *Fly (Austin)* **6**, 80–92 (2012).
91. R Core Team. R: A Language and Environment for Statistical Computing. Preprint at https://www.R-project.org/ (2023).
92. Klopfenstein, D. V. *et al.* GOATOOLS: A Python library for Gene Ontology analyses. *Sci. Rep.* **8**, (2018).
93. Jones, P. *et al.* InterProScan 5: genome-scale protein function classification. *Bioinformatics* **30**, 1236–1240 (2014).
94. Edwards, D. J., Duchene, S., Pope, B. & Holt, K. E. SNPPar: identifying convergent evolution and other homoplasies from microbial whole-genome alignments. *Microb. Genom.* **7**, (2021).
95. Murray, G. G. R. *et al.* Genome reduction is associated with bacterial pathogenicity across different scales of temporal and ecological divergence. *Mol. Biol. Evol.* **38**, 1570–1579 (2021).
96. Wickham, H. *et al.* Welcome to the tidyverse. *Journal of Open Source Software* vol. 4 1686 Preprint at https://doi.org/10.21105/joss.01686 (2019).
97. Barrett, T. *et al.* data.table: Extension of `data.frame`. Preprint at https://r-datatable.com (2025).
98. Kolde, R. pheatmap: Pretty Heatmaps. Preprint at https://github.com/raivokolde/pheatmap (2025).
99. Jackson, I., Woodman, P., Dowd, M., Fibiger, L. & Cassidy, L. M. Ancient genomes from bronze age remains reveal deep diversity and recent adaptive episodes for human oral pathobionts. *Mol. Biol. Evol.* **41**, msae017 (2024).
100. Larkin, M. A. *et al.* Clustal W and Clustal X version 2.0. *Bioinformatics* **23**, 2947–2948 (2007).
101. Gouy, M., Tannier, E., Comte, N. & Parsons, D. P. Seaview version 5: A multiplatform software for multiple sequence alignment, molecular phylogenetic analyses, and tree reconciliation.

*Methods Mol. Biol.* **2231**, 241–260 (2021).

112. Spurbeck, R. R., Tarrien, R. J. & Mobley, H. L. T. Enzymatically active and inactive phosphodiesterases and diguanylate cyclases are involved in regulation of motility or sessility in

Escherichia coli CFT073. *MBio* **3**, (2012).

123. Vollmer, W., Blanot, D. & de Pedro, M. A. Peptidoglycan structure and architecture. *FEMS*

*Microbiol. Rev.* **32**, 149–167 (2008).

133. Cordero-Alba, M. & Ramos-Morales, F. Patterns of expression and translocation of the ubiquitin

ligase SlrP in Salmonella enterica serovar Typhimurium. *J. Bacteriol.* **196**, 3912–3922 (2014).
